# Mechanistic quantification of the effects of genes, environment, and development on human behaviour

**DOI:** 10.1101/2025.03.17.643651

**Authors:** Mauricio González-Forero, Aida Gómez-Robles

## Abstract

A long-standing question is to what extent behaviour is determined by genes, environment, or else. Common practice estimates genetic components using heritability, which is correlational, population-level, and variously problematic to measure genetic contribution. Ideally, contributions would be quantified via experimental intervention, which is often infeasible for humans and other organisms, but possible *in silico*. We formulate mathematical methods to quantify contributions to phenotypic differences *in silico*, and apply them to an *in-silico* replica of human skill development and evolution. We find that, in this replica, adult skill level is more reactive to differences in developmental history than to genetic, environmental, or social differences from *Homo habilis* to *H. sapiens*. This occurs although, in this replica, adult skill level has evolved an increased reactivity to environmental and social differences, and a decreased reactivity to genetic differences, which is consistent with observation. Our results predict a window of sensitivity at the end of brain growth. They also predict that adverse genetic and environmental conditions that would developmentally yield low adult skill level can be compensated by time-specific environmental changes or slight changes in developmental history, potentially yielding outstanding adult skill level. Our analyses suggest a major capability of developmental history to influence adult human skill level.

## Introduction

Adult human behaviour is the product of a complex interplay of genetic, developmental, environmental, and social factors (Anreiter, Sokolowski, & Sokolowski, 2018; Boyce, Sokolowski, & Robinson, 2020; Johnson, 2007). The brain expresses thousands of genes (Zeng et al., 2012) and extensive research efforts seek to identify genes contributing to behaviour (Canela-Xandri, Rawlik, & Tenesa, 2018; Sakaue et al., 2021). Moreover, behaviour is affected by developmental feedbacks between genes, developmentally earlier behaviour, and environment, such that these feedbacks have been suggested to explain the otherwise puzzling increases in mean IQ over the past century (Dickens & Flynn, 2001). Additionally, behaviour is affected by developmental noise as evidenced by studies of essentially genetically identical individuals growing in the same environment who develop different behaviours (Freund et al., 2013; Mitchell, 2018; Turkheimer & Waldron, 2000). By some estimates, up to 50% of variation in human cognitive abilities may stem from developmental noise (Plomin & Kawakami, 2025). Furthermore, multiple lines of evidence point to the existence of human specialisations for neuroplasticity (Drennan, Sherwood, & Gómez-Robles, 2026; K. R. Rosenberg, 2021). The high level of plasticity of the human brain helps explain human learning abilities (van Schaik & Burkart, 2011) and makes it possible to develop intervention strategies that can change behaviours over life (Sparling, Dragomir, Ramey, & Florescu, 2005). Observed differences between chimpanzees and humans indicate that increased neuroplasticity evolved in humans after both lineages diverged, together with an overall increase in brain size and a general reorganization of the brain (Drennan et al., 2026).

Despite the recognition of these multiple factors, disentangling the causes of variation in human behaviour remains challenging. Various approaches have been devised to assess causality, but difficulties persist given the commonly observational nature of human data (Kievit et al., 2018; Ota et al., 2025; M. D. Rosenberg, Casey, & Holmes, 2018; Turkheimer & Waldron, 2000). For instance, a common but problematic way to quantify the causes of variation is via heritability: a trait with higher heritability is often interpreted at face value as having a stronger genetic component and being less plastic (Devlin, Fienberg, Resnick, & Roeder, 1997; Herrnstein & Murray, 1994; Jensen, 1972). However, heritability is problematic to assess genetic causality as it is correlational, depends on population-level phenomena such as genetic, phenotypic, and environmental variation, and does not quantify the causes of a phenotype for a given individual (Feldman & Lewontin, 1975). Moreover, unlike brain size and reorganization, which can be inferred to some extent from endocranial size and shape (Coqueugniot, Hublin, Veillon, Houët, & Jacob, 2004; Gunz et al., 2020; Gunz, Neubauer, Maureille, & Hublin, 2010), quantifying the relative influence of genetic and environmental factors on behaviour from fossils remains out of reach.

Ideally, quantifying the causes of phenotypic variation at the individual-level would be done by determining the effects of experimental intervention, which is a gold-standard to establish causality (Pearl, 2009). For many human behaviours, such interventions would involve infeasible actions such as altering the genetic and environmental make-up of an individual’s past. Although such interventions are often impossible with real individuals, they are possible *in silico*. Many mathematical or computational models exist that mechanistically describe the development of phenotypes (e.g., mammalian teeth, Salazar-Ciudad & Jernvall, 2010; vertebrate digits, Sheth et al., 2012; and human skill level, Koster et al., 2020). By analysing experimental *in-silico* interventions in these models, it is possible to quantify the causes of variation in the developed *in-silico* phenotypes, yielding an inference of the causes of variation in the observed phenotypes in real organisms.

In this paper, we use, develop, and apply four methods to quantify the causes of individual-level phenotypic variation when a model of development is available. First, we use the *developmental sensitivity* of a phenotype (González-Forero, 2024b), which quantifies the total effect of marginally changing a variable at a particular point earlier in development on another variable later in development. Second, we use the *developmental elasticity* of a phenotype (Caswell, 2019), which quantifies the proportional total effect of a variable on another variable. Third, we define *developmental reactivity*, which quantifies the proportional total effect of a set of variables on another variable relative to all variables affecting development. Fourth, we define *first-order contributions*, which quantify the mechanistic (not correlational) genetic and environmental contributions to the differences between the phenotypes of individuals developed under different conditions.

We apply these methods to quantify how much genetic, environmental, developmental, and social factors causally affect the development of different adult human skill levels in a model of human brain size development and evolution (González-Forero, 2024a; González-Forero, Faulwasser, & Lehmann, 2017; González-Forero & Gardner, 2018). We refer to this model as “brain model”, given its original emphasis on modelling brain size evolution. In this model, the brain enables a cognitive ability termed “skill”, defined as an ability to perform some behaviour(s), where some of the brain’s energy consumption is for increasing and maintaining this ability, that is, for “learning” and “memory” (Methods). We use this model because it replicates major patterns of human development and evolution, thereby describing mechanistically and quantitatively a wide array of observed patterns that are often viewed as characteristic of humans. This ability of the model suggests that it may already capture some of the important causes of human phenotypic variation. However, our conclusions should only be interpreted as preliminary since more conclusive statements would require that many models, where assumptions are modified, are competed to assess which model best explains the data (Cranmer, Brehmer, & Louppe, 2020). For further discussion of the model limitations, see Supplementary Information (SI) section S1.

Some of the major aspects of human development and evolution that the brain model recovers are the following (Fig. 1; González-Forero, 2024a; González-Forero et al., 2017; González-Forero & Gardner, 2018; González-Forero & Gómez-Robles, 2025). The model recovers the expansion of adult brain and body sizes from *Homo habilis* (empirical data from McHenry & Coffing, 2000) to *H. sapiens* scale (empirical data from autopsy hospital reports from Washington, DC, and Bethesda, MD collected between 1964 and 1973; Dekaban & Sadowsky, 1978). Simultaneously, the model recovers the evolution of childhood, adolescence, and adulthood periods with timing matching that observed in *H. sapiens* (empirical data reviewed in Gluckman & Hanson, 2006). Moreover, this model identifies a transition to seemingly cumulative culture in the Middle Pleistocene (González-Forero & Gardner, 2018; González-Forero & Gómez-Robles, 2025), which is consistent with the archaeological record (Barham et al., 2023; Paige & Perreault, 2024; Roebroeks & Villa, 2011).

**Figure 1:**
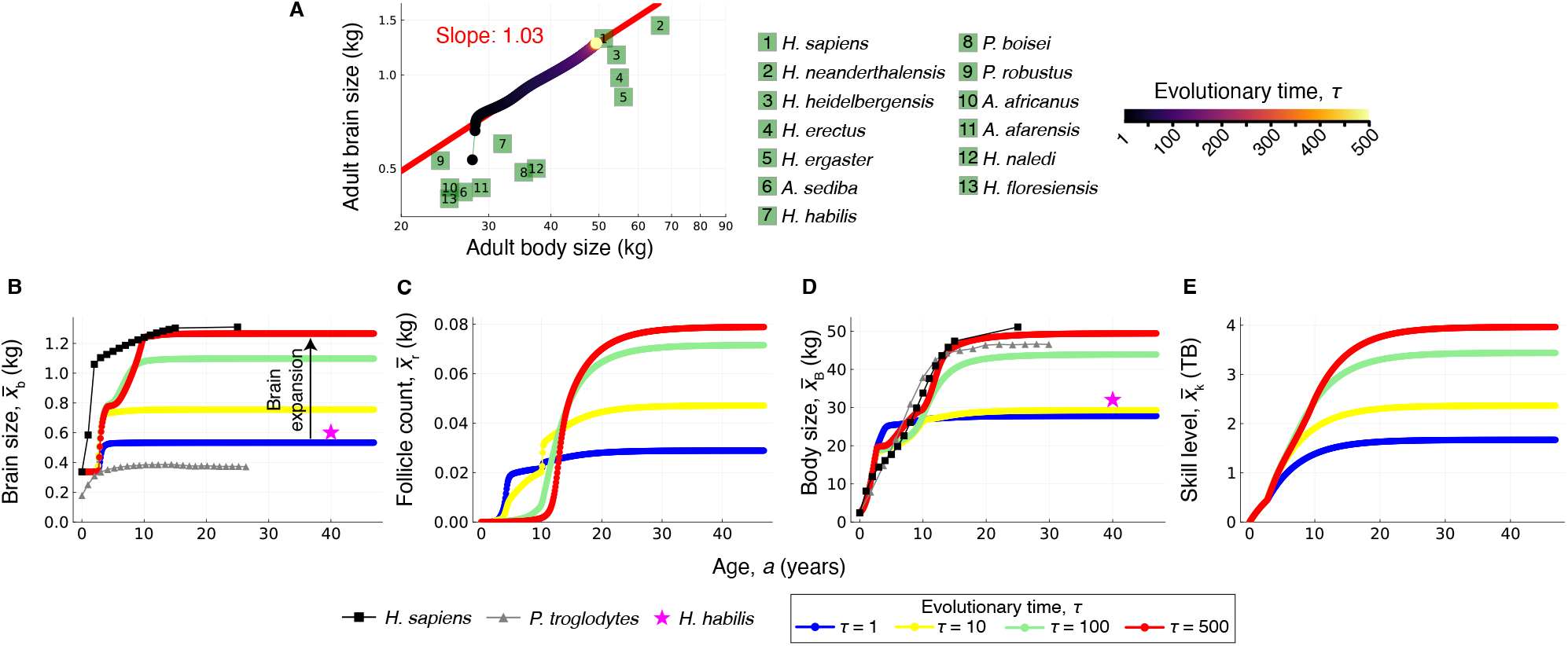
*In-silico* replica of the evolution and development of hominin brain size, body size, and skill level as obtained by the brain model. (**A**) Green squares show the empirically observed mean adult brain and body sizes in log-log scale for 13 hominin species (data for females when possible). Coloured dots give the recovered evolution of adult brain and body sizes by the brain model, shifting from those of *Homo habilis* to those of *H. sapiens*. (**B-E**) Developmental dynamics underlying the trajectory in **A**: for (**B**) brain size, (**C**) mature ovarian follicle count (in mass units), (**D**) body size, and (**E**) skill level (in information units, terabytes or TB). Panels **B**,**D** also show empirically observed cross-sectional mean values of *H. sapiens* females (black squares), *Pan troglodytes* females (gray triangles), and adult *H. habilis* of mixed sexes for brain size and of females for body size (pink stars). The ages at which tissues start or stop growing, the tissue sizes that develop, or the preadolescent growth spurt in body size are all emergent features of the model rather than being predefined parameters. One evolutionary time step is the time from mutation to fixation, which would be 11.5 kyr assuming fixation takes 500 generations and generation time is 23 years. All panels are taken from González-Forero (2024a), except that a typo in the computer code of González-Forero (2024a) has been corrected as described in SI section S6. This correction improves the ontogenetic fit to observed data relative to González-Forero (2024a), and matches the equilibrium results of González-Forero and Gardner (2018). See González-Forero (2024a) for data sources.

The model also mechanistically recovers a linear relationship between skill level (or cognitive ability) and brain size, which is consistent with observation. Specifically, the model finds that, if skill level and brain size plateau over life, adult skill level and adult brain size are proportional (eqs. 13 and 14 of González-Forero et al., 2017, briefly re-derived in SI section S.3.1.4): that is, adult skill level = proportionality factor × adult brain size, where the proportionality factor increases with the metabolic costs of brain maintenance, decreases with the metabolic costs of memory, and increases with the fraction of the brain metabolic rate allocated to the skill considered. This proportionality is not an assumption but a result of the model’s energy conservation analysis, involving fewer assumptions than those made by the whole model and so being relatively general. The proportionality is consistent with brain size and cognitive abilities being positively correlated in primates (Deaner, Isler, Burkart, & van Schaik, 2007; Heldstab et al., 2016) and other vertebrates (MacLean et al., 2014). It also agrees with the consistent yet modest positive association between brain size and cognitive ability in humans (Cox, Ritchie, Fawns-Ritchie, Tucker-Drob, & Deary, 2019; Nave, Jung, Linnér, Kable, & Koellinger, 2019). This proportionality predicts that some of the variation in the empirically observed relationship between brain size and cognitive ability could be due to variation in the metabolic costs of memory and of brain maintenance and in the fraction of brain metabolic rate allocated to the cognitive ability evaluated.

Here we use this model to causally quantify in *in-silico* individuals: 1) the responses of adult skill level to differences experienced earlier in life, 2) how these responses have evolved from *H. habilis* to *H. sapiens*, and 3) the genetic and environmental contributions to differences in adult skill level. This analysis finds relatively small genetic and large developmental effects on adult skill level, such that slightly intervening developmental history can compensate for genetic and environmental adversity. The analysis also finds a window of sensitivity of adult skill to perturbations at the age at which brain growth ends.

## Methods

Here we describe the brain model and our four methods to quantify the causes of individual phenotypic differences.

### Brain model overview

The brain model can be seen as having three major components: development, selection, and evolution.

#### Development

The development component mechanistically describes the construction of each individual’s phenotypes over life using dynamic equations based on energy conservation. The development component (Fig. 2A) may be subdivided into two parts: a metabolic part and a behavioural part. The metabolic part specifies the growth rates of tissues and their function based on energy conservation. The behavioural part specifies the brain’s function.

**Figure 2:**
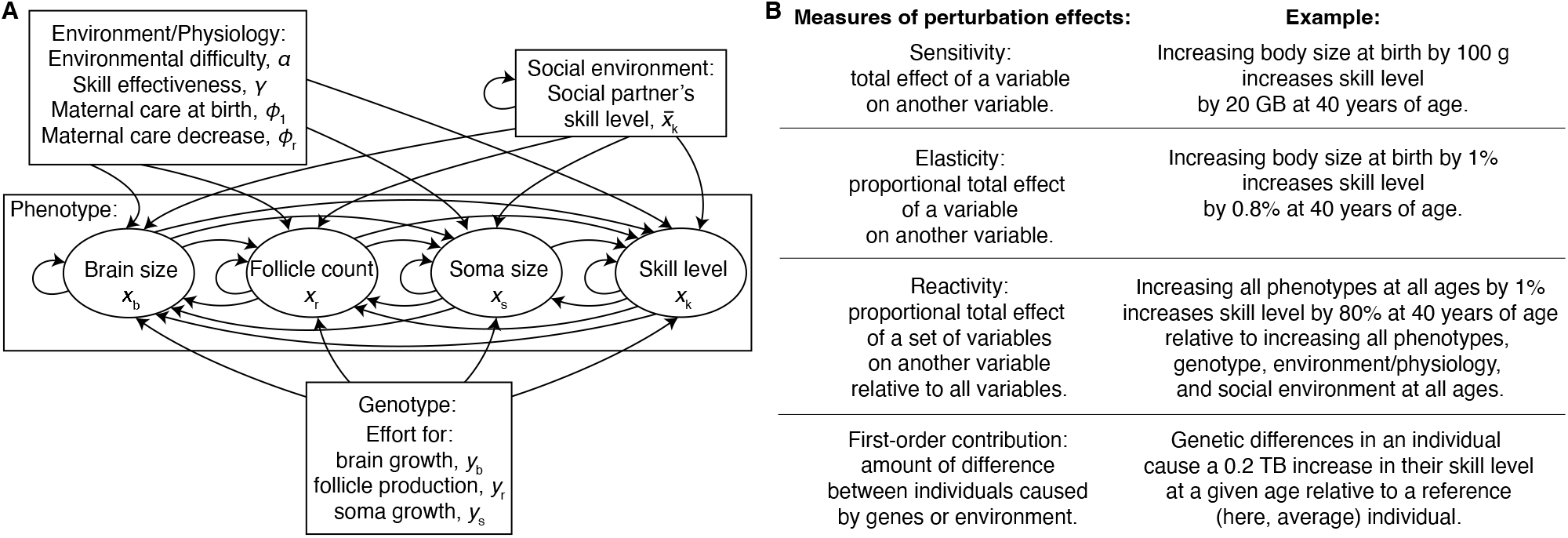
Causal diagram of development in the brain model and measures of the effects of perturbation. (**A**) Causal diagram of development in the model, where an arrow indicates a direct causal effect of a variable on another one or on itself at the next age. Variables are classified into genotypic, phenotypic, environmental/physiological, and social environmental. Genotypic variables are controlled by the individual’s genotype. Phenotypic variables are constructed over development; the developmental history for a given age is defined here as the phenotypic variables at earlier ages. Environmental or physiological variables affect the development of phenotypic variables. Social environmental variables are the skill level (and so are phenotypic variables) of social partners, which affect the development of the focal individual. (**B**) We quantify the effect of perturbation, that is, of altering a variable at a given age, on adult skill level, using four measures: sensitivity d*x*/d*y*, which measures total effects, that is, the effect of altering a variable over all causal pathways linking it to the response variable; elasticity (*y* /*x*)d*x*/d*y*, which measures the proportional total effects; reactivity, which normalizes elasticity over all the possible interventions over development; and first-order contributions, which quantify the mechanistic contribution to phenotypic differences of genes and environment.

#### Metabolic part

The metabolic part partitions the body of an individual into three tissues: brain, reproductive, and the remainder referred to as somatic tissue. This body partitioning was considered to model brain size evolution with a simple description of fitness, such that fertility depends on reproductive tissue size. The model can consider other body partitions (e.g., brain regions or digestive system). The metabolic part describes the growth rates of tissue sizes based on energy conservation. Specifically, following West, Brown, and Enquist (2001), the model assumes that a tissue’s energy consumption equals the tissue’s energy consumption on growth and maintenance, yielding an equation describing the tissue’s growth rate in terms of the fraction of body growth energy allocated to the tissue. This specifies a classic life history tradeoff between energy allocation to the growth of the different tissues. The metabolic part also describes the individual’s skill learning rate based on energy conservation. The metabolic part does not specify what the skills are nor their function, so in that sense skills can be interpreted quite generally. An equation describing this learning rate is obtained by assuming that some of the brain energetic consumption is due to learning and memory of these skills. This learning rate equation yields the proportionality between adult brain size and adult skill level (Eq. S8).

In symbols, we denote an individual’s brain mass, mass of reproductive tissue, mass of remaining somatic tissue, and skill level, each at age *a*, respectively by *x*_b*a*_, *x*_r*a*_, *x*_s*a*_, and *x*_k*a*_. The vector of the individual’s phenotypes at age *a* is **x**_*a*_ = (*x*_b*a*_, *x*_r*a*_, *x*_s*a*_, *x*_k*a*_)^⊺^. The development component describes the phenotypes **x**_*a*+1_ of an individual at age *a* + 1 as a function of the phenotypes at the previous age, of “genotypic traits” **y**_*a*_ = (*y*_b*a*_, *y*_r*a*_, *y*_s*a*_)^⊺^ that are under direct genetic control and that specify the energy allocated to the growth of each tissue at each age, parameter values ***ϵ***_*a*_ including empirically estimated brain metabolic costs, and the skill level of an average social partner 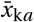 due to cooperation or competition:

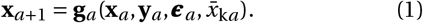

The function **g**_*a*_ is the developmental map, which is nonlinear. The description of development by means of dynamic equations modulated by genetically controlled traits, as we have done, has long roots (e.g., Alberch, Gould, Oster, & Wake, 1979; Rice, 1997). We sometimes denote the individual’s phenotype and genotype at age *a* by **z**_*a*_ = (**x**_*a*_; **y**_*a*_), where the semicolon denotes a linebreak so **z**_*a*_ is a vector. The vector of the phenotypes over the life course is 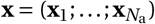, and similarly for **y** and **z**. An overbar as in 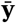 denotes population average. For further details, see SI sections S2-S3.

The metabolic part enables the model to be parameterised with empirical data of brain and skill metabolic costs. With this empirical parameterisation, the model makes quantitative predictions of brain size and skill level in the units given by the empirically estimated metabolic costs. For instance, empirical estimates of metabolic costs of brain maintenance in units of energy/(mass × time) such as joules/(kilograms × years) lead to predictions of brain size in kilograms at a given age in years. Similarly, empirical estimates of metabolic costs of skill maintenance (memory) in units such as joules/(skill unit × years) lead to predictions of skill level in the skill unit at a given age in years. Although empirical estimates of the metabolic costs of memory of human skills appear to be unavailable, there exist empirical estimates of the metabolic cost of information storage (in bytes) in synapses in Purkinje cells in rats (Brunel, Hakim, Isope, Nadal, & Barbour, 2004; Howarth, Peppiatt-Wildman, & Attwell, 2010; Steuber et al., 2007). The model has used these estimates as relevant benchmarks for the metabolic cost of memory, which entails that the model predicts skills in information units (bytes). Future empirical estimates of the metabolic costs of memory and learning of specific skills would be helpful to improve the model accuracy and measure skills in other units.

#### Behavioural part

The behavioural part of the model specifies the skills and their function. The developmental map **g**_*a*_ depends on the individual’s energy budget that in turn depends on the individual’s efficiency of extracting energy from the environment. The model assumes that some skills enable “energy extraction”, where these skills are defined as the capacity of an individual to overcome ecological or social challenges to extract energy from the environment. Ecological challenges refer to the individual having to obtain energy by using their own skills to solve problems posed by the environment excluding social partners (me vs nature). Social challenges can be either cooperative, between-individual competitive, or between-group competitive. For simplicity and as a first approximation, the model assumes that these social interactions are only between pairs of peers (i.e., individuals of the same age). In cooperative challenges, an individual must use their skills with those of a peer to obtain energy (us vs nature). In between-individual competitive challenges, an individual must use their skills against those of a peer to obtain energy (me vs you). In between-group competitive challenges, an individual must use their skills with those of a peer to compete against the skills of another pair of peers (us vs them). The energy extracted can be used for tissue growth and maintenance. The model implements maternal care phenomenologically rather than mechanistically, where the individual’s skills only become necessary for energy extraction as the individual’s age advances.

The model so far considers the effect of culture or social learning implicitly, via the shape of the function relating skills to the energy extraction efficiency: the function may have strongly or weakly diminishing returns of increasing skill. Strongly diminishing returns could in principle arise under “non-cumulative” cultural settings where individuals cannot increase their energy extraction efficiency substantially after achieving a high skill level; in contrast, weakly diminishing returns could in principle arise under cumulative cultural settings where individuals can increase their energy extraction efficiency despite already being highly skilled, if they can continue to learn from accumulated knowledge in the population. The model has thus not yet considered a vast array of complexities that are likely to be key to skill development, such as explicit social learning dynamics or social interactions between individuals of different ages. However, the model offers a stepping stone towards further realism.

#### Selection

The selection component of the model specifies fitness and so what selection acts on. This component makes strong assumptions that were made to simplify the first analysis of the model. Despite this, the model was able to recover major patterns of human brain and body size development and evolution, so we continue to make them here, again as a first approximation to be relaxed in the future.

The selection component assumes that the population is age-structured, at (evolving) carrying capacity, and composed of females only that reproduce clonally (i.e., asexually), which are standard assumptions in life history theory. Fitness *w* is then given by the standard expression under age structure, arranged in the form:

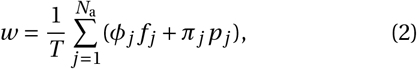

where *f* _*j*_ and *p*_*j*_ are respectively the fertility and survival probability at age *j*, with respective weights *φ*_*j*_ and *π*_*j*_ that decrease with age *j* particularly after maturity, and where generation time is *T* (Baudisch, 2005; Caswell, 1978; Hamilton, 1966). Further, the model assumes that the fertility of females is proportional to the size of reproductive tissue, defined as the number of mature ovarian follicles, which simplified the mathematical treatment but is consistent with empirical observations that human female fertility correlates with mature ovarian follicle count (Broekmans, Faddy, Scheffer, & te Velde, 2004; Chang, Chiang, Hsieh, Soong, & Hsu, 1998). Additionally, for simplicity and as a first approximation, the model assumes that the probability of surviving from one age to the next is constant; that is, having higher skill levels does not make individuals more likely to survive to the next age. As a result, the selection component entails that there is only selection for reproductive tissue but no selection for cognition, measured as skill level in the model.

These assumptions go clearly against standard thinking, but were implemented as a first attempt for simplicity and surprisingly enable the model to recover a wide array of major patterns of human development and evolution. This raises the possibility that conditions that are traditionally regarded as necessary for human evolution (e.g., selection for cognition) might not be as necessary. However, recall that this model should be interpreted as a baseline that can be used to build model competitors to identify those that best explain the data. Since such model competitions have not been done, the model predictions and explanations are preliminary rather than final.

#### Evolution

The final component is evolution, which allows the model to obtain macro-evolutionary trajectories of brain and body sizes over generations. Classic equations describing evolution do not consider a mechanistic, dynamic description of development as the brain model does. For instance, equations describing evolution in quantitative genetics assume that phenotypes are the sum of genetic contributions, which is known as Fisher’s infinitesimal model (Walsh & Lynch, 2018). Instead, the evolution component of the brain model uses recent equations that describe evolution with a mechanistic, dynamic description of development, which offers a mathematical synthesis of development and evolution (González-Forero, 2024b). This synthesis has some similarities but also fundamental differences from quantitative genetics. For instance, inheritance is not described by heritability in this framework, but by a mechanistically described ancestor-descendant covariance after the effects of social interaction have stabilised, which has different properties to heritability (González-Forero, 2024b).

The evolution component makes additional simplifying assumptions that are unrealistic, but again serve as a stepping stone for further realism. The model assumes that the effort *y*_*ia*_ to allocate energy to the growth of each tissue *i* at each age *a* is controlled by a separate genetic locus. Since the model considers three tissues (*i* ∈ {b, r, s}) and individuals are tracked from birth to 47 years of age, with ten age bins per year, this means that the model considers 3×47×10 = 1470 loci (each with a continuum of alleles, so each *y*_*ia*_ can take any real value). Because reproduction is clonal, it is unnecessary to specify how these loci are linked or whether they are in autosomes or sex chromosomes. This assumption of separate loci for each age is a genetic specification of the standard life history approach with genetic traits (control variables) that vary with age, where the genetic basis of these traits is often not made explicit (León, 1976); this assumption is also mathematically equivalent to assuming that a single locus controls how a genotypic trait varies with age over the life course (Avila & Lehmann, 2023). Given clonal reproduction, the genotypic traits are passed on to offspring without alteration barring rare mutation. These genetic idealisations are unrealistic, but are standard and as before we use them here for simplicity as a first approximation.

The model assumes that evolution occurs by mutation-fixation epochs: at the start of each epoch, all individuals have the same genotypic traits and the population attains carrying capacity, then a mutation of small effect on the genotypic traits arises, and mutants either vanish or spread to fixation while the population attains a new carrying capacity. These stylised assumptions of mutation-fixation epochs are again made for simplicity, are common in population genetics modelling, and are standard in adaptive dynamics (Dieckmann & Law, 1996; Metz, Geritz, Meszéna, Jacobs, & van Heerwaarden, 1996). These assumptions of evolution by mutation-fixation epochs are often found to yield approximately correct evolutionary trajectories if mutation is rare and of small effect (Champagnat, 2006).

As the genetic loci evolve, the development (1) of phenotypic traits is modified, which induces the evolution of mean tissue sizes and mean skill level across the life course. Despite the complex developmental dynamics, it is possible to write the evolutionary change in the average lifetime phenotype and genotype 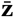 in the form of a Lande-like equation (Lande, 1979) as

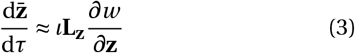

(González-Forero, 2024b). As the Lande equation, this equation is helpful to understand evolution and how it is modulated by development because it partitions evolutionary change into selection (*∂w* /*∂***z**) and ancestor-descendant covariation (**L**_**z**_, in place of Lande’s **G** matrix). Yet, unlike the Lande equation, this equation accounts for a dynamic description of development together with a description of (simplified) genetic evolution. Here, **L**_**z**_ is the matrix of “mechanistic socio-genetic covariation”:

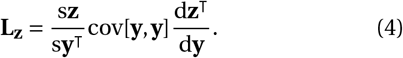

This matrix maps mutational covariation described by cov[**y, y**] onto its effects d**z**^⊺^/d**y** on the phenotypes and genotypes under selection and onto its effects s**z**/s**y**^⊺^ on the phenotypes and genotypes developed by the descendant population. The derivative d**z**^⊺^/d**y** is the developmental sensitivity of the phenotype and genotype to genotypic change, which synthetically describes all the developmental effects of changing the genotype over development, and is a mechanistic analogue of Fisher’s additive effect of allelic substitution (Lynch & Walsh, 1998). The s-derivative s**z**/s**y**^⊺^ gives the stabilised effects of perturbing the genotype on the individual’s phenotype after such effects have propagated through the population via social interactions. The framework mechanistically tracks how the developmental process maps genetic perturbations onto phenotypic effects, thus possibly generating “socio-genetic” associations (described by the entries of **L**_**z**_) that modulate the evolutionary process, by providing equations for the sensitivities above. Unlike Lande’s **G** matrix, **L**_**z**_ can be asymmetric and is always singular, the latter of which entails that evolutionary outcomes depend on **L**_**z**_ rather than only on selection.

The evolution component allows the model to show that the recovered hominin brain expansion occurs, not because a challenging ecology and cumulative culture increase selection for brain size, but because they affect development such that brain size and the only tissue under selection in the model, namely, reproductive tissue, become “socio-genetically” correlated (i.e., as described by the entries of **L**_**z**_) (González-Forero, 2024a). That is, the human brain size evolves in the model as a spandrel, or by-product of selection for reproductive tissues (González-Forero & Gómez-Robles, 2025). This conclusion is surprising but not definitive as further comparisons with model variations are necessary, particularly where selection for cognition is allowed (e.g., where survival is a function of skill level). However, this conclusion is unexpectedly consistent with empirical evidence of causally relevant genetic associations between brain related traits and reproductive traits in human females (Molz et al., 2024), human males (Bush & Goriely, 2024, 2025; Matos, Publicover, Castro, Esteves, & Fardilha, 2021), and fish (Buechel, Booksmythe, Kotrschal, Jennions, & Kolm, 2016; Corral-López et al., 2017; Kotrschal et al., 2015). Specifically, it has been found that genes associated with the shape of the braincase in human females are highly expressed in female reproductive tissues (Molz et al., 2024; i.e., pleiotropy, which can cause genetic correlations; Eq. (4)), that there is a large overlap in genes associated with cell proliferation in the human brain and testes (Bush & Goriely, 2024, 2025; Matos et al., 2021), and that artificially selecting for large brains in fish causes an associated evolution of enlarged male genitalia and of other sexual traits (Buechel et al., 2016; Corral-López et al., 2017; Kotrschal et al., 2015).

### Developmental sensitivity

We seek to quantify the extent to which genes, environment, and any other factors involved in development causally affect the developed skill level across the recovered hominin brain expansion. Causal effects can be estimated using sensitivity analyses of a model (Caswell, 2019). Such causal effects can be computed with derivatives, provided that such derivatives are causal (Henshaw, Morrissey, & Jones, 2020). For instance, if a variable *x* causally affects *y* = *f* (*x*) for some function *f* but *y* does not causally affect *x*, then causal derivatives of *x* with respect to *y* must be zero even if *f* is invertible (Lehtonen & Otsuka, 2023).

We compute developmental sensitivities using recently derived formulas for the sensitivity of a recurrence (González-Forero, 2024b), which give causal derivatives by considering a mechanistic description of the system by means of recurrence equations, allowing for arbitrarily complex and non-linear causal feedbacks (e.g., via all direct and indirect loops in Fig. 2A). This causality from derivatives is achieved by computing the effects of a perturbation on the recurrence from the time of the perturbation, specifically, by applying a perturbation at some time and following the effects of the perturbation from that time onwards (González-Forero, 2024b). This approach considers the causal arrow of time, such that perturbations can only causally affect the future, but not the past. These formulas allow for causal developmental, environmental, and social feedbacks, for instance, when a perturbation of a phenotype at age 1 affects another phenotype at age 2, which then affects the first phenotype at age 3.

We now describe the formulas we use for the sensitivity of a recurrence (González-Forero, 2024b). Despite the arbitrary developmental complexity allowed, these formulas are compact and biologically informative. The sensitivity of the phenotype to genotypic change is given by (all derivatives are evaluated at 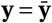

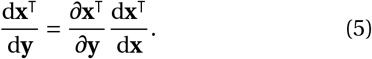

That is, the total effects of genotype on phenotype (d**x**^⊺^/d**y**) equal the direct effects of genotype on phenotype (*∂***x**^⊺^/*∂***y**) times the developmental feedbacks that ensue (d**x**^⊺^/d**x**). Partitioning the matrix d**x**^⊺^/d**y** into blocks of size 3 × 4 (since there are 3 genotypic traits and 4 phenotypes), the *a j*-th block entry gives the total effects of the genotype at age *a* on the phenotype at age *j* (that is, 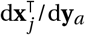, whose *ik*-th entry gives the total effect of the *i*-th genotypic trait at age *a* on the *k*-th phenotype at age *j*, d*x*_*k j*_ /d*y*_*ia*_). The sensitivity of the phenotype to changes in the parameters (d**x**^⊺^/d***ϵ***) or to social changes 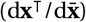 analogously given by (5) but replacing **y** for the perturbing variable.

Developmental feedbacks are given by the sensitivity of the phenotype to phenotypic change:

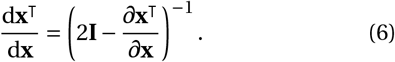

This matrix takes this form from the geometric series of matrices because it is the sum over all ages of the direct effects of phenotypes at a given age on the phenotypes at the next age (Eq. S5.1.19 of González-Forero 2024b). This matrix has the form of a classic formula of total effects of variables on themselves (Greene, 1977), but here it has underlying structure determined by the recurrence equations. Partitioning the matrix (6) into blocks of size 4 × 4 (because there are four phenotypes), the *a j*-th block entry gives the total effects of the phenotype at age *a* on the phenotype at age *j* (that is, 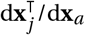, whose *ik*-th entry gives the total effect of the *i*-th phenotype at age *a* on the *k*-th phenotype at age *j*, d*x*_*k j*_ /d*x*_*ia*_).

The matrix of direct effects of the phenotype on the phenotype at all ages is:

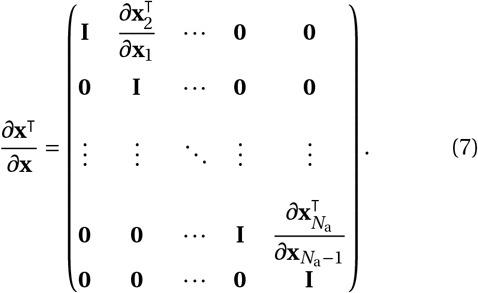

The block superdiagonal matrices 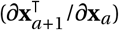 give the direct effects of the phenotype at age *a* on the phenotype at age *a* + 1. All other entries of (7) are zero except the main diagonal ones which are one, because the phenotype cannot directly affect other phenotypes at other ages unless it is itself at the same age.

Finally, the matrix of direct effects of the phenotype at age *a* on the phenotype at age *a* + 1 is

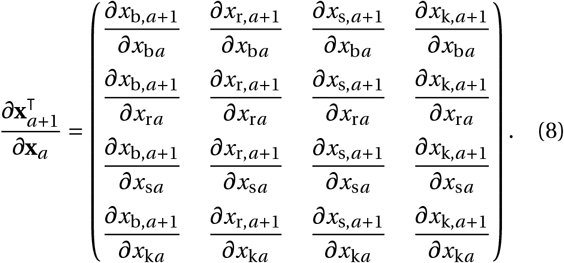

Each entry of this matrix gives the direct effect of a phenotype on another phenotype at at next age, and is obtained by differentiating Eq. (1). Once this matrix is derived, all the superdiagonal entries of (7) follow by simply evaluating (8) at each age. The direct effects of parameters (*∂***x**^⊺^/*∂****ϵ***) or social interactions 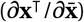 are obtained analogously as in (7) and (8) but replacing **y** for the perturbing variable and with the diagonal entries of (7) being zero because now the derivatives are with respect to other variables rather than with respect to the same variables.

For the brain model, we analyse the sensitivity of the skill level of an *in-silico* individual at a given age to changes earlier in development, and how such sensitivity has changed over evolution (Fig. 2B). We compute the sensitivity of skill level to developmentally earlier point perturbations in genotypic traits (i.e., the energy allocation efforts to grow each tissue), phenotypic traits (i.e., brain size, somatic size, mature ovarian follicle count, and skill level), the skill level of an average social partner, or the parameters affecting development; for short, we say that the parameters affecting development describe environment or physiology, as these parameters refer to environmental, metabolic, social, and maternal care properties. For instance, we compute the effect on the skill level of a 40-year-old individual caused by a genotypic change that marginally changed her energy allocation to brain growth at 1 year of age but only at that 1 year of age. Similarly, we also compute the effect on the skill level of a 40-year-old caused by a phenotypic change that marginally changed, for instance, her body size when she was 2 years old. Our approach does not need to specify how those interventions occur in practice, but only predicts their effects if the interventions were to occur (as discussed in p. 6 of Caswell, 2019). Although it may appear that the effects of an intervention would be completely diluted by random life events as the time between the intervention and the evaluated age increases, in a later section we introduce more realistic levels of randomness finding that the effects of the intervention do not necessarily wane and instead may increase as the effects of the perturbation propagate over development.

In quantifying developmental sensitivity, we only consider the effects of interventions on factors directly affecting the development of the focal individual, rather than interventions on factors that indirectly affect the focal individual’s development. For instance, in the brain model, the focal individual’s development directly depends on the skill level of social partners because of cooperation and competition, but does not directly depend on the body size, brain size, or environmental conditions experienced by social partners, which affect the social partner’s skill level and so indirectly affect the focal individual’s development. Thus, our developmental sensitivity calculations only consider the effects of intervening the skill level of social partners, but not the effects of perturbing the brain size, body size, or environmental conditions of social partners.

The model at present is likely to underestimate the developmental sensitivity to social interactions. This is because social learning has only been implicitly incorporated in the model. Thus, intervening the skill level of social partners only affects the focal individual’s development because of cooperation and competition for energy extraction, rather than via social learning. However, we do consider some aspects of social learning implicitly, specifically by intervening parameters whose value may be partly due to culture (i.e., skill effectiveness at energy extraction and environmental difficulty, which affect the shape of energy extraction efficiency vs skill level).

### Developmental elasticity

The units of the developmental sensitivities depend on the perturbed and response variables. To compare sensitivities despite their different units, a standard approach is to compute the elasticity (Caswell, 2019), which is the proportional sensitivity, so is dimensionless and is defined only for positive variables. As genotypic traits can take negative values (e.g., defined as deviations from a reference value), we compute a slight modification of elasticity in this situation, normalising by the absolute value of the genotypic traits. Specifically, we compute the elasticity of phenotype *i* at age *a* to change in *ζ* ∈ **ℝ** as

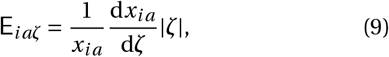

where |*ζ*| is the absolute value of *ζ* (we use the font E to distinguish it from the metabolic costs of growth denoted be *E*). When *ζ* is positive, E_*iaζ*_ is the standard elasticity of phenotype *i* at age *a* to change in *ζ*. Thus, to compare sensitivities to different types of variables, we analyse such developmental elasticities of skill level over the evolutionary brain expansion from *H. habilis* to *H. sapiens* as recovered by the brain model.

### Developmental reactivity

To quantify in a concise way the relevance of different types of variables for a phenotype of interest, we define developmental reactivity, which measures the contribution of a set of developmental variables to the total elasticity of a phenotype at a given age.

To do this, we proceed as follows. In the evo-devo dynamics framework we use (González-Forero, 2024b), the developmental map is in general of the form 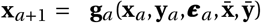. Thus, let us denote the vector of all the variables affecting development by 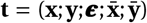. Next, we extend the notion of elasticity to let the perturbing variable be a vector (e.g., containing the variables of a certain type), so the elasticity of phenotype *i* at age *a* to change in ***ζ*** ∈ **ℝ**^*n*×1^ is

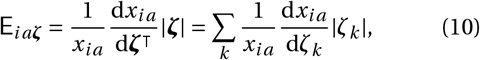

where |***ζ***| is the vector with the entries of ***ζ*** in absolute value and *ζ*_*k*_ is the *k*-th entry of ***ζ***. For instance, E_*ia***y**_ is the elasticity of phenotype *i* at age *a* to changes in genotypic traits throughout the life course and E_*ia***x**_ is the elasticity to changes in phenotypes throughout the life course. Then, we define the total elasticity of phenotype *i* at age *a* as E_*ia***t**_, which reduces to

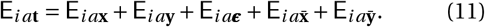

Thus, the total elasticity of a phenotype at a given age is the phenotype’s elasticity to marginally changing all the variables directly affecting development at all ages. As shown by (11), the total elasticity is the sum of elasticities to changing variables of the various types.

Using this, we define the reactivity of phenotype *i* at age *a* to change in ***ζ*** as

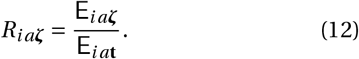

Because of (11), the reactivity *R*_*ia****ζ***_ gives the contribution of ***ζ*** to the total elasticity of phenotype *i* at age *a*. For instance, the reactivity to genotypic traits *R*_*ia***y**_ gives the contribution of genotype perturbations to the total elasticity. Similarly, *R*_*ia***x**_, 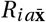, and *R*_*ia****ϵ***_ respectively give the contribution of phenotypic traits, social traits, and environment or physiology to the total elasticity of the phenotype at the indicated age.

Reactivities may be negative but add up to 1 (*R*_*ia***t**_ = 1), so reactivities offer a convenient summary of the developmental sensitivities. Although each reactivity may be interpreted as the contribution of each variable type to the total elasticity of skill level at a given age, such contributions are not independent (similarly to Caswell, 2001, p. 229).

The robustness (or canalisation) of a phenotype to perturbing a variable can be measured by one minus reactivity. Reactivity and robustness defined in this manner are individual-level properties rather than population-level quantities such as heritability. However, note that the reactivity to genetic change is not “the genetic component” of a trait, similarly to heritability which is not the genetic component of a trait. Instead, the reactivity to genetic change measures the extent to which a phenotype responds to genetic change, relative to all other possible changes in development. For other measures of robustness, see Félix and Barkoulas (2015) and Kitano (2007).

### First-order contributions to developed phenotypic differences

We consider a situation where two individuals have different phenotypes and there is an ethically justified interest in quantifying the contributions of genes and environment to these individuals’ different phenotypes. These contributions can be quantified in an *in-silico* model of development, by combining the developmental sensitivities above that use formulas for the sensitivity of a recurrence (González-Forero, 2024b) with the logic of Caswell’s life table response experiment method (Caswell, 1989) (and p. 261 of Caswell, 2001). To do this, we take a firstorder approximation of the phenotype of one individual around the conditions experienced by the other individual, which yields an approximated expression that partitions the difference in phenotypes between the two individuals into the contribution from the genetic and environmental conditions experienced.

Let 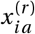 be the value of the *i*-th phenotype at age *a* for an individual that developed under conditions labelled *r*, under which the individual has genotype **y**^(*r*)^ and experiences an environment ***α***^(*r*)^ over life (in the brain model, the *a*-th entry of vector ***α***^(*r*)^ is the environmental difficulty 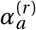 experienced at age *a*). The quantity 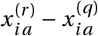 gives the difference in the *i*-th phenotype at age *a* between an individual who developed under conditions *r* and an individual who developed under conditions *q*. From Taylor’s theorem, to first order of approximation, the phenotype of the former individual is

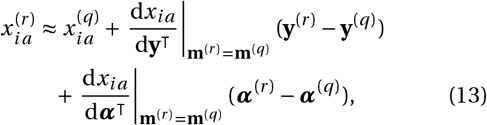

where **m**^(*r*)^ = (**y**^(*r*)^; ***α***^(*r*)^). Thus, the first-order genetic contribution to the phenotypic difference 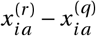 is

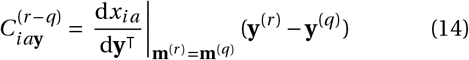

and the first-order environmental contribution to the phenotypic difference 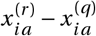 is

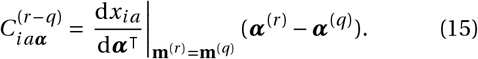

As this is a first-order approximation, this quantification is more accurate when the compared individuals experience similar genes and environments. We use the first-order contributions (14) and (15) to quantify the contributions of genes and environment to the differences in skill level between *in-silico* individuals having average bottom and top quantile adult skill levels. To generate empirically informed bottom and top skill levels, we introduce levels of genetic and environmental variation informed by empirical estimates of heritability as follows.

### Empirically informed variation

Developmental sensitivities, elasticities, reactivities, and first-order contributions can be computed at the individual level and so do not require consideration of population variability, except that first-order contributions require that the individuals compared have different phenotypes. In contrast, common approaches to quantify genetic and environmental contributions rely on heritability, which requires population variability. To compare our approach to heritability-based approaches and to apply our first-order contributions approach to empirically informed variation in skill levels, we proceed as follows.

First, we introduce genotypic and environmental variation around the average *in-silico* individual recovered in the *Homo sapiens* scenario (i.e., yellow dot in Fig. 1A and red lines in Fig. 1B-E). This average individual has genotypic traits 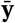 and experiences an environmental difficulty *α* at every age. A higher environmental difficulty means that individuals need a higher skill level to extract energy from the environment under ecological or cooperative challenges. At each age, we independently sample ten thousand values for both the genotypic traits and environmental difficulty from a normal distribution with means 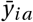 and *α* and variances 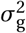 and 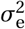, respectively. For each set of sampled genotypic traits and environmental difficulty over life, we develop an *in-silico* individual having such genotype and environment. We thus obtain a population of ten thousand *in-silico* individuals having different phenotypes distributed around the average individual that was recovered by the brain model under the *Homo sapiens* scenario in Fig. 1.

Second, we calculate the heritability of skill level over the life course among the *in-silico* individuals that develop under this genotypic and environmental variation. We use the standard definition of narrow sense heritability, namely, the ratio of additive genetic variance and phenotypic variance. The additive genetic variance is the variance in breeding value, defined as the best linear prediction (via leasts squares) of the phenotype from the genes. We thus obtain skill heritabilities that depend on the genotypic and environmental variance in the population (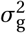 and 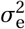).

Third, to obtain empirically informed first-order contributions, we identify values of the genotypic and environmental variances using empirical estimates of heritability. We do this by obtaining predicted heritabilities of skill level for different genotypic and environmental variances and identifying those under which the predicted heritability matches recent empirical estimates of SNP heritability of cognitive performance (Tan et al., 2024). We find that the empirically observed SNP heritability can be obtained under a variety of genotypic and environmental variances, illustrating the limited ability of heritability to quantify genetic and environmental contributions (Fig. S18). So, we use an intermediate value among the genotypic and environmental variances 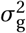 and 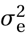 that yield a match between predicted and observed heritabilities.

This comparison with the heritability of cognitive performance does not assume that the heritability of skill level is the same as the heritability of cognitive performance in real people. Instead, as we are unaware of empirical estimates of the heritability of skill level for energy extraction, we use the most relevant empirical data we are aware of.

## Results

### Skill is little elastic to genetic change

We find that adult skill level is little elastic to change in genotypic traits in this *in-silico* replica of hominin evolution. With *H. habilis* brain and body sizes, perturbing genotypic traits at any age typically has small effects on the developed adult skill level (Fig. 3A; the computer code that generated all the figures is in González-Forero, 2026). Sensitive periods to genetic change occur early in development and coincide with the short period of brain growth (Fig. 1C, blue). With *H. sapiens* brain and body sizes, perturbing genotypic traits also has small effects on adult skill level, but perturbing genotypic traits at ages approaching the end of brain growth (approximately 10 yr; Fig. 1C, red) has more pronounced effects than at other ages (Fig. 3E).

**Figure 3:**
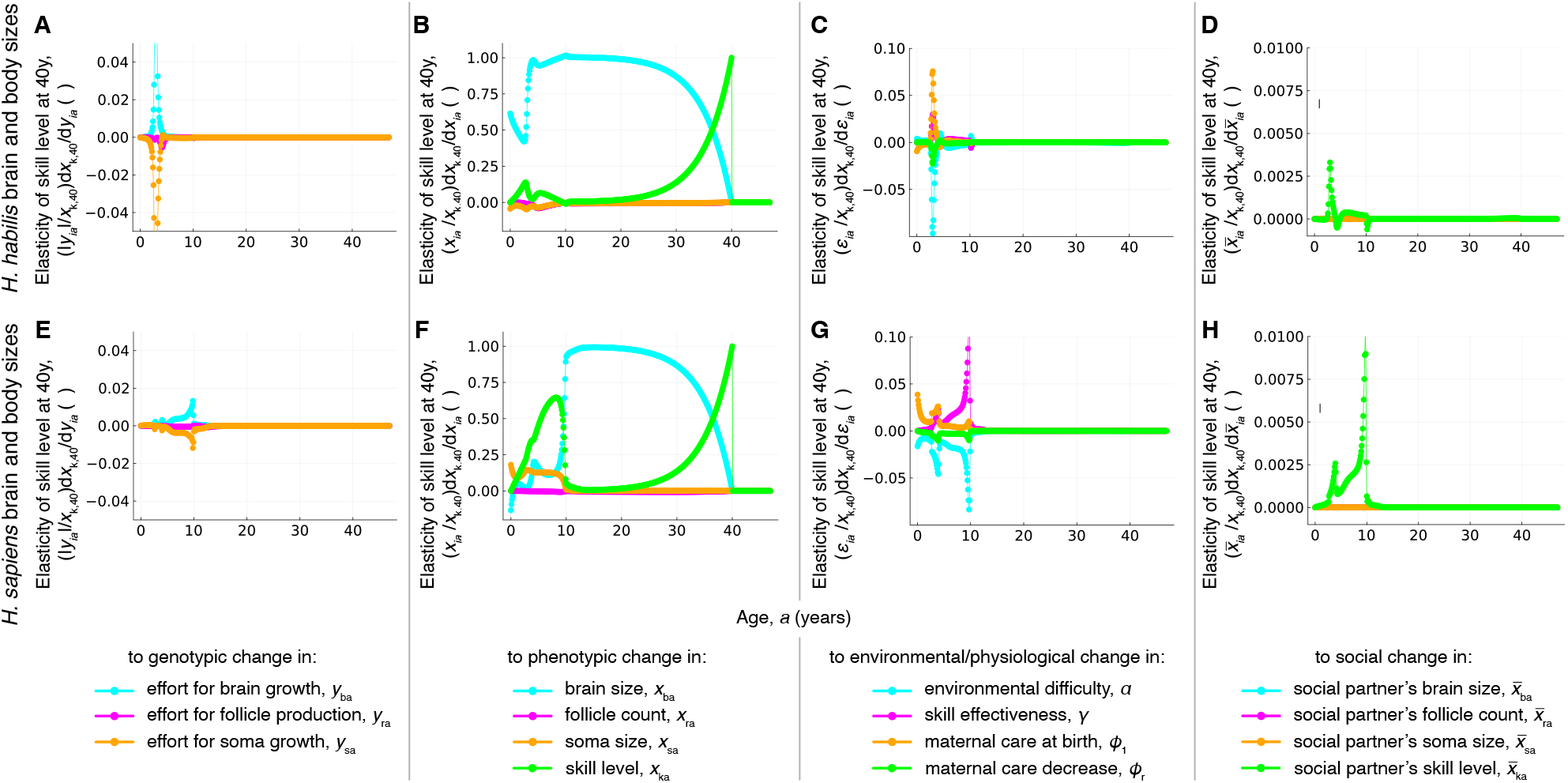
Adult skill level is most elastic to change in developmental history. Elasticity of adult skill level to (**A**,**E**) genotypic, (**B**,**F**) phenotypic, (**C**,**G**) environmental or physiological, and (**D**,**H**) social change: plots give the change in skill level at 40 yr when the indicated variable is marginally increased at the age and only at the age in the horizontal axis. Panels **A-D** are for individuals with brain and body sizes of *H. habilis* scale (i.e., for blue dots in Fig. 1B-E). Panels **E-H** are for individuals with brain and body sizes of *H. sapiens* scale (i.e., for red dots in Fig. 1B-E). In **B** and **F**, the elasticity after 40 yr is zero because the skill level at 40 yr is not affected by changes implemented in the future. The elasticity to phenotypic change is largest (**B**,**F**), followed by the elasticity to environmental (**C**,**G**), genetic (**A**), and social (**D**,**H**) change (see vertical axes).

### Skill is more elastic to developmental change

Adult skill level is more elastic to change in developmentally earlier phenotypes, exceeding the elasticity to genetic change by two orders of magnitude with either *H. habilis* or *H. sapiens* brain and body sizes (Fig. 3B,F). This is unexpected as both genotypic and phenotypic change induce developmental feedback (Eqs. S26 and S31), and both are implemented at a single age only. The analytical expressions of the sensitivity of a recurrence indicate that the sensitivities to genotypic change are relatively small because the direct effects of phenotypes on phenotypes at the next age are mostly positive (Eq. S30; originally derived in González-Forero, 2024a), whereas the direct effects of genotype on phenotypes at the next age are often negative due to trade-offs in energy allocation to growth of the various tissues (Eq. S36; originally derived in González-Forero, 2024a). For instance, increasing the genetically controlled effort to brain growth at a given age both increases brain size and decreases somatic size at the next age. These positive and negative direct effects mean that the total effect of genotype on phenotype may be positive via a path on the causal diagram but negative via another one, and the two effects cancel each other yielding little total genotypic effect on phenotype.

With *H. habilis* brain and body sizes, adult skill level is most elastic to perturbation in skill level or brain size at earlier ages, with a window of small elasticity to change in skill level (Fig. 3B) when the brain starts or ceases to grow (Fig. 1B). A larger elasticity of adult skill level to change in developmentally earlier skill level occurs as the assessed age approaches (Fig. 3B).

With *H. sapiens* brain and body sizes, adult skill level is most elastic to different phenotypic perturbations depending on the life stage (Fig. 3F). During infancy, perturbations increasing the body size of the infant have the strongest positive effect on their adult skill. During childhood, perturbations increasing the skill level of the child have the strongest positive effect on their adult skill. During adolescence and much of adulthood, perturbations increasing the brain size of the adolescent or adult have the strongest positive effect on their adult skill at the assessed age. Here adolescence is defined as the period where the individual is reproductively capable but keeps growing (approximately 12-19 yrs; Fig. 1C,D). During adulthood approaching the assessed age, perturbations increasing the skill level at those ages have the strongest positive effect on adult skill at the assessed age.

Also, during adolescence, interventions on variables other than brain size have essentially no proportional effect on adult skill level in the model. This inelasticity of adult skill to adolescent perturbation arises seemingly because, by adolescence, brain growth has ceased in the model, as perturbations occurring after brain growth arrest have vanishing effects on adult skill level except if they are on brain or skill.

### Skill is intermediately elastic to environmental change

Altering the environment and physiology has moderate proportional effects on adult skill. With *H. habilis* brain and body sizes, increasing the environmental difficulty (i.e., individuals need higher skill level to extract energy) has mostly negative proportional effects on adult skill, with a small positive proportional effect at ages when mature ovarian follicle production slows down (Fig. 3C, blue). Instead, with *H. sapiens* brain and body sizes, increasing the environmental difficulty during childhood, particularly when brain growth starts and ends, has negative effects on adult skill level (Fig. 3G, blue). The effectiveness of skills at energy extraction has largely opposite proportional effects to those of environmental difficulty (Fig. 3C,G, pink). Either increasing maternal care or decreasing the rate of maternal care decrease early in childhood typically has positive effects on adult skill level, except that with *H. habilis* brain and body sizes, increasing maternal care at birth has negative effects on adult skill level (Fig. 3C,G, orange and green).

Social interactions also have small developmental effects. With *H. habilis* brain and body sizes, increasing the social partners’ skill level during brain growth has positive proportional effects on the focal individual’s skill, but negative proportional effects when mature ovarian follicle production starts and ends (Fig. 3D). With *H. sapiens* brain and body sizes, increasing the skill level of social partners with whom the individual interacts during childhood has invariably positive proportional effects on adult skill level, particularly when brain growth starts and ends (Fig. 3H, green).

### Evolution of developmental reactivity from

#### Homo habilis to H. sapiens

The genotypic reactivity of skill level is small and has decreased over evolution. With brain and body sizes of *H. habilis* scale, the small reactivity of skill level to genotypic change peaks at around 3-4 yr (Fig. 4A, blue). Over evolution to *H. sapiens* brain and body sizes, genotypic reactivity decreases further, with a peak at around 9 yr (Fig. 4A, red). These ages coincide with the end of brain growth (Fig. 1B).

**Figure 4:**
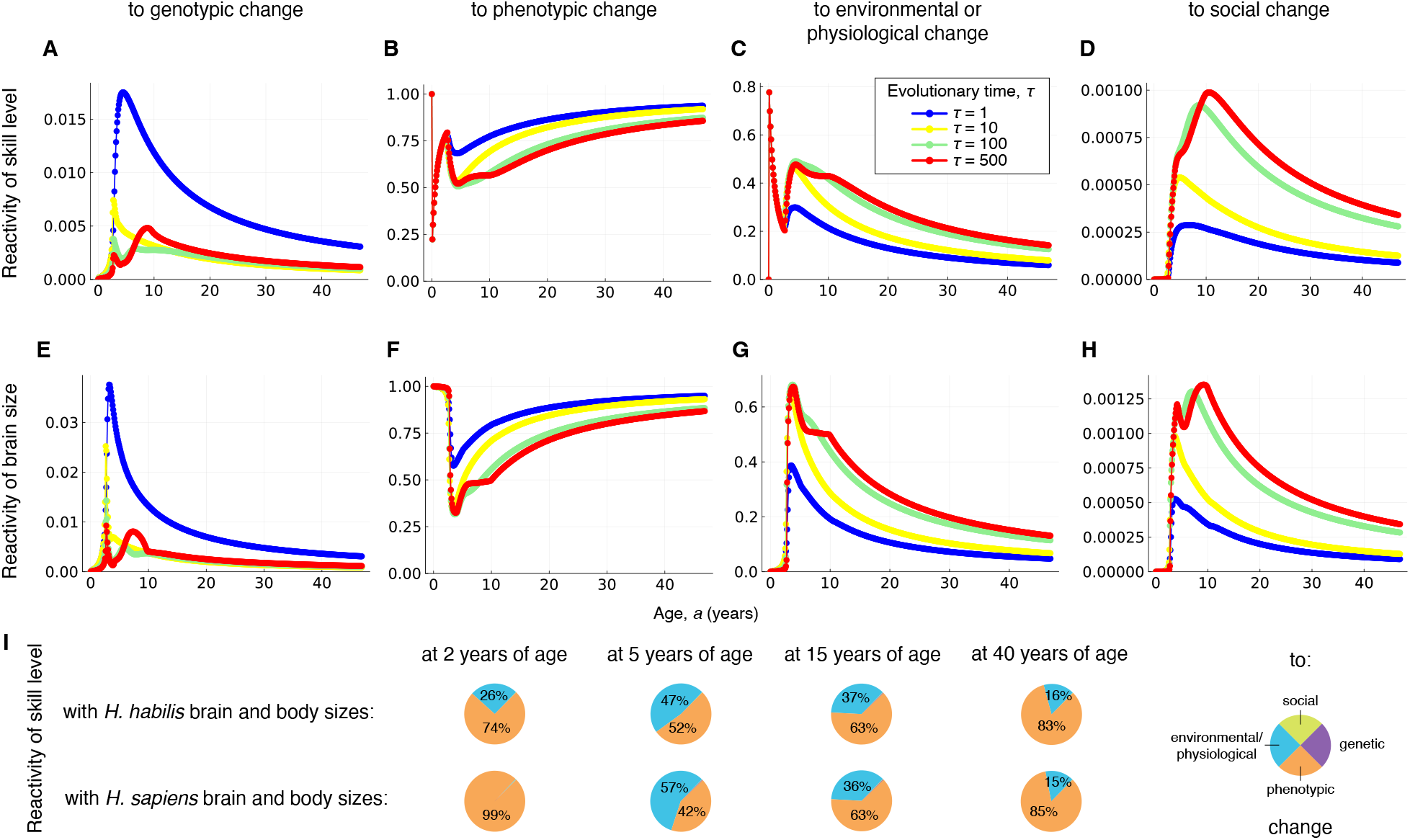
Evolution of developmental reactivity of skill level and brain size over hominin brain expansion. (**A**) Reactivity of skill level to genotypic change; for instance, the red dot at the age *a* = 10 yr gives the response of a 10-year-old’s skill level to marginally increasing all genotypic traits across all ages relative to changing all variables developmentally affecting skill level at the evolutionary time where adult brain and body sizes are of *H. sapiens* scale. The blue dot at the same age is for the evolutionary time when adult brain and body sizes are of *H. habilis* scale. Reactivity of skill level to (**B**) phenotypic, (**C**) environmental or physiological, and (**D**) social change. Skill level reactivity to genotypic and phenotypic change has decreased over evolution (**A, B**) and that to environmental, physiological, and social change has increased over evolution (**C, D**). Adult skill level remains most reactive to phenotypic change (**B**), environmental or physiological change (**C**), genotypic change (**A**), and social change (**D**) in decreasing order (see vertical axes). (**E-H**) Brain size reactivities have similar patterns. (**I**) Pie charts illustrating skill level reactivities in **A-D** at *H. sapiens* (*τ* = 500) and *H. habilis* (*τ* = 1) brain and body sizes for some ages.

The reactivity of skill level to developmental history (i.e., to changes in developmentally earlier phenotypes) is much higher despite having decreased over evolution. With *H. habilis* brain and body sizes, skill level throughout life reacts strongly to phenotypic change, and this reactivity increases with age (Fig. 4B, blue). Over evolution to *H. sapiens* brain and body sizes, the reactivity of skill level to developmental history decreases but remains substantial still increasing with age (Fig. 4B, red).

The reactivity of skill level to environmental or physiological change is moderate and has increased over evolution. With *H. habilis* brain and body sizes, skill level reacts less strongly to environmental or physiological change than to developmental history particularly after childhood, and this reactivity decreases with age (Fig. 4C, blue). Over the evolution to *H. sapiens* brain and body sizes, the reactivity of skill level to environmental or physiological change increases, but still decreases with age (Fig. 4C, red).

The reactivity of skill level to social change is very small, but it increases over evolution. With *H. habilis* brain and body sizes, skill level reacts slightly to changes in the skill level of social partners, with a developmental peak at around 5 yr (Fig. 4D, blue). Over evolution to *H. sapiens*, the social reactivity of skill level nearly quadruples, with a developmental peak at around 11 yr (Fig. 4D, red). These ages are shortly after the end of brain growth. For brain and body sizes of *H. sapiens* scale, the reactivity of skill level to social change is about three orders of magnitude smaller than to phenotypic, environmental or physiological change, and one order of magnitude smaller than to genotypic change (Fig. 4A-D, red). Such small reactivity to social change may result partly because we only consider the effects of perturbing variables that directly affect development (so we do not consider the effects of perturbing the development of social partners that then affect the focal individual) and because we do not model social learning dynamics explicitly although we model social learning implicitly via the shape of the energy extraction efficiency vs skill level.

Some of these reactivities are summarised in Fig. 4I. The sensitivities and elasticities underlying the reactivities presented are shown in Figs. S1-S16. The reactivities of the mature ovarian follicle count and somatic tissue size over the recovered human brain expansion are shown in Fig. S17.

The overall pattern of evolutionary change in the reactivity of brain size is similar to that of skill level, due to the proportionality between adult brain size and adult skill level in the model (Fig. 4E-H).

### Identifying interventions that compensate for adversity

We now introduce genetic and environmental variation around the *in-silico H. sapiens* means. Specifically, we explored a few values for the genotypic and environmental variances (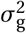 and 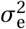) and multiple values yield a heritability of skill level that approximately match empirically observed SNP heritability for cognitive performance (Tan et al., 2024) (Fig. S18). Among these values, we use intermediate values (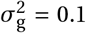 and 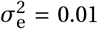). These variances cause some individuals to develop low or high adult skill level (Fig. 5A-D), and yield predicted heritabilities over the life course for the four phenotypes considered (Fig. 5E-H). For contrast, we also plot empirical estimates of SNP heritability for cognitive performance (Tan et al., 2024) and of twin-based heritability of general cognitive ability (Haworth et al., 2010) (Fig. 5H, gray triangle and squares, respectively). Twin-based empirical estimates of heritabilities for the other phenotypes we consider are: 66-97% for total brain volume (Jiska S. Peper, Boomsma, Kahn, & Pol, 2007) and 47-90% for body mass index (Elks et al., 2012), with the estimate for body mass index appearing to be larger in childhood than in adulthood (Elks et al., 2012).

**Figure 5:**
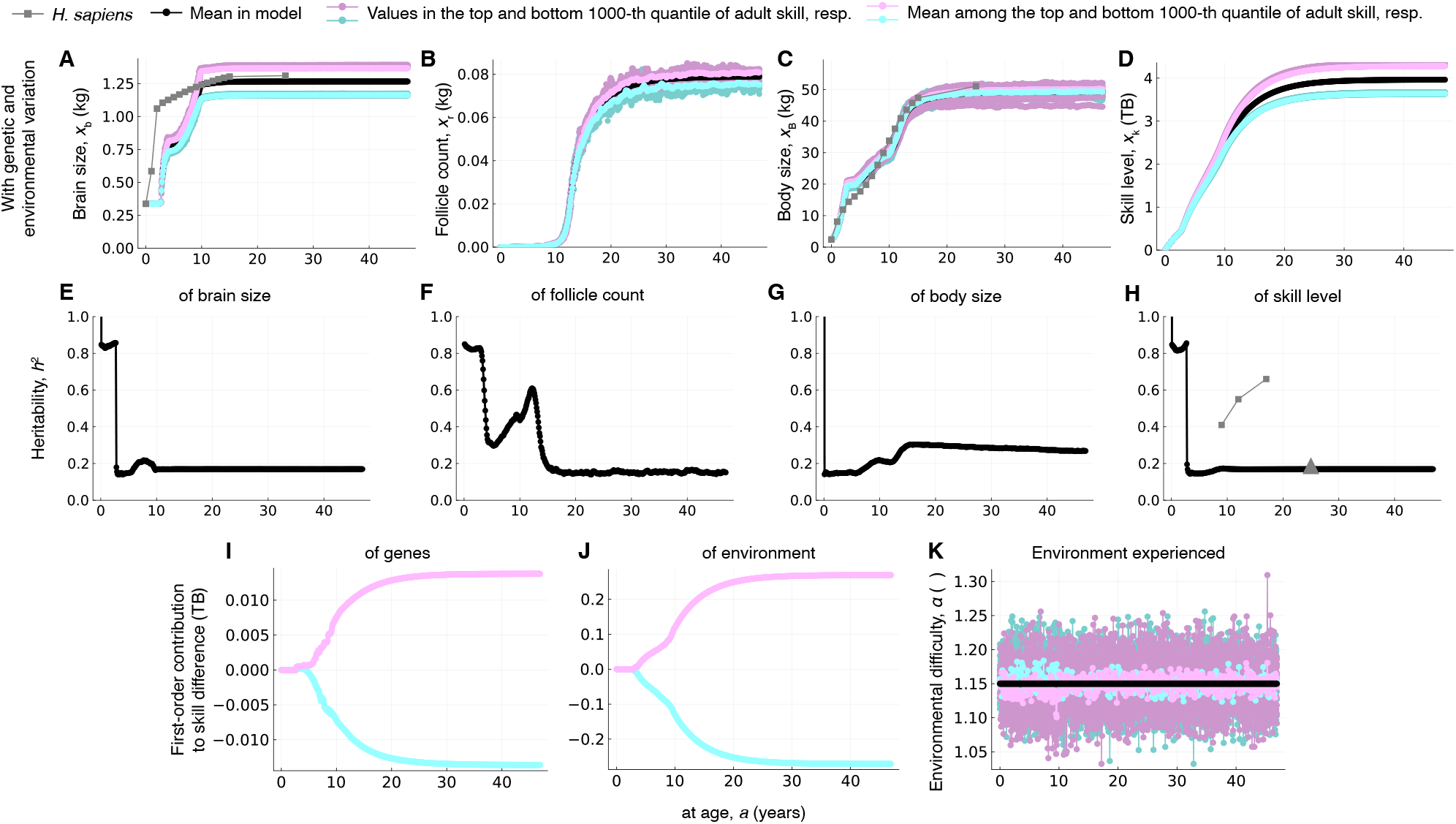
Simulating genetic and environmental adversity. For each age, we independently draw 10000 genotypic traits and environmental difficulties, normally distributed (variance 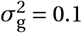 and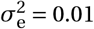, respectively) around their means given by the *H. sapiens* scenario in Fig. 1, under which we develop 10000 individuals. (**A-D**) Phenotypes of individuals in the top (dark pink) and bottom (dark blue) 1000th quantile of adult skill level (with respective means in light pink and light blue). Bottom-quantile individuals develop (**A**) smaller adult brain size, (**B**) lower adult fertility, (**C**) approximately average adult body size, and (**D**) smaller adult skill level than the mean (black). (**E-H**) Resulting heritabilities across the 10000 individuals; these heritabilities depend on 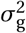 and 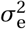(Fig. S18). We use values of 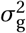 and 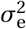 that yield an adult heritability of skill level (**H**, black) that roughly matches the empirical estimate of Tan et al. (2024) of SNP heritability of cognitive performance (**H**, gray triangle); however, other values 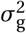 and 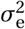 also yield a similar heritability of adult skill level (Fig. S18). For comparison, the phenotypic-based empirical estimates of heritability of general cognitive ability of Haworth et al. (2010) are also shown (**H**, gray squares). First-order contribution of (**I**) genes and (**J**) environment to the difference in skill level between the average top-quantile individual (light pink) or bottom-quantile individual (light blue) and the population-average individual, showing that the environment contributes about 15 times more to the simulated differences than genes (about 0.3 TB compared to 0.02 TB). However, this environmental-to-genetic contribution ratio varies widely (from −20 to 267) for other 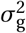 and 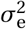 that yield the same adult skill heritability (Fig. S18), illustrating the limited ability of heritability to identify genetic contribution. Specifically, we plot the first-order (**I**) genetic 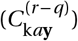 and (**J**) environmental 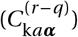 contributions to the difference in skill level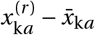, where the genotype 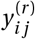 and environment 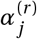 for a given age *j* are the mean among the values for individuals in the top (light pink) or bottom (light blue) 1000-th quantile of adult skill level, and the reference conditions *q* are those of the average individual in the population. (**K**) Environment experienced by the population average individual (black), top (dark pink) and bottom (dark blue) quantile individuals, and by respective mean top (light pink) and bottom (light blue) quantile individuals. The average top-quantile individual thus experiences an easier environment in childhood than the average bottom-quantile individual.

Next, we quantify the causal, mechanistic contribution of genes and environment to the simulated differences in adult skill level. For the genotypic and environmental variation used, we find that the environment causally contributes about 15 times as much as genes to the developed differences in adult skill level between average top-and bottom-skilled *in-silico* individuals (i.e., the genes of the average top-skilled individual increase their adult skill level by ≈ 0.02 TB, whereas the environment increases it by ≈ 0.3 TB; Fig. 5I,J, pink at 40 yr— the genes and environment of the average bottom-skilled individual decrease their adult skill level by similar magnitudes; Fig. 5I,J, blue). This exemplifies that heritability, reactivity, and genetic contributions measure different properties: reactivity refers to what would happen after intervention, whereas both heritability and genetic contributions refer to realised variation; however, heritability refers to the fraction of phenotypic variation that is additive genetic, whereas genetic contributions refer to how much phenotypic differences are caused by differences in genes.

The first-order contribution of genes to the differences in skill level between average top-and bottom-skilled individuals during early childhood is nearly zero as skill differences only become appreciable after about 5 years of age (Fig. 5I). The first-order contribution of the environment to the differences in skill level becomes appreciable earlier in childhood, after about 2-3 years of age.

Subsequently, we identify interventions that compensate for the simulated adversity. We implement environmental interventions in the direction of predicted steepest increase in adult skill level, given by the sensitivity of adult skill to environmental change (Fig. 6A, green). These interventions make the environment less difficult during childhood, particularly when brain growth starts and ends (Fig. 6A, green). Under the intervened environment, bottom-skilled individuals, who have genes that are detrimental to their skill, develop higher skill level than otherwise top-skilled individuals (Fig. 6B-E, green).

**Figure 6:**
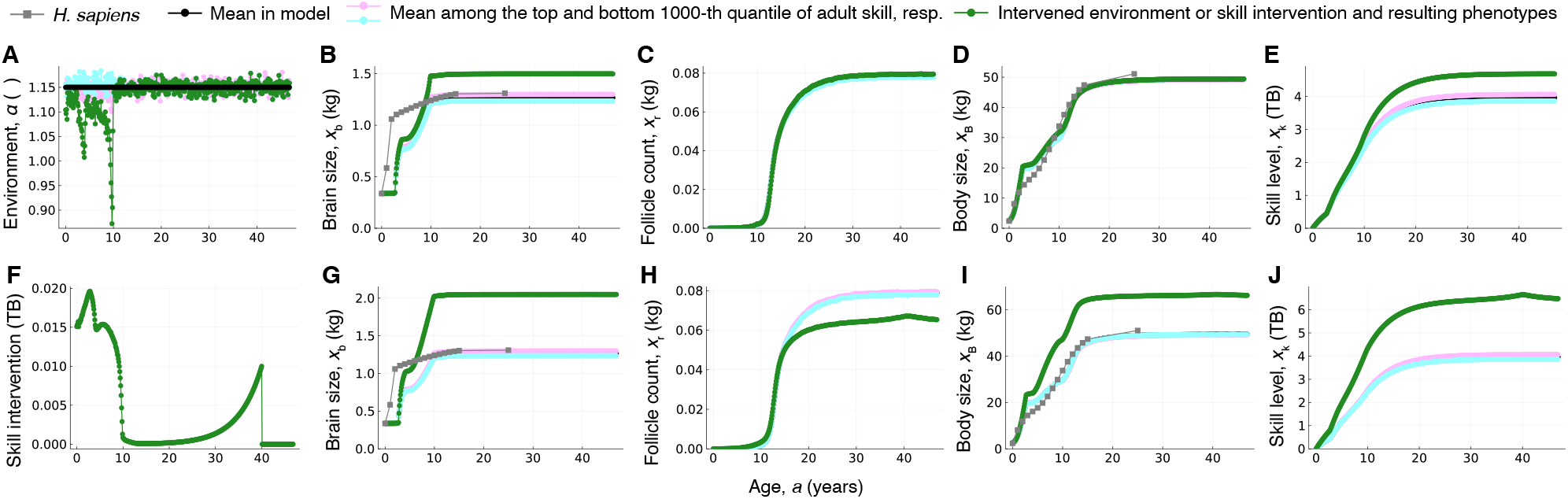
Slight interventions in developmental history compensate for genetic and environmental adversity. (**A**) Environmental difficulty on average: across the population (black), among the bottom (light blue) and top (light pink) 1000th quantile of adult skill level, and among the bottom 1000th quantile of adult skill level facing environmental intervention (green). The environment is intervened in the direction of steepest increase of adult skill level (i.e., the environment in green is that in blue plus the sensitivity of adult skill level with respect to environmental difficulty). (**B-E**) Corresponding phenotypes developed. Under the intervened environment, bottom-quantile individuals (green) develop (**B**) larger adult brain size, (**C**) average fertility, (**D**) average adult body size, and (**E**) higher adult skill level than top-quantile individuals (pink). (**F**) Rather than intervening the environment, here skill level is slightly intervened in the direction of steepest increase of adult skill level (skill level at each age is increased by 0.01 times the sensitivity of adult skill level with respect to skill level at that age). (**G-J**) Corresponding phenotypes developed. Bottom-quantile individuals (green), who have genes detrimental to adult skill, develop (**G**) much larger adult brain size, (**H**) low fertility, (**I**) larger adult body size, and (**J**) substantially higher adult skill level than top-quantile individuals (pink).

Now, rather than intervening the environment, we slightly intervene skill level earlier in life in the direction of predicted steepest increase in adult skill level, given by the sensitivity of adult skill to earlier skill level (Fig. 6F). These interventions slightly increase skill level during childhood, particularly when brain growth starts, and as the assessed age approaches. These slight interventions in skill produce a much larger compensation in adult skill level than the environmental interventions (Fig. 6G-J). This stronger compensation in skill requires slighter interventions because of the higher elasticity of skill level to developmental history than to the environment. This higher elasticity entails that interventions in skill level in the same direction but of greater magnitude disrupt development (Fig. S19). In practice, these interventions may occur exogenously, for instance, via a supportive learning environment, or endogenously, via developmental noise in brain connectivity.

## Discussion

Our findings indicate that adult human skill level may be much more sensitive to the individual’s developmental history than to other factors, including genetic change, environmental change, and changes in the skill level of social partners. Developmental history here refers to developmentally earlier phenotypes, which can be understood as storing the effects that all developmentally relevant factors (genotype, environment, social context, developmentally earlier phenotypes, and how they all interact during development) have had on the individual until a given age.

Our finding that the genotype has limited effects on adult skill level is surprising but consistent with recent findings of very small genetic effects on complex phenotypes in humans, where confounders have been increasingly removed from classic estimates of larger effects (Berg et al., 2019; Sohail et al., 2019). Specifically, recent estimates for cognitive performance find a direct genetic effect heritability of 0.233 and a polygenic index with a coefficient of determination of 0.002 (Supplementary Tables 5 and 9 of Tan et al., 2024). However, we find that a single value of SNP heritability is consistent with widely divergent genetic and environmental contributions to trait differences. This illustrates a limited ability of heritability to quantify genetic contributions.

Our finding that developmental history has the strongest effects on adult skill level is agnostic to what factors change developmental history. Those factors may be endogenous, such as developmental noise (Freund et al., 2013; Mitchell, 2018; Plomin & Kawakami, 2025; Turkheimer & Waldron, 2000), or exogenous, such as targeted interventions (Sparling et al., 2005). The strong sensitivity to developmental history entails that intervening it can be beneficial if done sufficiently slightly but detrimental if done too strongly. This highlights the potential usefulness of predictive approaches to identify desired interventions.

Our results indicate that developmental drivers of cognition are not necessarily reliable indicators of its evolutionary drivers. Indeed, we find that developmental effects can be opposite to evolutionary effects. Specifically, increasing environmental difficulty often *decreases* skill level over development, whereas increasing environmental difficulty *increases* skill level over evolution (Figs. 6 and V of González-Forero et al., 2017). This contrasts with previous expectations that developmental drivers of cognition may inform about its evolutionary drivers (Ashton, Ridley, Edwards, & Thornton, 2018).

Our results indicate that developmental history and environmental change have an opposite age-specific pattern in influencing skill level. In early life in *H. sapiens* (up to approximately 10 yr), skill level is influenced by developmental history and environmental variation in roughly equal proportions (approximately 50%) (Fig. 4B,C). As individuals age, skill level becomes increasingly reactive to developmental history and insensitive to environmental change, with only 15% of skill-level reactivity to environmental change at 40 yr.

Our results suggest that the predominant role of developmental history in affecting skill level may have been shared with many *Homo* species. According to our results, the proportion of factors influencing skill level in *H. sapiens* would have been similar 1.1 Myrs after the origin of the genus *Homo*, roughly corresponding with the *H. erectus* time period (at evolutionary time *τ* = 100; Fig. 4).

We find a reduction in the reactivity to genetic change over evolution and an increase in the reactivity to non-genetic factors, which is consistent with the view that humans have evolved increased plasticity in skill level and brain anatomy (Drennan et al., 2026; Gómez-Robles, Hopkins, Schapiro, & Sherwood, 2015; Gómez-Robles, Hopkins, & Sherwood, 2013; Hrvoj-Mihic, Bienvenu, Stefanacci, Muotri, & Semendeferi, 2013; Snell-Rood, 2013). Our results are also consistent with the view that a protracted brain development in *H. sapiens* allows for stronger environmental influences (Gómez-Robles, Nicolaou, Smaers, & Sherwood, 2024). At *H. habilis* brain size, skill level substantially reacts to environmental and social factors particularly up to 5 yr, roughly coinciding with the estimated weaning age (Robson & Wood, 2008). Skill level in early *Homo* species would then be comparatively little influenced by environmental factors after weaning, which is consistent with findings that many primate species reach adult skill levels around weaning age due to the lack of post-weaning provisioning (Heldstab, Isler, Schuppli, & van Schaik, 2020). However, at *H. sapiens* brain size, environmental and, to a lesser extent, social factors can influence skill level until almost 15 yr, well into adolescence. This finding is consistent with previous studies indicating late brain maturation in humans (D. J. Miller et al., 2012; Sotiras et al., 2017) associated with late acquisition of adult skill levels (Kaplan, Hill, Lancaster, & Hurtado, 2000; Schuppli, Isler, & van Schaik, 2012). That is, *H. sapiens* brains and skill levels can be strongly influenced by environmental factors at an age at which *H. habilis* skill level would have been largely insensitive to influences other than developmental history (Fig. 4I).

Our results also find a window of sensitivity as brain growth ceases. This finding is partly consistent with empirical work documenting adolescence as a sensitive period in mammals and birds (Sachser, Hennessy, & Kaiser, 2018; Sachser, Zimmermann, Hennessy, & Kaiser, 2020), but our predicted window of sensitivity closes before our recovered adolescence as by then brain growth has ceased in the model. Previous work has evaluated whether sensitive periods are adaptive (Sachser et al., 2018) and whether they evolve in response to an organism’s uncertainty about the future (Walasek, Panchanathan, & Frankenhuis, 2025). The sensitive periods we recover do not evolve because of selection for them, but are a “spandrel” or by-product of the developmental process. This is because the model so far only involves selection for mature ovarian follicles, so there is no selection for sensitive periods in the model. This does not mean that the recovered sensitive periods are maladaptive in the model (i.e., that their total effects on fitness are negative), only that they are not an adaptation under the textbook definition of adaptation as a trait that evolved because of direct selection for the trait (Futuyma & Kirkpatrick, 2017).

In addition to heritability, another quantity that has been suggested as being informative regarding a trait’s ability to evolve is the additive genetic coefficient of variation, defined as 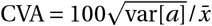, where var[*a*] is the additive genetic variance of the trait and 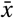 is the trait’s population mean (Houle, 1992). Empirical estimates have found small CVAs for human brain size (of 7.8) relative to other volumetric traits (around 15 to 27), which has been interpreted as suggesting stabilising rather than directional selection for brain size (G. F. Miller & Penke, 2007). We find a CVA of 1.03 for adult brain size with the used genotypic and environmental variation. However, interpreting this value in terms of selection is challenging, as illustrated by the fact that there is no selection for brain size in the model, neither balancing nor directional. Our results should be interpreted as tentative rather than conclusive. This is because conclusive results with this approach can only come after extensive empirical and statistical tests are accomplished, specifically by competing models making different assumptions (Cranmer et al., 2020). At present, our results should thus be interpreted as providing methods and a baseline model to pursue such more conclusive studies.

Overall, our results suggest a predominant role of developmental history in shaping adult skill level in humans and uncover opportunities for predicting interventions with specific timing and magnitude to mitigate and overcome adverse conditions that would otherwise yield detrimental developmental outcomes.

## Acknowledgements

MGF thanks Andy Gardner for continued support that led to this work. We thank Elisabeth Zimmermann and six anonymous reviewers for comments that were very helpful greatly to improve the paper.

## Author Contributions

MGF conceived the work and did the mathematical and computational analyses. MGF and AGR analysed the results and wrote the paper.

## Data Availability Statement

No data were collected in this study. All data used were previously published in references provided in the text. All computer codes used to obtain the results are available at https://doi.org/10.5281/zenodo.23393134 (González-Forero, 2026).

