## Supplementary Information for "Mechanistic quantification of the effects of genes, environment, and development on human behaviour"

Mauricio González-Forero*[1,2] and Aida Gómez-Robles†[3]

[1]School of Biology, University of St Andrews, Dyers Brae, St Andrews, KY16 9TH, Fife, UK
[2]Present address: Konrad Lorenz Institute for Evolution and Cognition Research, Klosterneuburg A-3400, Austria
[3]Department of Anthropology, University College London, 14 Taviton St, London, WC1H 0BW, UK

#### Contents

**S1 Model discussion** **S2**

**S2 Overview of Method details** **S2**

**S3 Development component of the brain model** **S3**
S3.1 Metabolic part . . . . . S3
S3.1.1 Energy budget for tissue growth, $B_{\text{syn}}$ . . . . . S3
S3.1.2 Developmental map for tissue mass, $g_{ia}$ for $i \in \{\text{b}, \text{r}, \text{s}\}$ . . . . . S3
S3.1.3 Brain's energy budget for learning, $B_{\text{syn,k}}$ . . . . . S4
S3.1.4 Developmental map for skill level, $g_{\text{k}a}$ . . . . . S4
S3.2 Behavioural part . . . . . S4
S3.3 Summary . . . . . S5

**S4 Derivation of developmental sensitivities** **S5**
S4.1 Developmental sensitivity to parameters in the brain model . . . . . S5
S4.2 Developmental sensitivity to developmental history . . . . . S7
S4.3 Developmental sensitivity to the genotype . . . . . S10
S4.4 Developmental sensitivity to social interactions . . . . . S11

**S5 Heritability** **S12**

**S6 Code correction** **S13**

**S7 Supplementary Figures** **S15**

### S1 Model discussion

The brain model is relatively complex, which raises concerns that are standard with complex models, specifically, unfalsifiability, underdeterminacy, and overfitting. First, an important concern with models with many parameters is that they may be able to fit any data, thereby being unfalsifiable (Burnham & Anderson, 2002; Dyson, 2004) (for an interactive illustration, see https://demonstrations.wolfram.com/FittingAnElephant/). This concern is of particular relevance when a model is based on little theory and key parameters are not empirically measured but estimated by model fitting, so such a model has largely no (active) constraints that prevent it from fitting any data. The brain model has many parameters (26 that affect the output) but has a particular form derived from mechanistic considerations, with about half of the parameters having values from empirical estimates (12, including 7 that pertain to metabolic costs, which are thought to be key given the high metabolic cost of brain tissue). This mechanistic form and empirically estimated parameter values appear to make the model actively constrained, a technical term that entails that the model is unable to generate any observation. For instance, the model has so far been unable to recover brain growth in the first year of age, and ontogenetic patterns do not exactly match observation (Fig. 1B,D; González-Forero, 2024a; González-Forero, Faulwasser, & Lehmann, 2017; González-Forero & Gardner, 2018; González-Forero & Gómez-Robles, 2025). This inability to generate any pattern makes the model rather easily falsified, which is an advantage to identify gaps in understanding (McElreath, 2018).

Second, a concern with fitting complex models is that if the model has more parameters than data points, the problem may be underdetermined, which means that multiple parameter values could fit the data equally well. Similarly, this problem is particularly relevant when a model is relatively unconstrained. This problem is relevant here as the brain model has many parameters but is fitted to sparse, fossil data. However, underdeterminacy has not been a problem in the model, as the recovered adult hominin-brain and body sizes with associated human life history occur only under unique parameter combinations, with fit quickly falling away from them, at least for the region of the parameter space evaluated (González-Forero & Gardner, 2018). Again, this is presumably because the model's mechanistic basis and empirically estimated parameter values constrain it sufficiently to prevent underdeterminacy.

Third, complex models may be overfit, where despite fitting the training data well, they predict out of sample data poorly. This is an important yet still unaddressed challenge for the brain model. Methods exist to estimate overfitting, such as cross-validation, where the model is trained excluding some of the data to evaluate its ability to predict the excluded data (McElreath, 2020). The complexity of the brain model has prevented such analyses but substantial improvements in its runtime make those analyses feasible in the near future (González-Forero, 2024a). Overall, at present, this brain model mechanistically, falsifiably, and in a determined manner recovers many, but not all, major patterns of human development and evolution while mechanistically describing skill ontogeny, but the model's level of overfitting remains to be assessed.

### S2 Overview of Method details

We use the brain model of González-Forero et al. (2017) and González-Forero and Gardner (2018), which González-Forero (2024a) implemented into the evo-devo dynamics framework of González-Forero (2024b) to model the evo-devo dynamics of human brain size. Using the model results of González-Forero (2024a) with a slight code correction described in section S6, we compute the developmental sensitivities and elasticities of the phenotypes described by the brain model, and how these developmental sensitivities and elasticities change over hominin evolution.

We arrange the remainder of this Supplementary Information as follows. First, we provide the mathematical description of the development component of the brain model, which specifies a system of recurrence equations given by the developmental map (section S3). Second, we derive the developmental sensitivities in the brain model to the parameters affecting development, to developmental history, to the genotype, and to social interactions (section S4). The text of sections S3, S4.1, S4.2, S4.3, and S4.4 is taken from the Supplementary Information of González-Forero (2024a) and is repeated here with some rearrangement for the reader's convenience. Third, we describe how we calculate heritability (section S5). Fourth, we describe the code correction we implement relative to González-Forero (2024a) (section S6). Fifth, we provide supplementary figures (section S7).

We use the following notation from matrix calculus throughout. The Jacobian matrix of a vector $\mathbf{a} \in \mathbb{R}^{n\times 1}$

with respect to a vector $\mathbf{b} \in \mathbb{R}^{m\times 1}$ in its standard or transposed form is, respectively,

$$\frac{\partial \mathbf{a}}{\partial \mathbf{b}^\intercal} = \begin{pmatrix} \frac{\partial a_1}{\partial b_1} & \cdots & \frac{\partial a_1}{\partial b_m} \\ \vdots & \ddots & \vdots \\ \frac{\partial a_n}{\partial b_1} & \cdots & \frac{\partial a_n}{\partial b_m} \end{pmatrix} \in \mathbb{R}^{n\times m} \quad \text{or} \quad \frac{\partial \mathbf{a}^\intercal}{\partial \mathbf{b}} = \begin{pmatrix} \frac{\partial a_1}{\partial b_1} & \cdots & \frac{\partial a_n}{\partial b_1} \\ \vdots & \ddots & \vdots \\ \frac{\partial a_1}{\partial b_m} & \cdots & \frac{\partial a_n}{\partial b_m} \end{pmatrix} \in \mathbb{R}^{m\times n}. \tag{S1}$$

The transpose of $\partial \mathbf{a}/\partial \mathbf{b}^\intercal$ is $(\partial \mathbf{a}/\partial \mathbf{b}^\intercal)^\intercal = \partial \mathbf{a}^\intercal/\partial \mathbf{b}$. The analogous notation applies for total derivatives.

### S3 Development component of the brain model

We now describe the developmental map of the brain model. The text of this section is taken from the Supplementary Information of González-Forero (2024a) and is repeated here with some rearrangement for the reader's convenience.

The brain model considers four phenotypic traits (state variables), namely the mass of brain, reproductive, and somatic tissues and the skill level at each age ($N_\mathrm{p} = 4$). The model additionally has three genotypic traits (control variables), namely the energy allocation effort to the growth of brain, reproductive, and somatic tissues across life ($N_\mathrm{g} = 3$). Phenotypic traits have developmental dynamics modulated by the genotypic traits. In turn, genotypic traits have evolutionary dynamics that alter the developmental dynamics of phenotypic traits. The developmental dynamics are given by the developmental map $\mathbf{g}_a$ ($a \in \{1,\ldots,N_\mathrm{a}\}$).

#### S3.1 Metabolic part

##### S3.1.1 Energy budget for tissue growth, $B_\mathrm{syn}$.

We quantify an individual's energy budget for tissue growth by the growth metabolic rate, which is the rate of heat release by the body at rest due to tissue production and equals the resting metabolic rate minus the metabolic rate due to tissue maintenance. A mutant's phenotypic trait $x_{ia}$ is her mass of tissue $i \in \{\mathrm{b,r,s}\}$ at age $a$, where $i \in \{\mathrm{b,r,s}\}$ refers to brain, reproductive, and somatic tissues, respectively. The mutant's fourth phenotypic trait $x_{\mathrm{k}a}$ is her skill level at age $a$. Thus, a mutant's growth metabolic rate at age $a$ is

$$B_\mathrm{syn}(\mathbf{x}_a, \bar{x}_{\mathrm{k}a}, a) = Ke(x_{\mathrm{k}a}, \bar{x}_{\mathrm{k}a}, a)x_{\mathrm{B}a}^\beta - \sum_{i\in\{\mathrm{b,r,s}\}} x_{ia}B_i$$

where the mutant's resting metabolic rate is $Ke(x_{\mathrm{k}a}, \bar{x}_{\mathrm{k}a}, a)x_{\mathrm{B}a}^\beta$ from Kleiber's law, with $K$ and $\beta$ being the Kleiber's law parameters. A mutant's body mass at age $a$ is $x_{\mathrm{B}a} = \sum_{i\in\{\mathrm{b,r,s}\}} x_{ia}$. The mutant's energy extraction efficiency (EEE) is $e(x_{\mathrm{k}a}, \bar{x}_{\mathrm{k}a}, a) \in [0,1]$, which depends on the mutant individual's skill level and on the skill level of social partners of the same age. EEE decreases exogenously with age $a$ to describe the effect of diminishing maternal care with age. The mutant's maintenance metabolic rate is $\sum_{i\in\{\mathrm{b,r,s}\}} x_{ia}B_i$. The mass-specific metabolic cost of maintenance of tissue $i$ is $B_i$.

##### S3.1.2 Developmental map for tissue mass, $g_{ia}$ for $i \in \{\mathrm{b,r,s}\}$.

We now specify the developmental map for tissue mass, $g_{ia}$ for $i \in \{\mathrm{b,r,s}\}$. The fraction of growth metabolic rate allocated to the production of tissue $i$ by a mutant individual of age $a$ is $q_{ia}(\mathbf{y}_a)$ for $i \in \{\mathrm{b,r,s}\}$. We let

$$q_{ia}(\mathbf{y}_a) = \frac{\exp(y_{ia})}{\sum_{i\in\{\mathrm{b,r,s}\}} \exp(y_{ia})}, \tag{S2}$$

for $i \in \{\mathrm{b,r,s}\}$ where the genotypic trait $y_{ia} \in (-\infty,\infty)$ is the energy allocation effort (relative to a baseline value) devoted to growth of tissue $i$ at age $a$. González-Forero and Gardner (2018) used $q_{ia}(\mathbf{y}_a)$ as control variables, but here we use $y_{ia}$ as control variables because numerical solution is easier as $y_{ia}$ does not have to be truncated to remain between zero and one and instead can take any real value (i.e., there are no mutational constraints). From Eq. (S2), we have

$$q_{ia}(\mathbf{y}_a) \in [0,1] \text{ and } \sum_{i\in\{\mathrm{b,r,s}\}} q_{ia}(\mathbf{y}_a) = 1 \tag{S3}$$

for all $i \in \{\mathrm{b},\mathrm{r},\mathrm{s}\}$ and all $a \in \{1,\ldots,N_\mathrm{a}\}$, and similarly for the resident (or average) $q_{ia}(\bar{\mathbf{y}}_a)$. We can now specify the developmental map for tissue mass. The size of tissue $i \in \{\mathrm{b},\mathrm{r},\mathrm{s}\}$ of a mutant individual at age $a+1$ is

$$x_{i,a+1} = g_{ia}(\mathbf{z}_a, \bar{x}_{\mathrm{k}a}, a) = \begin{cases} x_{ia} + \dfrac{q_{ia}(\mathbf{y}_a)}{E_i} B_\mathrm{syn}(\mathbf{x}_a, \bar{x}_{\mathrm{k}a}, a) & \text{if } x_{ia} + \dfrac{q_{ia}(\mathbf{y}_a)}{E_i} B_\mathrm{syn}(\mathbf{x}_a, \bar{x}_{\mathrm{k}a}, a) \geq 0 \\ 0 & \text{otherwise.} \end{cases} \tag{S4}$$

where $E_i$ is the heat released for producing an average unit of mass of tissue $i \in \{\mathrm{b},\mathrm{r},\mathrm{s}\}$.

##### S3.1.3 Brain's energy budget for learning, $B_\mathrm{syn,k}$.

We quantify the brain's energy budget for learning by the learning metabolic rate, which is the rate of heat release by the brain at rest due to learning and equals the brain metabolic rate allocated to skills minus the brain metabolic rate due to skill maintenance (memory). For a mutant of age $a$, the learning metabolic rate is

$$B_\mathrm{syn,k}(\mathbf{z}_a, \bar{x}_{\mathrm{k}a}, a) = s_\mathrm{k} B_\mathrm{rest,b}(\mathbf{z}_a, \bar{x}_{\mathrm{k}a}, a) - x_{\mathrm{k}a} B_\mathrm{k}, \tag{S5}$$

where $B_\mathrm{rest,b}(\mathbf{z}_a, \bar{x}_{\mathrm{k}a}, a)$ is the brain metabolic rate, $s_\mathrm{k}$ is the fraction of brain metabolic rate allocated to skills, and $B_\mathrm{k}$ is the metabolic cost of learning an average unit of skill. From energy conservation, the brain metabolic rate is

$$B_\mathrm{rest,b}(\mathbf{z}_a, \bar{x}_{\mathrm{k}a}, a) = x_{\mathrm{b}a} B_\mathrm{b} + \left[g_{\mathrm{b}a}(\mathbf{z}_a, \bar{x}_{\mathrm{k}a}, a) - x_{\mathrm{b}a}\right] E_\mathrm{b}. \tag{S6}$$

Indeed, $x_{\mathrm{b}a} B_\mathrm{b}$ is the rate of heat release by the brain due to maintenance of existing brain tissue, and $\left[g_{\mathrm{b}a}(\mathbf{z}_a, \bar{x}_{\mathrm{k}a}, a) - x_{\mathrm{b}a}\right] E_\mathrm{b}$ is the rate of heat release by the brain due to production of new brain tissue.

##### S3.1.4 Developmental map for skill level, $g_{\mathrm{k}a}$.

We can now specify the developmental map for skill level. The skill level of a mutant individual at age $a+1$ is

$$x_{\mathrm{k},a+1} = g_{\mathrm{k}a}(\mathbf{z}_a, \bar{x}_{\mathrm{k}a}, a) = \begin{cases} x_{\mathrm{k}a} + \dfrac{1}{E_\mathrm{k}} B_\mathrm{syn,k}(\mathbf{z}_a, \bar{x}_{\mathrm{k}a}, a) & \text{if } x_{\mathrm{k}a} + \dfrac{1}{E_\mathrm{k}} B_\mathrm{syn,k}(\mathbf{z}_a, \bar{x}_{\mathrm{k}a}, a) \geq 0 \\ 0 & \text{otherwise.} \end{cases} \tag{S7}$$

This expression means that a mutant's skill level at age $a+1$ is the individual's skill level at the previous age plus the brain's rate of heat release due to skill growth $B_\mathrm{syn,k}(\mathbf{z}_a, \bar{x}_{\mathrm{k}a}, a)$ divided by the metabolic cost $E_\mathrm{k}$ of learning an average unit of skill.

We now re-derive the resulting proportionality between adult skill level and adult brain size. From (S7), if skill level growth has ceased at some (say adult) age $A$, then $B_\mathrm{syn,k} = 0$, which using (S5) yields $s_\mathrm{k} B_\mathrm{rest,b} = x_{\mathrm{k}A} B_\mathrm{k}$. If additionally brain growth has ceased at that age, then $g_{\mathrm{b}A}(\mathbf{z}_A, \bar{x}_{\mathrm{k}A}, A) - x_{\mathrm{b}A} = 0$, and so using (S6) we obtain $B_\mathrm{rest,b} = x_{\mathrm{b}A} B_\mathrm{b}$. Bringing these two expressions together, we obtain that when both skill level and brain size have ceased growing, we have that $s_\mathrm{k} x_{\mathrm{b}A} B_\mathrm{b} = x_{\mathrm{k}A} B_\mathrm{k}$. Solving for skill level yields the proportionality between adult skill level and adult brain size:

$$x_{\mathrm{k}A} = \frac{B_\mathrm{b}}{B_\mathrm{k}} s_\mathrm{k} x_{\mathrm{b}A}. \tag{S8}$$

Note this equation only depends on the assumptions made in the metabolic part, so is relatively general.

#### S3.2 Behavioural part

González-Forero et al. (2017) and González-Forero and Gardner (2018) make phenomenological considerations to obtain an expression for EEE as follows. At each age, a mutant faces energy extraction challenges of type $j \in \{1,2,3,4\}$ with probability $P_j$ (with $\sum_{j=1}^{4} P_j = 1$), where challenge types are:

Ecological if $j = 1$
Cooperative if $j = 2$
Between-individual competitive if $j = 3$
Between-group competitive if $j = 4$.

A mutant's energy extraction efficiency when facing type-$j$ challenges is $e_j(x_{\mathrm{k}a}, \bar{x}_{\mathrm{k}a}, a)$. Thus, a mutant's EEE is

$$e(x_{\mathrm{k}a}, \bar{x}_{\mathrm{k}a}, a) = \sum_{j=1}^{4} P_j e_j(x_{\mathrm{k}a}, \bar{x}_{\mathrm{k}a}, a).$$

A mutant succeeds at type-$j$ challenges a fraction $S_j(x_{\mathrm{k}a}, \bar{x}_{\mathrm{k}a})$ of the time. The EEE from maternal provisioning in case of individual failure at energy extraction at age $a$ is $\varphi(a)$. Hence, we let

$$e_j(x_{\mathrm{k}a}, \bar{x}_{\mathrm{k}a}, a) = S_j(x_{\mathrm{k}a}, \bar{x}_{\mathrm{k}a}) + [1 - S_j(x_{\mathrm{k}a}, \bar{x}_{\mathrm{k}a})]\varphi(a). \tag{S9}$$

We let the success proportion for type-$j$ challenges be

$$S_j(x_{\mathrm{k}a}, \bar{x}_{\mathrm{k}a}) = \frac{c_j(h_j(x_{\mathrm{k}a}, \bar{x}_{\mathrm{k}a}))}{c_j(h_j(x_{\mathrm{k}a}, \bar{x}_{\mathrm{k}a})) + d_j(\bar{x}_{\mathrm{k}a})}, \tag{S10}$$

where $c_j(h_j(x_{\mathrm{k}a}, \bar{x}_{\mathrm{k}a}))$ measures the individual's "competence" at type-$j$ challenges and $d_j(\bar{x}_{\mathrm{k}a})$ measures the type-$j$ challenge "difficulty" at age $a$. We let the competence take the same functional form across challenge types, so we write $c_j(h_j(x_{\mathrm{k}a}, \bar{x}_{\mathrm{k}a})) = c(h_j(x_{\mathrm{k}a}, \bar{x}_{\mathrm{k}a}))$, where the competence function takes one of two forms:

$$c(h_j(x_{\mathrm{k}a}, \bar{x}_{\mathrm{k}a})) = \begin{cases} [h_j(x_{\mathrm{k}a}, \bar{x}_{\mathrm{k}a})]^{\gamma} & \text{with power competence} \\ \exp(\gamma h_j(x_{\mathrm{k}a}, \bar{x}_{\mathrm{k}a})) & \text{with exponential competence.} \end{cases}$$

The function $h_j(x_{\mathrm{k}a}, \bar{x}_{\mathrm{k}a})$ describes how the skills of cooperating partners interact to yield a "joint" competence $c(h_j(x_{\mathrm{k}a}, \bar{x}_{\mathrm{k}a}))$. We consider the following forms

$$h_j(x_{\mathrm{k}a}, \bar{x}_{\mathrm{k}a}) = \begin{cases} x_{\mathrm{k}a} & \text{if } j \in \{1,3\} \\ x_{\mathrm{k}a} + \bar{x}_{\mathrm{k}a} & \text{if } j \in \{2,4\} \text{ with additive cooperation} \\ x_{\mathrm{k}a}\bar{x}_{\mathrm{k}a} & \text{if } j \in \{2,4\} \text{ with multiplicative cooperation} \\ (x_{\mathrm{k}a}\bar{x}_{\mathrm{k}a})^{1/2} & \text{if } j \in \{2,4\} \text{ with sub-multiplicative cooperation.} \end{cases}$$

We let environmental difficulty be

$$d_j(\bar{x}_{\mathrm{k}a}) = \begin{cases} \alpha & \text{if } j \in \{1,2\} \\ c(h_j(\bar{x}_{\mathrm{k}a}, \bar{x}_{\mathrm{k}a})) & \text{if } j \in \{3,4\}, \end{cases}$$

where $\alpha$ is a fixed parameter measuring the difficulty of ecological challenges. Finally, we let EEE from maternal provisioning be $\varphi(a) = \varphi_1 \exp(-\varphi_{\mathrm{r}}(a-1))$, with parameters $\varphi_1 \in [0,1]$ and $\varphi_{\mathrm{r}} \geq 0$.

##### S3.3 Summary

The mutant phenotype at age $a+1$ is thus given by the recurrence

$$\mathbf{x}_{a+1} = \mathbf{g}_a(\mathbf{z}_a, \bar{x}_{\mathrm{k}a}, a) = \begin{pmatrix} g_{\mathrm{b}a}(\mathbf{z}_a, \bar{x}_{\mathrm{k}a}, a) & g_{\mathrm{r}a}(\mathbf{z}_a, \bar{x}_{\mathrm{k}a}, a) & g_{\mathrm{s}a}(\mathbf{z}_a, \bar{x}_{\mathrm{k}a}, a) & g_{\mathrm{k}a}(\mathbf{z}_a, \bar{x}_{\mathrm{k}a}, a) \end{pmatrix}, \tag{S11}$$

with initial condition $\mathbf{x}_1 = \bar{\mathbf{x}}_1$ fixed, where $\mathbf{z}_a = (\mathbf{x}_a; \mathbf{y}_a)$, and with the developmental maps given by Eqs. (S4) and (S7). Since development is social, evaluation of the developmental map at resident genotypic traits (i.e., $\mathbf{g}_a(\mathbf{x}_a, \bar{\mathbf{y}}_a, \bar{x}_{\mathrm{k}a}, a)$) yields the resident phenotype only if the phenotype is socio-devo stable, which is obtained via socio-devo stabilization dynamics (González-Forero, 2024b).

#### S4 Derivation of developmental sensitivities

##### S4.1 Developmental sensitivity to parameters in the brain model

We now derive these developmental sensitivities. As this involves differentiating (S4) and (S7), in this section we assume that the phenotypes do not shrink (i.e., $x_{i,a+1} \geq x_{ia}$ for all $i \in \{\mathrm{b,r,s,k}\}$ and all $a \in \{1, \dots N_{\mathrm{a}} - 1\}$), which is the case for the developmental dynamics we analyse.

The developmental sensitivity of the phenotype to parameters affecting development is

$$\frac{\mathrm{d}\mathbf{x}^{\intercal}}{\mathrm{d}\boldsymbol{\epsilon}} = \frac{\partial \mathbf{x}^{\intercal}}{\partial \boldsymbol{\epsilon}} \frac{\mathrm{d}\mathbf{x}^{\intercal}}{\mathrm{d}\mathbf{x}}, \tag{S12}$$

(from Layer 4, Eq. S3 of González-Forero, 2024b) where $\boldsymbol{\epsilon} = (\boldsymbol{\epsilon}_1;\ldots;\boldsymbol{\epsilon}_{N_\mathrm{a}})$ is the vector of parameters affecting development over life. This is because the parameters affecting development are constant, and can thus be taken as "environmental traits" sensu González-Forero (2024b) trivially satisfying the environmental constraint in Eq. 2 of González-Forero (2024b).

We now compute the direct effects on the phenotype of the parameters in the developmental map. To do this, we build the vector of parameters directly affecting development at age $a$:

$$\boldsymbol{\epsilon}_a = (K,\beta,B_\mathrm{b},B_\mathrm{r},B_\mathrm{s},B_\mathrm{k},E_\mathrm{b},E_\mathrm{r},E_\mathrm{s},E_\mathrm{k},s_\mathrm{k},\alpha,\gamma,\varphi_1,\varphi_\mathrm{r},P_1,P_2,P_3,P_4)^\intercal \in \mathbb{R}_+^{19\times 1}, \tag{S13}$$

where $N_\mathrm{e} = 19$ is the number of such parameters. The developmental map does not directly depend on other parameters of the brain model modulating survival and fertility (e.g., mortality $\mu$ and the fertility proportionality constant $\tilde{f}$), so these parameters have no direct phenotypic effects ($\partial\mathbf{x}^\intercal/\partial\mu = \mathbf{0}$ and $\partial\mathbf{x}^\intercal/\partial\tilde{f} = \mathbf{0}$), and consequently their total phenotypic effects are also zero ($\mathrm{d}\mathbf{x}^\intercal/\mathrm{d}\mu = \mathbf{0}$ and $\mathrm{d}\mathbf{x}^\intercal/\mathrm{d}\tilde{f} = \mathbf{0}$). Hence, we do not include these two parameters in $\boldsymbol{\epsilon}_a$.

The matrix of the direct effects of parameters (Layer 2, Eq. S2c of González-Forero, 2024b) is

$$\left.\frac{\partial\mathbf{x}^\intercal}{\partial\boldsymbol{\epsilon}}\right|_{\mathbf{y}=\bar{\mathbf{y}}} = \left.\begin{pmatrix} \frac{\partial\mathbf{x}_1^\intercal}{\partial\boldsymbol{\epsilon}_1} & \cdots & \frac{\partial\mathbf{x}_{N_\mathrm{a}}^\intercal}{\partial\boldsymbol{\epsilon}_1} \\ \vdots & \ddots & \vdots \\ \frac{\partial\mathbf{x}_1^\intercal}{\partial\boldsymbol{\epsilon}_{N_\mathrm{a}}} & \cdots & \frac{\partial\mathbf{x}_{N_\mathrm{a}}^\intercal}{\partial\boldsymbol{\epsilon}_{N_\mathrm{a}}} \end{pmatrix}\right|_{\mathbf{y}=\bar{\mathbf{y}}} = \left.\begin{pmatrix} \mathbf{0} & \frac{\partial\mathbf{x}_2^\intercal}{\partial\boldsymbol{\epsilon}_1} & \cdots & \mathbf{0} & \mathbf{0} \\ \mathbf{0} & \mathbf{0} & \cdots & \mathbf{0} & \mathbf{0} \\ \vdots & \vdots & \ddots & \vdots & \vdots \\ \mathbf{0} & \mathbf{0} & \cdots & \mathbf{0} & \frac{\partial\mathbf{x}_{N_\mathrm{a}}^\intercal}{\partial\boldsymbol{\epsilon}_{N_\mathrm{a}-1}} \\ \mathbf{0} & \mathbf{0} & \cdots & \mathbf{0} & \mathbf{0} \end{pmatrix}\right|_{\mathbf{y}=\bar{\mathbf{y}}} \in \mathbb{R}^{N_\mathrm{a}N_\mathrm{e}\times N_\mathrm{a}N_\mathrm{p}}, \tag{S14}$$

where for all $a \in \{1,\ldots,N_\mathrm{a}-1\}$ we have

$$\left.\frac{\partial\mathbf{x}_{a+1}^\intercal}{\partial\boldsymbol{\epsilon}_a}\right|_{\mathbf{y}=\bar{\mathbf{y}}} \equiv \left.\begin{pmatrix} \frac{\partial x_{\mathrm{b},a+1}}{\partial K} & \frac{\partial x_{\mathrm{r},a+1}}{\partial K} & \frac{\partial x_{\mathrm{s},a+1}}{\partial K} & \frac{\partial x_{\mathrm{k},a+1}}{\partial K} \\ \frac{\partial x_{\mathrm{b},a+1}}{\partial \beta} & \frac{\partial x_{\mathrm{r},a+1}}{\partial \beta} & \frac{\partial x_{\mathrm{s},a+1}}{\partial \beta} & \frac{\partial x_{\mathrm{k},a+1}}{\partial \beta} \\ \vdots & \vdots & \vdots & \vdots \\ \frac{\partial x_{\mathrm{b},a+1}}{\partial P_4} & \frac{\partial x_{\mathrm{r},a+1}}{\partial P_4} & \frac{\partial x_{\mathrm{s},a+1}}{\partial P_4} & \frac{\partial x_{\mathrm{k},a+1}}{\partial P_4} \end{pmatrix}\right|_{\mathbf{y}=\bar{\mathbf{y}}} \in \mathbb{R}^{N_\mathrm{e}\times N_\mathrm{p}}. \tag{S15}$$

So we only need to calculate the matrix $\partial\mathbf{x}_{a+1}^\intercal/\partial\boldsymbol{\epsilon}_a$ for $a \in \{1,\ldots,N_\mathrm{a}-1\}$.

Differentiating (S4) and (S7) shows that

$$\frac{\partial\mathbf{x}_{a+1}^\intercal}{\partial K} = \begin{pmatrix} \frac{q_{\mathrm{b}a}}{E_\mathrm{b}} & \frac{q_{\mathrm{r}a}}{E_\mathrm{r}} & \frac{q_{\mathrm{s}a}}{E_\mathrm{s}} & \frac{q_{\mathrm{b}a}}{E_\mathrm{k}}s_\mathrm{k} \end{pmatrix} e x_{\mathrm{B}a}^\beta. \tag{S16}$$

Similarly,

$$\frac{\partial\mathbf{x}_{a+1}^\intercal}{\partial \beta} = \begin{pmatrix} \frac{q_{\mathrm{b}a}}{E_\mathrm{b}} & \frac{q_{\mathrm{r}a}}{E_\mathrm{r}} & \frac{q_{\mathrm{s}a}}{E_\mathrm{s}} & \frac{q_{\mathrm{b}a}}{E_\mathrm{k}}s_\mathrm{k} \end{pmatrix} K e x_{\mathrm{B}a}^\beta \ln x_{\mathrm{B}a}. \tag{S17}$$

Letting $\mathbf{B} = (B_\mathrm{b},B_\mathrm{r},B_\mathrm{s},B_\mathrm{k})^\intercal$, differentiating (S4) and (S7) shows that

$$\frac{\partial\mathbf{x}_{a+1}^\intercal}{\partial \mathbf{B}} = \begin{pmatrix} -\frac{q_{\mathrm{b}a}}{E_\mathrm{b}}x_{\mathrm{b}a} & -\frac{q_{\mathrm{r}a}}{E_\mathrm{r}}x_{\mathrm{b}a} & -\frac{q_{\mathrm{s}a}}{E_\mathrm{s}}x_{\mathrm{b}a} & \frac{s_\mathrm{k}}{E_\mathrm{k}}x_{\mathrm{b}a}(1-q_{\mathrm{b}a}) \\ -\frac{q_{\mathrm{b}a}}{E_\mathrm{b}}x_{\mathrm{r}a} & -\frac{q_{\mathrm{r}a}}{E_\mathrm{r}}x_{\mathrm{r}a} & -\frac{q_{\mathrm{s}a}}{E_\mathrm{s}}x_{\mathrm{r}a} & -\frac{s_\mathrm{k}}{E_\mathrm{k}}x_{\mathrm{r}a}q_{\mathrm{b}a} \\ -\frac{q_{\mathrm{b}a}}{E_\mathrm{b}}x_{\mathrm{s}a} & -\frac{q_{\mathrm{r}a}}{E_\mathrm{r}}x_{\mathrm{s}a} & -\frac{q_{\mathrm{s}a}}{E_\mathrm{s}}x_{\mathrm{s}a} & -\frac{s_\mathrm{k}}{E_\mathrm{k}}x_{\mathrm{s}a}q_{\mathrm{b}a} \\ 0 & 0 & 0 & -\frac{1}{E_\mathrm{k}}x_{\mathrm{k}a} \end{pmatrix}. \tag{S18}$$

The negative entries give the direct phenotypic decrease due to the metabolic costs of maintenance. In turn, the possibly positive entry gives the direct skill increase due to the metabolic cost of brain maintenance. The zero entries arise because the metabolic cost of memory $B_\mathrm{k}$ does not directly affect tissue mass.

Similarly, letting $\mathbf{E} = (E_\mathrm{b}, E_\mathrm{r}, E_\mathrm{s}, E_\mathrm{k})^\intercal$, differentiating (S4) and (S7) shows that

$$\frac{\partial \mathbf{x}_{a+1}^\intercal}{\partial \mathbf{E}} = \begin{pmatrix} -\frac{q_{\mathrm{b}a}}{E_\mathrm{b}^2} B_\mathrm{syn} & 0 & 0 & \frac{s_\mathrm{k}}{E_\mathrm{k}} x_{\mathrm{b}a} \\ 0 & -\frac{q_{\mathrm{r}a}}{E_\mathrm{r}^2} B_\mathrm{syn} & 0 & 0 \\ 0 & 0 & -\frac{q_{\mathrm{s}a}}{E_\mathrm{s}^2} B_\mathrm{syn} & 0 \\ 0 & 0 & 0 & -\frac{1}{E_\mathrm{k}^2} B_\mathrm{syn,k} \end{pmatrix}. \tag{S19}$$

As before, the negative entries give the direct phenotypic decrease due to the metabolic costs of growth. In turn, the positive entry gives the direct skill increase due to the metabolic cost of brain growth. The zero entries arise because the metabolic costs of growth only directly affect the corresponding phenotype, except for the positive entry as the metabolic cost of brain growth affects the brain's energy budget for skills.

Differentiating (S4) and (S7) also shows that

$$\frac{\partial \mathbf{x}_{a+1}^\intercal}{\partial s_\mathrm{k}} = \begin{pmatrix} 0 & 0 & 0 & \frac{1}{E_\mathrm{k}} B_\mathrm{rest,b} \end{pmatrix}. \tag{S20}$$

The non-zero entry is the skill increase due to brain's energy allocation to skills. All other entries are zero as such allocation does not directly affect tissue mass. In turn, letting competence be exponential (as this is the functional form used in the *Homo sapiens* scenario of the main text),

$$\frac{\partial \mathbf{x}_{a+1}^\intercal}{\partial \alpha} = -\begin{pmatrix} \frac{q_{\mathrm{b}a}}{E_\mathrm{b}} & \frac{q_{\mathrm{r}a}}{E_\mathrm{r}} & \frac{q_{\mathrm{s}a}}{E_\mathrm{s}} & \frac{q_{\mathrm{b}a}}{E_\mathrm{k}} s_\mathrm{k} \end{pmatrix} K x_{\mathrm{B}a}^\beta (1-\varphi) \sum_{j\in\{1,2\}} P_j S_j \frac{1}{c_j + d_j}. \tag{S21}$$

As all entries are negative, these are the direct phenotypic decrease due to environmental difficulty. Similarly, letting competence be exponential,

$$\frac{\partial \mathbf{x}_{a+1}^\intercal}{\partial \gamma} = \begin{pmatrix} \frac{q_{\mathrm{b}a}}{E_\mathrm{b}} & \frac{q_{\mathrm{r}a}}{E_\mathrm{r}} & \frac{q_{\mathrm{s}a}}{E_\mathrm{s}} & \frac{q_{\mathrm{b}a}}{E_\mathrm{k}} s_\mathrm{k} \end{pmatrix} K x_{\mathrm{B}a}^\beta (1-\varphi) \sum_{j\in\{1,2\}} P_j S_j (1-S_j) h_j. \tag{S22}$$

As all entries are positive, these are the direct phenotypic increase due to skill effectiveness.

In turn, differentiating (S4) and (S7) yields

$$\frac{\partial \mathbf{x}_{a+1}^\intercal}{\partial \varphi_1} = \begin{pmatrix} \frac{q_{\mathrm{b}a}}{E_\mathrm{b}} & \frac{q_{\mathrm{r}a}}{E_\mathrm{r}} & \frac{q_{\mathrm{s}a}}{E_\mathrm{s}} & \frac{q_{\mathrm{b}a}}{E_\mathrm{k}} s_\mathrm{k} \end{pmatrix} K x_{\mathrm{B}a}^\beta \exp(-\varphi_\mathrm{r}(a-1)) \sum_{j=1}^{4} P_j (1-S_j). \tag{S23}$$

As the entries are positive, these are the phenotypic increase due to maternal care towards newborns. Similarly, differentiating (S4) and (S7) yields

$$\frac{\partial \mathbf{x}_{a+1}^\intercal}{\partial \varphi_\mathrm{r}} = -\begin{pmatrix} \frac{q_{\mathrm{b}a}}{E_\mathrm{b}} & \frac{q_{\mathrm{r}a}}{E_\mathrm{r}} & \frac{q_{\mathrm{s}a}}{E_\mathrm{s}} & \frac{q_{\mathrm{b}a}}{E_\mathrm{k}} s_\mathrm{k} \end{pmatrix} K x_{\mathrm{B}a}^\beta \varphi_1 (a-1) \varphi \sum_{j=1}^{4} P_j (1-S_j). \tag{S24}$$

As the entries are negative, these are the phenotypic decrease due to increasing the rate of decrease of maternal care with age.

Differentiating (S4) and (S7) also shows that

$$\frac{\partial \mathbf{x}_{a+1}^\intercal}{\partial P_j} = \begin{pmatrix} \frac{q_{\mathrm{b}a}}{E_\mathrm{b}} & \frac{q_{\mathrm{r}a}}{E_\mathrm{r}} & \frac{q_{\mathrm{s}a}}{E_\mathrm{s}} & \frac{q_{\mathrm{b}a}}{E_\mathrm{k}} s_\mathrm{k} \end{pmatrix} K e_j x_{\mathrm{B}a}^\beta. \tag{S25}$$

As all entries are positive, perturbation of all challenge types causes direct phenotypic increases, without considering that $\sum_{j=1}^4 P_j = 1$.

#### S4.2 Developmental sensitivity to developmental history

The text of this section is taken from the Supplementary Information of González-Forero (2024a) and is repeated here for the reader's convenience.

The sensitivity of the phenotype to phenotypic change is given by the developmental feedback matrix

$$\frac{\mathrm{d}\mathbf{x}^\intercal}{\mathrm{d}\mathbf{x}} = \left(2\mathbf{I} - \frac{\partial \mathbf{x}^\intercal}{\partial \mathbf{x}}\right)^{-1}. \tag{S26}$$

We now compute the direct effects of the phenotype on the phenotype. This is given by the matrix quantifying the direct developmental bias of the phenotype from the phenotype, which is

$$\left.\frac{\partial \mathbf{x}^\intercal}{\partial \mathbf{x}}\right|_{\mathbf{y}=\bar{\mathbf{y}}} \equiv \left.\begin{pmatrix} \frac{\partial \mathbf{x}_1^\intercal}{\partial \mathbf{x}_1} & \cdots & \frac{\partial \mathbf{x}_{N_\mathrm{a}}^\intercal}{\partial \mathbf{x}_1} \\ \vdots & \ddots & \vdots \\ \frac{\partial \mathbf{x}_1^\intercal}{\partial \mathbf{x}_{N_\mathrm{a}}} & \cdots & \frac{\partial \mathbf{x}_{N_\mathrm{a}}^\intercal}{\partial \mathbf{x}_{N_\mathrm{a}}} \end{pmatrix}\right|_{\mathbf{y}=\bar{\mathbf{y}}} = \left.\begin{pmatrix} \mathbf{I} & \frac{\partial \mathbf{x}_2^\intercal}{\partial \mathbf{x}_1} & \cdots & \mathbf{0} & \mathbf{0} \\ \mathbf{0} & \mathbf{I} & \cdots & \mathbf{0} & \mathbf{0} \\ \vdots & \vdots & \ddots & \vdots & \vdots \\ \mathbf{0} & \mathbf{0} & \cdots & \mathbf{I} & \frac{\partial \mathbf{x}_{N_\mathrm{a}}^\intercal}{\partial \mathbf{x}_{N_\mathrm{a}-1}} \\ \mathbf{0} & \mathbf{0} & \cdots & \mathbf{0} & \mathbf{I} \end{pmatrix}\right|_{\mathbf{y}=\bar{\mathbf{y}}} \in \mathbb{R}^{N_\mathrm{a}N_\mathrm{p}\times N_\mathrm{a}N_\mathrm{p}} \tag{S27}$$

(Layer 2, Eq. S2a of ref. González-Forero, 2024b), where for all $a \in \{1,\ldots,N_\mathrm{a}-1\}$

$$\left.\frac{\partial \mathbf{x}_{a+1}^\intercal}{\partial \mathbf{x}_a}\right|_{\mathbf{y}=\bar{\mathbf{y}}} \equiv \left.\begin{pmatrix} \frac{\partial x_{\mathrm{b},a+1}}{\partial x_{\mathrm{b}a}} & \frac{\partial x_{\mathrm{r},a+1}}{\partial x_{\mathrm{b}a}} & \frac{\partial x_{\mathrm{s},a+1}}{\partial x_{\mathrm{b}a}} & \frac{\partial x_{\mathrm{k},a+1}}{\partial x_{\mathrm{b}a}} \\ \frac{\partial x_{\mathrm{b},a+1}}{\partial x_{\mathrm{r}a}} & \frac{\partial x_{\mathrm{r},a+1}}{\partial x_{\mathrm{r}a}} & \frac{\partial x_{\mathrm{s},a+1}}{\partial x_{\mathrm{r}a}} & \frac{\partial x_{\mathrm{k},a+1}}{\partial x_{\mathrm{r}a}} \\ \frac{\partial x_{\mathrm{b},a+1}}{\partial x_{\mathrm{s}a}} & \frac{\partial x_{\mathrm{r},a+1}}{\partial x_{\mathrm{s}a}} & \frac{\partial x_{\mathrm{s},a+1}}{\partial x_{\mathrm{s}a}} & \frac{\partial x_{\mathrm{k},a+1}}{\partial x_{\mathrm{s}a}} \\ \frac{\partial x_{\mathrm{b},a+1}}{\partial x_{\mathrm{k}a}} & \frac{\partial x_{\mathrm{r},a+1}}{\partial x_{\mathrm{k}a}} & \frac{\partial x_{\mathrm{s},a+1}}{\partial x_{\mathrm{k}a}} & \frac{\partial x_{\mathrm{k},a+1}}{\partial x_{\mathrm{k}a}} \end{pmatrix}\right|_{\mathbf{y}=\bar{\mathbf{y}}} \in \mathbb{R}^{N_\mathrm{p}\times N_\mathrm{p}} \tag{S28}$$

(see also Eqs. 18-19 of ref. González-Forero, 2024b). So we only need to calculate the matrix $\partial \mathbf{x}_{a+1}^\intercal/\partial \mathbf{x}_a$ for $a \in \{1,\ldots,N_\mathrm{a}-1\}$.

From Eq. (S4), the direct developmental bias of the mass of tissue $i \in \{\mathrm{b},\mathrm{r},\mathrm{s}\}$ from itself is

$$\frac{\partial x_{i,a+1}}{\partial x_{ia}} = \begin{cases} 1 + \frac{q_{ia}(\mathbf{y}_a)}{E_i}\frac{\partial B_\mathrm{syn}(\mathbf{x}_a,\bar{x}_{\mathrm{k}a},a)}{\partial x_{ia}} & \text{if } x_{ia} + \frac{q_{ia}(\mathbf{y}_a)}{E_i}B_\mathrm{syn}(\mathbf{x}_a,\bar{x}_{\mathrm{k}a},a) \geq 0 \\ 0 & \text{otherwise.} \end{cases} \tag{S29a}$$

Also from Eq. (S4), the direct developmental bias of the mass of tissue $j \in \{\mathrm{b},\mathrm{r},\mathrm{s}\}$ from the phenotypic trait $i \in \{\mathrm{b},\mathrm{r},\mathrm{s},\mathrm{k}\}$ with $i \neq j$ is

$$\frac{\partial x_{j,a+1}}{\partial x_{ia}} = \begin{cases} \frac{q_{ja}(\mathbf{y}_a)}{E_j}\frac{\partial B_\mathrm{syn}(\mathbf{x}_a,\bar{x}_{\mathrm{k}a},a)}{\partial x_{ia}} & \text{if } x_{ja} + \frac{q_{ja}(\mathbf{y}_a)}{E_j}B_\mathrm{syn}(\mathbf{x}_a,\bar{x}_{\mathrm{k}a},a) \geq 0 \\ 0 & \text{otherwise.} \end{cases} \tag{S29b}$$

Similarly, from Eq. (S7), the direct developmental bias of skill level from itself is

$$\frac{\partial x_{\mathrm{k},a+1}}{\partial x_{\mathrm{k}a}} = \begin{cases} 1 + \frac{1}{E_\mathrm{k}}\frac{\partial B_\mathrm{syn,k}(\mathbf{z}_a,\bar{x}_{\mathrm{k}a},a)}{\partial x_{\mathrm{k}a}} & \text{if } x_{\mathrm{k}a} + \frac{1}{E_\mathrm{k}}B_\mathrm{syn,k}(\mathbf{z}_a,\bar{x}_{\mathrm{k}a},a) \geq 0 \\ 0 & \text{otherwise.} \end{cases}$$

Also from Eq. (S7), the direct developmental bias of skill level from the mass of tissue $i \in \{\mathrm{b},\mathrm{r},\mathrm{s}\}$ is

$$\frac{\partial x_{\mathrm{k},a+1}}{\partial x_{ia}} = \begin{cases} \frac{1}{E_\mathrm{k}}\frac{\partial B_\mathrm{syn,k}(\mathbf{z}_a,\bar{x}_{\mathrm{k}a},a)}{\partial x_{ia}} & \text{if } x_{\mathrm{k}a} + \frac{1}{E_\mathrm{k}}B_\mathrm{syn,k}(\mathbf{z}_a,\bar{x}_{\mathrm{k}a},a) \geq 0 \\ 0 & \text{otherwise.} \end{cases}$$

Thus, if $x_{ja} + \frac{q_{ja}(\mathbf{y}_a)}{E_j}B_\mathrm{syn}(\mathbf{x}_a,\bar{x}_{\mathrm{k}a},a) \geq 0$ for $j \in \{\mathrm{b},\mathrm{r},\mathrm{s}\}$ and $x_{\mathrm{k}a} + \frac{1}{E_\mathrm{k}}B_\mathrm{syn,k}(\mathbf{z}_a,\bar{x}_{\mathrm{k}a},a) \geq 0$, using Eq. (S28), the age-specific matrix of the direct developmental bias of the phenotype from the phenotype is

$$\frac{\partial \mathbf{x}_{a+1}^{\mathsf{T}}}{\partial \mathbf{x}_a} = \begin{pmatrix} 1+\frac{q_{\mathrm{b}a}}{E_{\mathrm{b}}}\frac{\partial B_{\mathrm{syn}}}{\partial x_{\mathrm{b}a}} & \frac{q_{\mathrm{r}a}}{E_{\mathrm{r}}}\frac{\partial B_{\mathrm{syn}}}{\partial x_{\mathrm{b}a}} & \frac{q_{\mathrm{s}a}}{E_{\mathrm{s}}}\frac{\partial B_{\mathrm{syn}}}{\partial x_{\mathrm{b}a}} & \frac{1}{E_{\mathrm{k}}}\frac{\partial B_{\mathrm{syn,k}}}{\partial x_{\mathrm{b}a}} \\ \frac{q_{\mathrm{b}a}}{E_{\mathrm{b}}}\frac{\partial B_{\mathrm{syn}}}{\partial x_{\mathrm{r}a}} & 1+\frac{q_{\mathrm{r}a}}{E_{\mathrm{r}}}\frac{\partial B_{\mathrm{syn}}}{\partial x_{\mathrm{r}a}} & \frac{q_{\mathrm{s}a}}{E_{\mathrm{s}}}\frac{\partial B_{\mathrm{syn}}}{\partial x_{\mathrm{r}a}} & \frac{1}{E_{\mathrm{k}}}\frac{\partial B_{\mathrm{syn,k}}}{\partial x_{\mathrm{r}a}} \\ \frac{q_{\mathrm{b}a}}{E_{\mathrm{b}}}\frac{\partial B_{\mathrm{syn}}}{\partial x_{\mathrm{s}a}} & \frac{q_{\mathrm{r}a}}{E_{\mathrm{r}}}\frac{\partial B_{\mathrm{syn}}}{\partial x_{\mathrm{s}a}} & 1+\frac{q_{\mathrm{s}a}}{E_{\mathrm{s}}}\frac{\partial B_{\mathrm{syn}}}{\partial x_{\mathrm{s}a}} & \frac{1}{E_{\mathrm{k}}}\frac{\partial B_{\mathrm{syn,k}}}{\partial x_{\mathrm{s}a}} \\ \frac{q_{\mathrm{b}a}}{E_{\mathrm{b}}}\frac{\partial B_{\mathrm{syn}}}{\partial x_{\mathrm{k}a}} & \frac{q_{\mathrm{r}a}}{E_{\mathrm{r}}}\frac{\partial B_{\mathrm{syn}}}{\partial x_{\mathrm{k}a}} & \frac{q_{\mathrm{s}a}}{E_{\mathrm{s}}}\frac{\partial B_{\mathrm{syn}}}{\partial x_{\mathrm{k}a}} & 1+\frac{1}{E_{\mathrm{k}}}\frac{\partial B_{\mathrm{syn,k}}}{\partial x_{\mathrm{k}a}} \end{pmatrix}, \tag{S30}$$

where the direct effect on the growth metabolic rate of perturbing the phenotype $i$ at age $a$ is

$$\frac{\partial B_{\mathrm{syn}}}{\partial x_{ia}} = \begin{cases} Ke(x_{\mathrm{k}a},\bar{x}_{\mathrm{k}a},a)\beta x_{\mathrm{B}}^{\beta-1} - B_i & \text{if } i \in \{\mathrm{b},\mathrm{r},\mathrm{s}\} \\ K\frac{\partial e}{\partial x_{\mathrm{k}a}} x_{\mathrm{B}}^{\beta} & \text{if } i = \mathrm{k}, \end{cases}$$

and the direct effect on the learning metabolic rate of perturbing the phenotype $i$ at age $a$ is

$$\frac{\partial B_{\mathrm{syn,k}}}{\partial x_{ia}} = \begin{cases} s_{\mathrm{k}}\frac{\partial B_{\mathrm{rest,b}}}{\partial x_{ia}} & \text{if } i \in \{\mathrm{b},\mathrm{r},\mathrm{s}\} \\ s_{\mathrm{k}}\frac{\partial B_{\mathrm{rest,b}}}{\partial x_{\mathrm{k}a}} - B_{\mathrm{k}} & \text{if } i = \mathrm{k}. \end{cases}$$

In turn, noting that $g_{\mathrm{b}a} = x_{\mathrm{b},a+1}$, the direct effect on the brain metabolic rate of the phenotype $i$ at age $a$ is

$$\frac{\partial B_{\mathrm{rest,b}}}{\partial x_{ia}} = \begin{cases} B_{\mathrm{b}} + \left(\frac{\partial x_{\mathrm{b},a+1}}{\partial x_{\mathrm{b}a}} - 1\right) E_{\mathrm{b}} & \text{if } i = \mathrm{b} \\ \frac{\partial x_{\mathrm{b},a+1}}{\partial x_{ia}} E_{\mathrm{b}} & \text{if } i \in \{\mathrm{r},\mathrm{s},\mathrm{k}\}, \end{cases}$$

where $\partial x_{\mathrm{b},a+1}/\partial x_{ia}$ for $i \in \{\mathrm{b},\mathrm{r},\mathrm{s},\mathrm{k}\}$ is given by Eq. (S29).

Now, the direct effect on the EEE of the individual's skill level at age $a$ is

$$\frac{\partial e}{\partial x_{\mathrm{k}a}} = \sum_{j=1}^{4} P_j \frac{\partial e_j}{\partial x_{\mathrm{k}a}},$$

where the direct effect on the EEE for challenge type $j$ of the individual's skill level at age $a$ is

$$\frac{\partial e_j}{\partial x_{\mathrm{k}a}} = [1-\varphi(a)]\frac{\partial S_j}{\partial x_{\mathrm{k}a}},$$

and the direct effect on the success proportion for challenges of type $j$ from perturbing the individual's skill level at age $a$ is

$$\frac{\partial S_j}{\partial x_{\mathrm{k}a}} = (1-S_j)\frac{1}{c(h_j(x_{\mathrm{k}a},\bar{x}_{\mathrm{k}a})) + d_j(\bar{x}_{\mathrm{k}a})}\frac{\mathrm{d}c}{\mathrm{d}h_j}\frac{\partial h_j}{\partial x_{\mathrm{k}a}}.$$

In turn, the direct effect on the individual's competence from perturbing her skill level at age $a$ is

$$\frac{\mathrm{d}c}{\mathrm{d}h_j} = \begin{cases} \gamma h_j^{\gamma-1} & \text{with power competence} \\ \gamma \exp(\gamma h_j) & \text{with exponential competence,} \end{cases}$$

and the direct effect on the joint action of skills from perturbing the individual's skill level at age $a$ is

$$\frac{\partial h_j}{\partial x_{\mathrm{k}a}} = \begin{cases} 1 & \text{if } j \in \{1,3\} \\ 1 & \text{if } j \in \{2,4\} \text{ with additive cooperation} \\ \bar{x}_{\mathrm{k}a} & \text{if } j \in \{2,4\} \text{ with multiplicative cooperation} \\ \frac{1}{2}\left(\frac{\bar{x}_{\mathrm{k}a}}{x_{\mathrm{k}a}}\right)^{1/2} & \text{if } j \in \{2,4\} \text{ with sub-multiplicative cooperation.} \end{cases}$$

#### S4.3 Developmental sensitivity to the genotype

The text of this section is taken from the Supplementary Information of González-Forero (2024a) and is repeated here for the reader's convenience.

The sensitivity of the phenotype to genotypic change is given by

$$\frac{\mathrm{d}\mathbf{x}^\intercal}{\mathrm{d}\mathbf{y}} = \frac{\partial \mathbf{x}^\intercal}{\partial \mathbf{y}} \frac{\mathrm{d}\mathbf{x}^\intercal}{\mathrm{d}\mathbf{x}}. \tag{S31}$$

We now compute the direct effects of the genotype on the phenotype. This is given by the matrix quantifying the direct developmental bias of the phenotype from the genotype, which is

$$\left.\frac{\partial \mathbf{x}^\intercal}{\partial \mathbf{y}}\right|_{\mathbf{y}=\bar{\mathbf{y}}} \equiv \left.\begin{pmatrix} \frac{\partial \mathbf{x}_1^\intercal}{\partial \mathbf{y}_1} & \cdots & \frac{\partial \mathbf{x}_{N_\mathrm{a}}^\intercal}{\partial \mathbf{y}_1} \\ \vdots & \ddots & \vdots \\ \frac{\partial \mathbf{x}_1^\intercal}{\partial \mathbf{y}_{N_\mathrm{a}}} & \cdots & \frac{\partial \mathbf{x}_{N_\mathrm{a}}^\intercal}{\partial \mathbf{y}_{N_\mathrm{a}}} \end{pmatrix}\right|_{\mathbf{y}=\bar{\mathbf{y}}} = \left.\begin{pmatrix} \mathbf{0} & \frac{\partial \mathbf{x}_2^\intercal}{\partial \mathbf{y}_1} & \cdots & \mathbf{0} & \mathbf{0} \\ \mathbf{0} & \mathbf{0} & \cdots & \mathbf{0} & \mathbf{0} \\ \vdots & \vdots & \ddots & \vdots & \vdots \\ \mathbf{0} & \mathbf{0} & \cdots & \mathbf{0} & \frac{\partial \mathbf{x}_{N_\mathrm{a}}^\intercal}{\partial \mathbf{y}_{N_\mathrm{a}-1}} \\ \mathbf{0} & \mathbf{0} & \cdots & \mathbf{0} & \mathbf{0} \end{pmatrix}\right|_{\mathbf{y}=\bar{\mathbf{y}}} \in \mathbb{R}^{N_\mathrm{a}N_\mathrm{g} \times N_\mathrm{a}N_\mathrm{p}} \tag{S32}$$

(Layer 2, Eq. S2b of ref. González-Forero, 2024b), where for all $a \in \{1, \ldots, N_\mathrm{a}-1\}$

$$\left.\frac{\partial \mathbf{x}_{a+1}^\intercal}{\partial \mathbf{y}_a}\right|_{\mathbf{y}=\bar{\mathbf{y}}} \equiv \left.\begin{pmatrix} \frac{\partial x_{\mathrm{b},a+1}}{\partial y_{\mathrm{b}a}} & \frac{\partial x_{\mathrm{r},a+1}}{\partial y_{\mathrm{b}a}} & \frac{\partial x_{\mathrm{s},a+1}}{\partial y_{\mathrm{b}a}} & \frac{\partial x_{\mathrm{k},a+1}}{\partial y_{\mathrm{b}a}} \\ \frac{\partial x_{\mathrm{b},a+1}}{\partial y_{\mathrm{r}a}} & \frac{\partial x_{\mathrm{r},a+1}}{\partial y_{\mathrm{r}a}} & \frac{\partial x_{\mathrm{s},a+1}}{\partial y_{\mathrm{r}a}} & \frac{\partial x_{\mathrm{k},a+1}}{\partial y_{\mathrm{r}a}} \\ \frac{\partial x_{\mathrm{b},a+1}}{\partial y_{\mathrm{s}a}} & \frac{\partial x_{\mathrm{r},a+1}}{\partial y_{\mathrm{s}a}} & \frac{\partial x_{\mathrm{s},a+1}}{\partial y_{\mathrm{s}a}} & \frac{\partial x_{\mathrm{k},a+1}}{\partial y_{\mathrm{s}a}} \end{pmatrix}\right|_{\mathbf{y}=\bar{\mathbf{y}}} \in \mathbb{R}^{N_\mathrm{g} \times N_\mathrm{p}}. \tag{S33}$$

So we only need to calculate the matrix $\partial \mathbf{x}_{a+1}^\intercal / \partial \mathbf{y}_a$ for $a \in \{1, \ldots, N_\mathrm{a}-1\}$.

From Eq. (S4), the direct developmental bias of mass of tissue $j \in \{\mathrm{b}, \mathrm{r}, \mathrm{s}\}$ from genotypic trait $i \in \{\mathrm{b}, \mathrm{r}, \mathrm{s}\}$ is

$$\frac{\partial x_{j,a+1}}{\partial y_{ia}} = \begin{cases} \frac{\partial q_{ja}(\mathbf{y}_a)}{\partial y_{ia}} \frac{1}{E_j} B_\mathrm{syn}(\mathbf{x}_a, \bar{x}_{\mathrm{k}a}, a) & \text{if } x_{ja} + \frac{q_{ja}(\mathbf{y}_a)}{E_j} B_\mathrm{syn}(\mathbf{x}_a, \bar{x}_{\mathrm{k}a}, a) \geq 0 \\ 0 & \text{otherwise,} \end{cases} \tag{S34}$$

where the direct effect on the growth allocation of growth efforts is

$$\frac{\partial q_{ja}(\mathbf{y}_a)}{\partial y_{ia}} = \begin{cases} q_{ja}(\mathbf{y}_a)[1 - q_{ja}(\mathbf{y}_a)] & \text{if } i = j \\ -q_{ja}(\mathbf{y}_a) q_{ia}(\mathbf{y}_a) & \text{if } i \neq j. \end{cases}$$

Also, from Eq. (S7), the direct developmental bias of skill level from genotypic trait $i \in \{\mathrm{b}, \mathrm{r}, \mathrm{s}\}$ is

$$\frac{\partial x_{\mathrm{k},a+1}}{\partial y_{ia}} = \begin{cases} \frac{1}{E_\mathrm{k}} \frac{\partial B_\mathrm{syn,k}(\mathbf{z}_a, \bar{x}_{\mathrm{k}a}, a)}{\partial y_{ia}} & \text{if } x_{\mathrm{k}a} + \frac{1}{E_\mathrm{k}} B_\mathrm{syn,k}(\mathbf{z}_a, \bar{x}_{\mathrm{k}a}, a) \geq 0 \\ 0 & \text{otherwise,} \end{cases}$$

where the direct effect on the learning metabolic rate of the growth efforts is

$$\frac{\partial B_\mathrm{syn,k}(\mathbf{z}_a, \bar{x}_{\mathrm{k}a}, a)}{\partial y_{ia}} = s_\mathrm{k} \frac{\partial B_\mathrm{rest,b}(\mathbf{z}_a, \bar{x}_{\mathrm{k}a}, a)}{\partial y_{ia}},$$

and the direct effect on the brain metabolic rate of the growth efforts is

$$\frac{\partial B_\mathrm{rest,b}(\mathbf{z}_a, \bar{x}_{\mathrm{k}a}, a)}{\partial y_{ia}} = \frac{\partial g_{\mathrm{b}a}(\mathbf{z}_a, \bar{x}_{\mathrm{k}a}, a)}{\partial y_{ia}} E_\mathrm{b} = \frac{\partial x_{\mathrm{b},a+1}}{\partial y_{ia}} E_\mathrm{b}. \tag{S35}$$

Thus, the direct developmental bias of skill level from genotypic trait $i \in \{\mathrm{b}, \mathrm{r}, \mathrm{s}\}$ reduces to

$$\frac{\partial x_{\mathrm{k},a+1}}{\partial y_{ia}} = \begin{cases} \frac{1}{E_\mathrm{k}} s_\mathrm{k} \frac{\partial q_{\mathrm{b}a}(\mathbf{y}_a)}{\partial y_{ia}} B_\mathrm{syn}(\mathbf{x}_a, \bar{x}_{\mathrm{k}a}, a) & \text{if } x_{\mathrm{k}a} + \frac{1}{E_\mathrm{k}} B_\mathrm{syn,k}(\mathbf{z}_a, \bar{x}_{\mathrm{k}a}, a) \geq 0 \text{ and } x_{\mathrm{b}a} + \frac{q_{\mathrm{b}a}(\mathbf{y}_a)}{E_\mathrm{b}} B_\mathrm{syn}(\mathbf{x}_a, \bar{x}_{\mathrm{k}a}, a) \geq 0 \\ 0 & \text{otherwise.} \end{cases}$$

Hence, if $x_{ja} + \dfrac{q_{ja}(\mathbf{y}_a)}{E_j} B_{\text{syn}}(\mathbf{x}_a, \bar{x}_{\text{k}a}, a) \geq 0$ for $j \in \{\text{b}, \text{r}, \text{s}\}$ and $x_{\text{k}a} + \dfrac{1}{E_\text{k}} B_{\text{syn,k}}(\mathbf{z}_a, \bar{x}_{\text{k}a}, a) \geq 0$, using Eq. (S28), the age-specific matrix of the direct developmental bias of the phenotype from the genotype is

$$\frac{\partial \mathbf{x}_{a+1}^{\intercal}}{\partial \mathbf{y}_a} = \begin{pmatrix} q_{\text{b}a}(1-q_{\text{b}a})\frac{1}{E_\text{b}} & -q_{\text{r}a}q_{\text{b}a}\frac{1}{E_\text{r}} & -q_{\text{s}a}q_{\text{b}a}\frac{1}{E_\text{s}} & q_{\text{b}a}(1-q_{\text{b}a})\frac{s_\text{k}}{E_\text{k}} \\ -q_{\text{b}a}q_{\text{r}a}\frac{1}{E_\text{b}} & q_{\text{r}a}(1-q_{\text{r}a})\frac{1}{E_\text{r}} & -q_{\text{s}a}q_{\text{r}a}\frac{1}{E_\text{s}} & -q_{\text{b}a}q_{\text{r}a}\frac{s_\text{k}}{E_\text{k}} \\ -q_{\text{b}a}q_{\text{s}a}\frac{1}{E_\text{b}} & -q_{\text{r}a}q_{\text{s}a}\frac{1}{E_\text{r}} & q_{\text{s}a}(1-q_{\text{s}a})\frac{1}{E_\text{s}} & -q_{\text{b}a}q_{\text{s}a}\frac{s_\text{k}}{E_\text{k}} \end{pmatrix} B_{\text{syn}}. \tag{S36}$$

#### S4.4 Developmental sensitivity to social interactions

The text of this section is taken from the Supplementary Information of González-Forero (2024a) and is repeated here for the reader's convenience.

The sensitivity of the phenotype to social change is given by

$$\frac{\text{d}\mathbf{x}^{\intercal}}{\text{d}\bar{\mathbf{x}}} = \frac{\partial \mathbf{x}^{\intercal}}{\partial \bar{\mathbf{x}}} \frac{\text{d}\mathbf{x}^{\intercal}}{\text{d}\mathbf{x}}. \tag{S37}$$

We now compute the direct effects of the phenotype of social partners on the phenotype. This is given by the matrix quantifying the direct social developmental bias from the phenotype, which is

$$\left.\frac{\partial \mathbf{x}^{\intercal}}{\partial \bar{\mathbf{x}}}\right|_{\mathbf{y}=\bar{\mathbf{y}}} = \left.\begin{pmatrix} \frac{\partial \mathbf{x}_1^{\intercal}}{\partial \bar{\mathbf{x}}_1} & \cdots & \frac{\partial \mathbf{x}_{N_\text{a}}^{\intercal}}{\partial \bar{\mathbf{x}}_1} \\ \vdots & \ddots & \vdots \\ \frac{\partial \mathbf{x}_1^{\intercal}}{\partial \bar{\mathbf{x}}_{N_\text{a}}} & \cdots & \frac{\partial \mathbf{x}_{N_\text{a}}^{\intercal}}{\partial \bar{\mathbf{x}}_{N_\text{a}}} \end{pmatrix}\right|_{\mathbf{y}=\bar{\mathbf{y}}} = \left.\begin{pmatrix} \mathbf{0} & \frac{\partial \mathbf{x}_2^{\intercal}}{\partial \bar{\mathbf{x}}_1} & \cdots & \mathbf{0} & \mathbf{0} \\ \mathbf{0} & \mathbf{0} & \cdots & \mathbf{0} & \mathbf{0} \\ \vdots & \vdots & \ddots & \vdots & \vdots \\ \mathbf{0} & \mathbf{0} & \cdots & \mathbf{0} & \frac{\partial \mathbf{x}_{N_\text{a}}^{\intercal}}{\partial \bar{\mathbf{x}}_{N_\text{a}-1}} \\ \mathbf{0} & \mathbf{0} & \cdots & \mathbf{0} & \mathbf{0} \end{pmatrix}\right|_{\mathbf{y}=\bar{\mathbf{y}}} \in \mathbb{R}^{N_\text{a}N_\text{p} \times N_\text{a}N_\text{p}}$$

(Layer 2, Eq. S4 of ref. González-Forero, 2024b). From Eqs. (S4) and (S7), the only direct dependence of a mutant's phenotype on social partners' phenotype is via the skill level of social partners of her same age, so for all $a \in \{1, \ldots, N_\text{a} - 1\}$

$$\left.\frac{\partial \mathbf{x}_{a+1}^{\intercal}}{\partial \bar{\mathbf{x}}_a}\right|_{\mathbf{y}=\bar{\mathbf{y}}} \equiv \left.\begin{pmatrix} 0 & 0 & 0 & 0 \\ 0 & 0 & 0 & 0 \\ 0 & 0 & 0 & 0 \\ \frac{\partial x_{\text{b},a+1}}{\partial \bar{x}_{\text{k}a}} & \frac{\partial x_{\text{r},a+1}}{\partial \bar{x}_{\text{k}a}} & \frac{\partial x_{\text{s},a+1}}{\partial \bar{x}_{\text{k}a}} & \frac{\partial x_{\text{k},a+1}}{\partial \bar{x}_{\text{k}a}} \end{pmatrix}\right|_{\mathbf{y}=\bar{\mathbf{y}}} \in \mathbb{R}^{N_\text{p} \times N_\text{p}}. \tag{S38}$$

Hence, we only need to consider the direct social developmental bias from social partners' skill level.

Consequently, using Eq. (S4), the direct social developmental bias of the mass of tissue $j \in \{\text{b}, \text{r}, \text{s}\}$ from social partners' skill level is

$$\frac{\partial x_{j,a+1}}{\partial \bar{x}_{\text{k}a}} = \begin{cases} \frac{q_{ja}(\mathbf{y}_a)}{E_j} \frac{\partial B_{\text{syn}}(\mathbf{x}_a, \bar{x}_{\text{k}a}, a)}{\partial \bar{x}_{\text{k}a}} & \text{if } x_{ja} + \frac{q_{ja}(\mathbf{y}_a)}{E_j} B_{\text{syn}}(\mathbf{x}_a, \bar{x}_{\text{k}a}, a) \geq 0 \\ 0 & \text{otherwise,} \end{cases}$$

where the direct effect on the growth metabolic rate of perturbing a social partner's skill level at age $a$ is

$$\frac{\partial B_{\text{syn}}(\mathbf{x}_a, \bar{x}_{\text{k}a}, a)}{\partial \bar{x}_{\text{k}a}} = K \frac{\partial e(x_{\text{k}a}, \bar{x}_{\text{k}a}, a)}{\partial \bar{x}_{\text{k}a}} x_\text{B}^{\beta},$$

and the direct effect on the EEE of perturbing a social partner's skill level at age $a$ is

$$\frac{\partial e(x_{\text{k}a}, \bar{x}_{\text{k}a}, a)}{\partial \bar{x}_{\text{k}a}} = \sum_{j=1}^{4} P_j \frac{\partial e_j(x_{\text{k}a}, \bar{x}_{\text{k}a}, a)}{\partial \bar{x}_{\text{k}a}}.$$

The direct effect on the EEE for a challenge of type $j$ from perturbing a social partner's skill level at age $a$ is

$$\frac{\partial e_j}{\partial \bar{x}_{\mathrm{k}a}} = [1-\varphi(a)]\frac{\partial S_j}{\partial \bar{x}_{\mathrm{k}a}},$$

and the direct effect on the success proportion from perturbing a social partner's skill level at age $a$ is

$$\frac{\partial S_j}{\partial \bar{x}_{\mathrm{k}a}} = \frac{1}{c(h_j(x_{\mathrm{k}a},\bar{x}_{\mathrm{k}a}))+d_j(\bar{x}_{\mathrm{k}a})}\left[(1-S_j)\frac{\partial c}{\partial h_j}\frac{\partial h_j}{\partial \bar{x}_{\mathrm{k}a}} - S_j\frac{\mathrm{d}d_j}{\mathrm{d}\bar{x}_{\mathrm{k}a}}\right].$$

In turn, the direct effect of the social partner's skill level on the joint action of the skills is

$$\frac{\partial h_j}{\partial \bar{x}_{\mathrm{k}a}} = \begin{cases} 0 & \text{if } j\in\{1,3\} \\ 1 & \text{if } j\in\{2,4\} \text{ with additive cooperation} \\ x_{\mathrm{k}a} & \text{if } j\in\{2,4\} \text{ with multiplicative cooperation} \\ \dfrac{1}{2}\left(\dfrac{x_{\mathrm{k}a}}{\bar{x}_{\mathrm{k}a}}\right)^{1/2} & \text{if } j\in\{2,4\} \text{ with sub-multiplicative cooperation,} \end{cases}$$

and the direct (and total) effect of the social partner's skill level on the difficulty of challenges of type $j$ is

$$\frac{\mathrm{d}d_j}{\mathrm{d}\bar{x}_{\mathrm{k}a}} = \begin{cases} 0 & \text{if } j\in\{1,2\} \\ \dfrac{\mathrm{d}c}{\mathrm{d}h_j}\left(\dfrac{\partial h_j}{\partial x_{\mathrm{k}a}}+\dfrac{\partial h_j}{\partial \bar{x}_{\mathrm{k}a}}\right) & \text{if } j\in\{3,4\}. \end{cases}$$

Similarly, using Eq. (S7), the direct social developmental bias of skill level from social partners' skill level is

$$\frac{\partial x_{\mathrm{k},a+1}}{\partial \bar{x}_{\mathrm{k}a}} = \begin{cases} \dfrac{1}{E_\mathrm{k}}\dfrac{\partial B_{\mathrm{syn,k}}(\mathbf{z}_a,\bar{x}_{\mathrm{k}a},a)}{\partial \bar{x}_{\mathrm{k}a}} & \text{if } x_{\mathrm{k}a}+\dfrac{1}{E_\mathrm{k}}B_{\mathrm{syn,k}}(\mathbf{z}_a,\bar{x}_{\mathrm{k}a},a)\geq 0 \\ 0 & \text{otherwise,} \end{cases}$$

where the direct effect on the learning metabolic rate from the perturbation of a social partner's skill level at age $a$ is

$$\frac{\partial B_{\mathrm{syn,k}}(\mathbf{z}_a,\bar{x}_{\mathrm{k}a},a)}{\partial \bar{x}_{\mathrm{k}a}} = s_\mathrm{k}\frac{\partial B_{\mathrm{rest,b}}(\mathbf{z}_a,\bar{x}_{\mathrm{k}a},a)}{\partial \bar{x}_{\mathrm{k}a}},$$

and the direct effect on the brain metabolic rate from the perturbation of a social partner's skill level at age $a$ is

$$\frac{\partial B_{\mathrm{rest,b}}(\mathbf{z}_a,\bar{x}_{\mathrm{k}a},a)}{\partial \bar{x}_{\mathrm{k}a}} = \frac{\partial g_{\mathrm{b}a}(\mathbf{z}_a,\bar{x}_{\mathrm{k}a},a)}{\partial \bar{x}_{\mathrm{k}a}}E_\mathrm{b} = \frac{\partial x_{\mathrm{b},a+1}}{\partial \bar{x}_{\mathrm{k}a}}E_\mathrm{b}.$$

Hence, if $x_{ja}+\dfrac{q_{ja}(\mathbf{y}_a)}{E_j}B_{\mathrm{syn}}(\mathbf{x}_a,\bar{x}_{\mathrm{k}a},a)\geq 0$ for $j\in\{\mathrm{b,r,s}\}$ and $x_{\mathrm{k}a}+\dfrac{1}{E_\mathrm{k}}B_{\mathrm{syn,k}}(\mathbf{z}_a,\bar{x}_{\mathrm{k}a},a)\geq 0$, the direct social developmental bias matrix of the phenotype from the phenotype is

$$\frac{\partial \mathbf{x}_{a+1}^\intercal}{\partial \bar{\mathbf{x}}_a} = \begin{pmatrix} 0 & 0 & 0 & 0 \\ 0 & 0 & 0 & 0 \\ 0 & 0 & 0 & 0 \\ \dfrac{q_{\mathrm{b}a}}{E_\mathrm{b}} & \dfrac{q_{\mathrm{r}a}}{E_\mathrm{r}} & \dfrac{q_{\mathrm{s}a}}{E_\mathrm{s}} & s_\mathrm{k}\dfrac{q_{\mathrm{b}a}}{E_\mathrm{k}} \end{pmatrix}\frac{\partial B_{\mathrm{syn}}}{\partial \bar{x}_{\mathrm{k}a}}.$$

### S5 Heritability

Now we describe how we calculate heritability. We introduce genotypic variation by letting genotypic traits be given by $y_{ij}^{(r)} = \bar{y}_{ij}+\delta_{yij}^{(r)}$, where $\bar{y}_{ij}$ are the genotypic traits evolved by the brain model under the *H. sapiens* scenario and $\delta_{yij}^{(r)}$ is a value drawn from the normal distribution with mean zero and variance $\sigma_\mathrm{g}^2$ (we draw $N = 10000$ independent values of $\delta_{yij}^{(r)}$, with $r\in\{1,\ldots,N\}$). We introduce environmental variation by letting the environmental difficulty faced at age $j$ be given by $\alpha_j^{(r)} = \bar{\alpha}+\delta_{\alpha j}^{(r)}$, where $\bar{\alpha}=\alpha$ is the environmental difficulty used by the brain model under the *H. sapiens* scenario and $\delta^{(r)}_{\alpha j}$ is a value drawn from the normal distribution with mean zero and variance $\sigma^2_\mathrm{e}$ (we draw $N = 10000$ independent values of $\delta^{(r)}_{\alpha j}$). Using the vector notation defined in the main text, we then obtain a vector $\mathbf{y}^{(r)}$ of genotypic traits and a vector of environmental difficulty $\boldsymbol{\alpha}^{(r)}$ across life for replica $r$. We then compute the developed phenotypes $\mathbf{x}^{(r)}$ by solving the developmental dynamics in Eqs. (S4) and (S7) evaluated at the genotypic traits $\mathbf{y}^{(r)}$ and environment $\boldsymbol{\alpha}^{(r)}$.

We compute heritability using its standard definition as follows. We form the matrix of predictor variables $\mathbf{X} \in \mathbb{R}^{N\times(N_\mathrm{g}N_\mathrm{a}+1)}$, whose first column is filled with ones and the $r$-th row in the remainder columns is the vector of genotypic differences $\mathrm{vec}(\boldsymbol{\delta}^{(r)})$ (the first column of ones is to estimate the intercepts, which yields a higher coefficient of determination). We also form the vector of response variables $\mathbf{Y}_{ij} \in \mathbb{R}^{N\times 1}$, whose $r$-th entry is $x^{(r)}_{ij}$ for a given phenotype $i \in \{\mathrm{b},\mathrm{r},\mathrm{s},\mathrm{k}\}$ at a given age $j \in \{1,\dots,N_\mathrm{a}\}$. Then, we consider the linear regression of phenotype on genotype

$$\mathbf{Y}_{ij} = \mathbf{X}\boldsymbol{\beta}_{ij} + \text{residual}, \tag{S39}$$

where the $l$-th entry of the vector $\boldsymbol{\beta}_{ij} \in \mathbb{R}^{(N_\mathrm{g}N_\mathrm{a}+1)\times 1}$ is the partial regression coefficient of the $i$-th phenotype at age $j$ on the $(l-1)$-st genotypic difference in $\mathrm{vec}(\boldsymbol{\delta}^{(r)})$ (for $l > 1$) and the intercept (for $l = 1$). The least-squares estimate of these coefficients is given by

$$\hat{\boldsymbol{\beta}}_{ij} = (\mathbf{X}^\intercal\mathbf{X})^{-1}(\mathbf{X}^\intercal\mathbf{Y}_{ij}). \tag{S40}$$

The replicas of the breeding values of phenotype $i$ at age $j$ are listed by the vector $\mathbf{X}\hat{\boldsymbol{\beta}}_{ij}$. Denoting by $\mathrm{var}[\mathbf{Y}_{ij}]$ the variance among the entries of $\mathbf{Y}_{ij}$, the heritability of the phenotype $i$ at age $j$ is thus given by

$$h^2_{ij} = \frac{\mathrm{var}[\mathbf{X}\hat{\boldsymbol{\beta}}_{ij}]}{\mathrm{var}[\mathbf{Y}_{ij}]}. \tag{S41}$$

With these definitions, heritabilities, contributions to phenotypic differences, and the regression effects ($\hat{\boldsymbol{\beta}}$) depend on the population sample (the former two via $\mathbf{X}$ and $\mathbf{Y}_{ij}$, the latter via $\mathbf{y}^{(r)}$ and $\boldsymbol{\alpha}^{(r)}$) but sensitivities (e.g., $\mathrm{d}x_{ia}/\mathrm{d}\mathbf{y}|_{\mathbf{m}=\bar{\mathbf{m}}}$), elasticities, and reactivities do not, so in this sense sensitivities, elasticities, and reactivities are individual-level properties. Mechanistic heritability may be defined in terms of mechanistic breeding values (i.e., mean value $\bar{x}_{ia}$ plus genetic contribution $C^{(r-q)}_{ia\mathbf{y}}$ to difference relative to conditions $q$ being the average; González-Forero, 2023, 2024b). However, mechanistic heritability can be greater than one because mechanistic breeding values may be correlated with the mechanistic residual, whereas breeding values are not correlated with the residual because of least squares (González-Forero, 2023, 2024b).

### S6 Code correction

The computer code of González-Forero (2024a) contains a typo, which has been corrected here. Specifically, note that Eq. (S35) is

$$\frac{\partial B_{\mathrm{rest,b}}(\mathbf{x}_a, \mathbf{y}_a, \bar{x}_{\mathrm{k}a}, a)}{\partial y_{ia}} = \frac{\partial x_{\mathrm{b},a+1}}{\partial y_{ia}} E_\mathrm{b}. \tag{S42}$$

In the computer code of González-Forero (2024a), this equation was implemented in the file `BrainFunctions.jl` in line 237 as

$$\partial\mathtt{Brestb}\partial\mathtt{y(i,a,x,y,par)} = \partial\mathtt{xj}\partial\mathtt{yia(1,i,a,x,y,par)} * \mathtt{par.E[1]}, \tag{S43}$$

where $\partial$`xj`$\partial$`yia(i,j,a,x,y,par)` is defined in line 216 of that file giving the direct effect of genotype $i$ at age $a$ on the phenotype $j$ at the next age. Thus, the implementation (S43) is incorrect as it exchanged the order of the first two arguments of $\partial$`xj`$\partial$`yia(i,j,a,x,y,par)`; that is, instead of implementing $\partial x_{\mathrm{b},a+1}/\partial y_{ia}$, the code implemented $\partial x_{i,a+1}/\partial y_{\mathrm{b}a}$. This means that rather than using the two bottom entries of the leftmost column of the matrix in Eq. (S36), the second and third entries of the top row of that matrix were used. This entails that this typo only exchanged $E_\mathrm{b}$ with $E_\mathrm{r}$ or $E_\mathrm{s}$ in the evaluation of $\partial x_{\mathrm{b},a+1}/\partial y_{ia}$ in (S42).

The correct implementation is instead

$$\partial\mathtt{Brestb}\partial\mathtt{y(i,a,x,y,par)} = \partial\mathtt{xj}\partial\mathtt{yia(i,1,a,x,y,par)} * \mathtt{par.E[1]}. \tag{S44}$$

This correction has now been implemented here.

After re-running all results, we find that this correction primarily affects the ontogenetic patterns but leaves essentially the same adult patterns. Specifically, González-Forero (2024a) found evolved ontogenetic patterns of brain and body size under the *H. sapiens* scenario that differed somewhat from, and were a poorer fit to observation than those found by the optimisation approach of González-Forero and Gardner (2018) that assumed evolutionary equilibrium and used optimal control. González-Forero (2024a) suggested that the differences could have been due to the discretisation of developmental time, since González-Forero and Gardner (2018) used continuous time. In the correction implemented here, the evolved ontogenetic patterns now match those obtained by González-Forero and Gardner (2018) via optimisation, so the discrepancies were due to the typo identified. A correction of González-Forero (2024a) will be submitted soon, which shows that the primary conclusions of González-Forero (2024a) are not affected by this correction.

### S7 Supplementary Figures

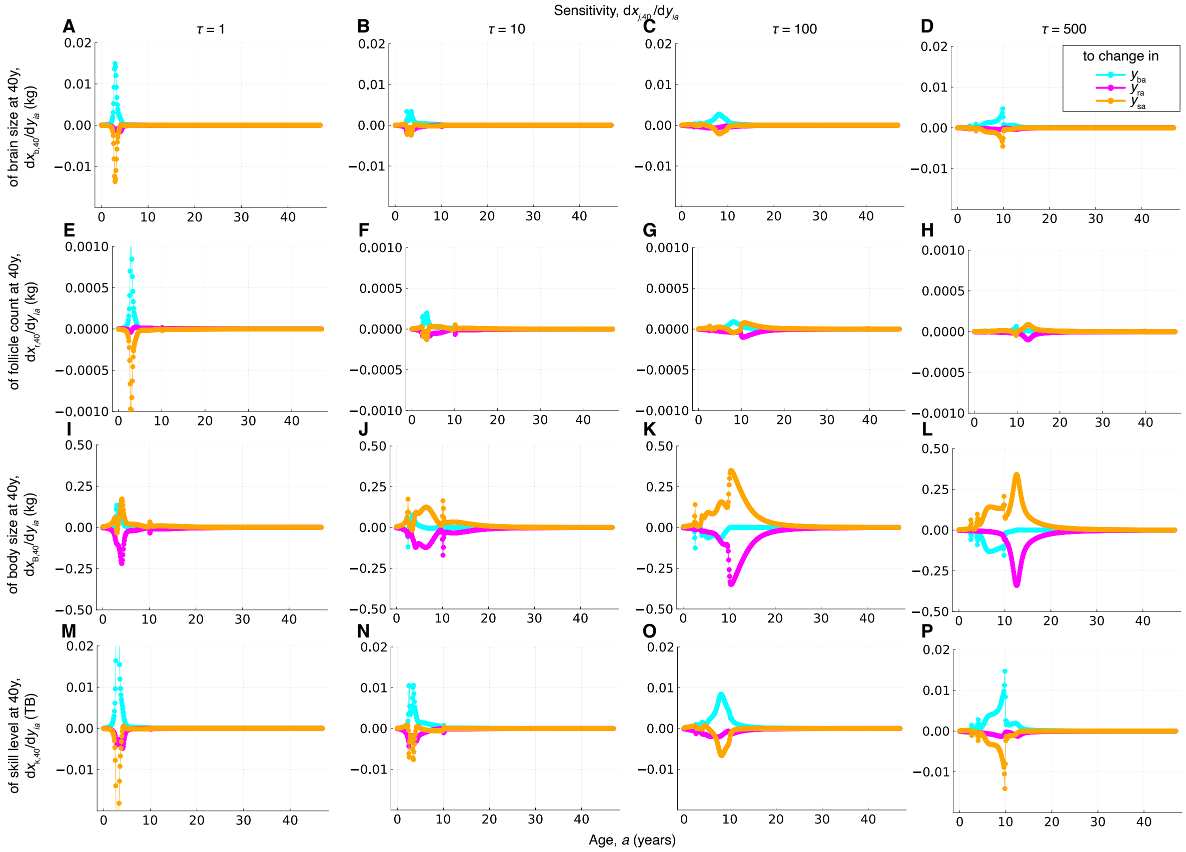


Figure S1: **Sensitivity of the phenotype at 40 years of age to genotypic change.** The skill level at 40 years of age is particularly sensitive to changes in allocation to growth of brain or soma at the age of brain growth arrest (**P** blue and orange). Body size at 40 years of age is particularly sensitive to changes in allocation to growth of soma or mature ovarian follicle count during childhood and specially at the age of menarche (**L** pink and orange).

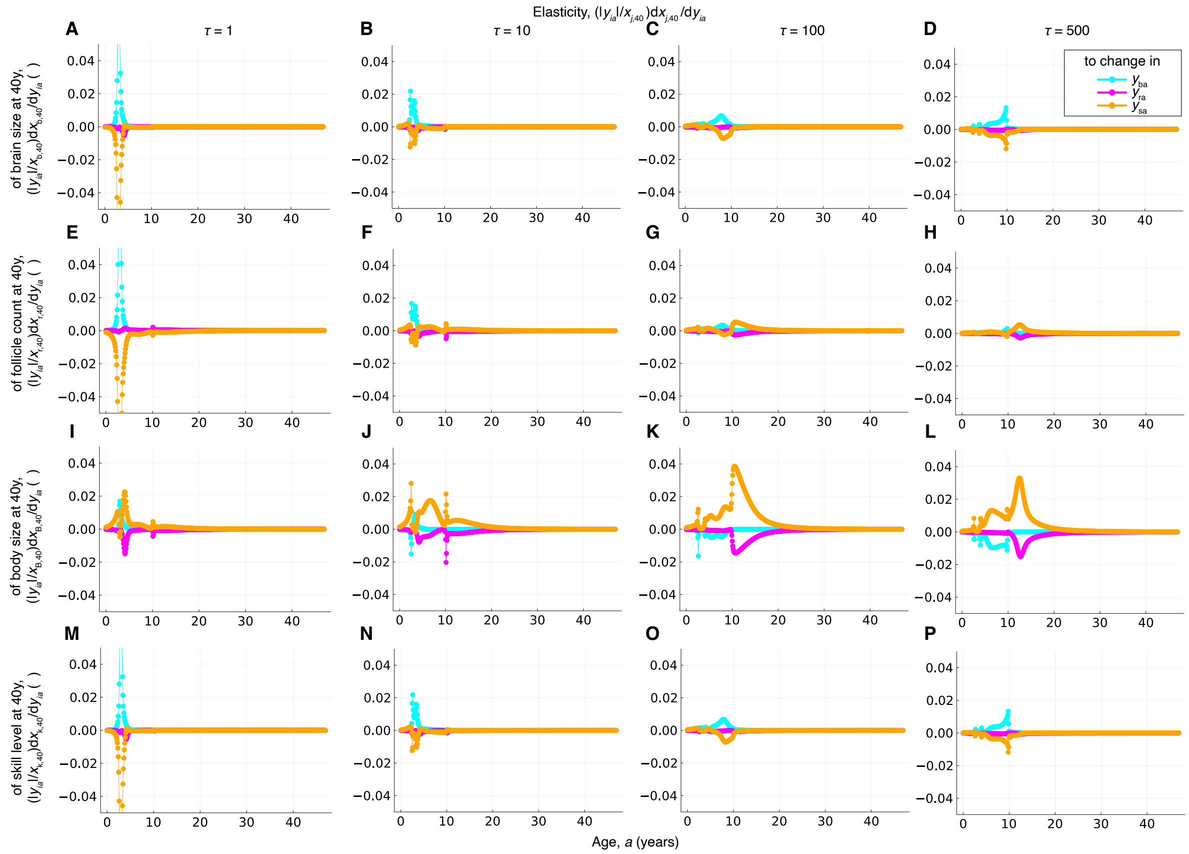


Figure S2: **Elasticity of the phenotype at 40 years of age to genotypic change.**

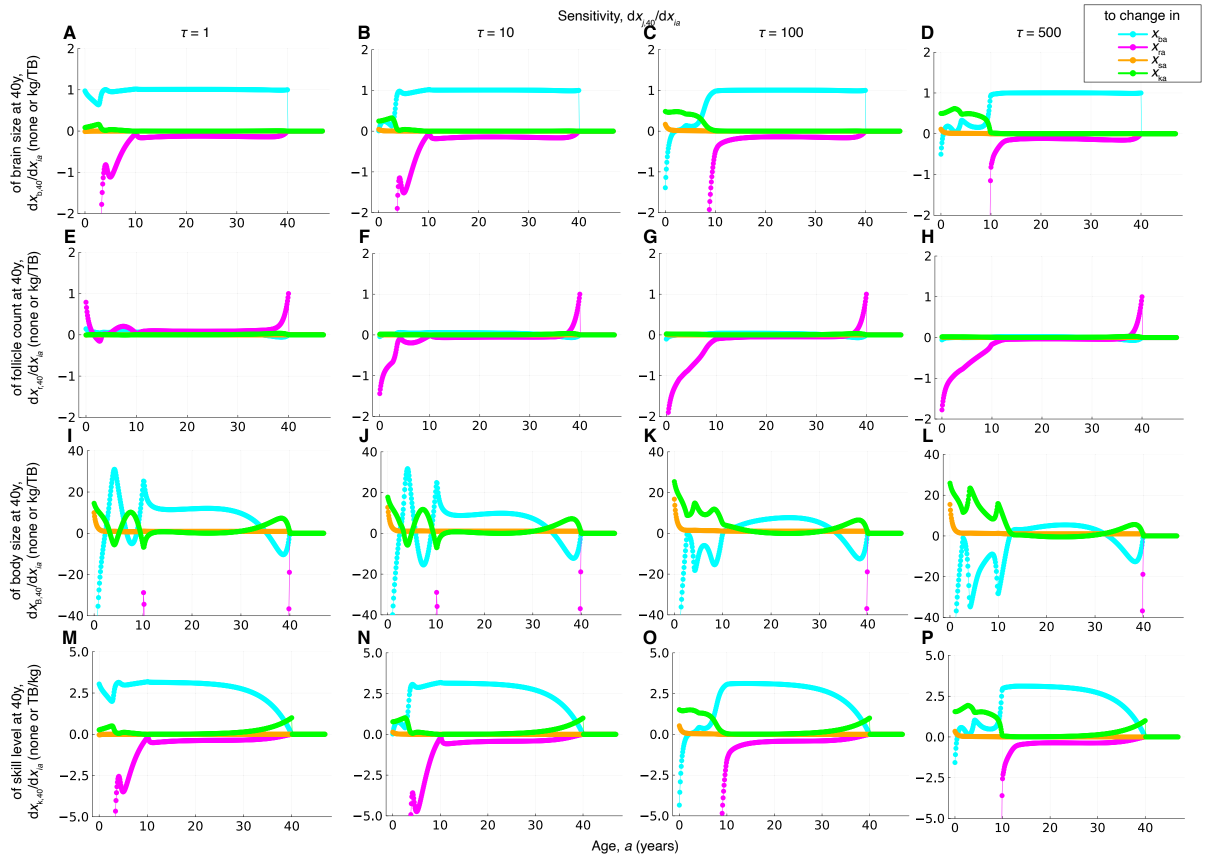


Figure S3: **Sensitivity of the phenotype at 40 years of age to phenotypic change.** All phenotypes at 40 years of age are extremely negatively sensitive to increases in mature ovarian follicle count, particularly during childhood (**D**, **H**, **L**, **P** pink).

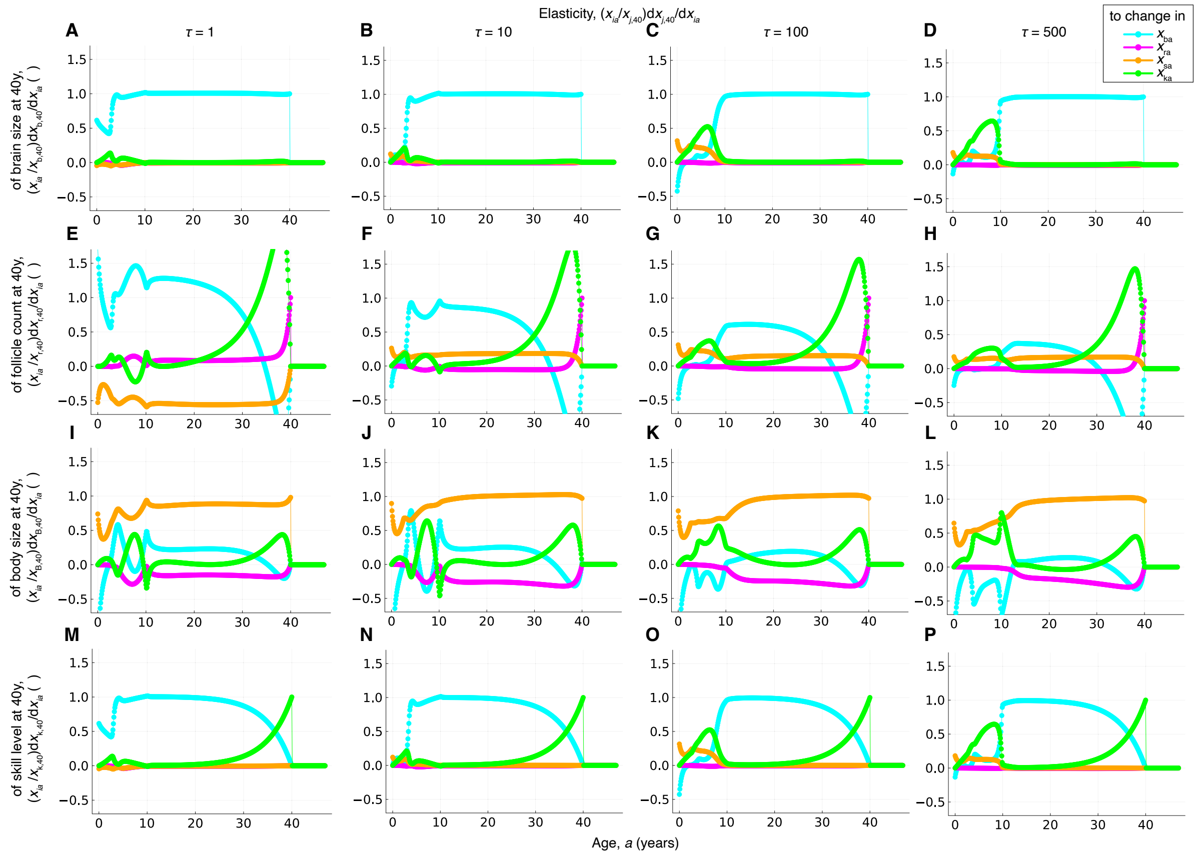


Figure S4: **Elasticity of the phenotype at 40 years of age to phenotypic change.**

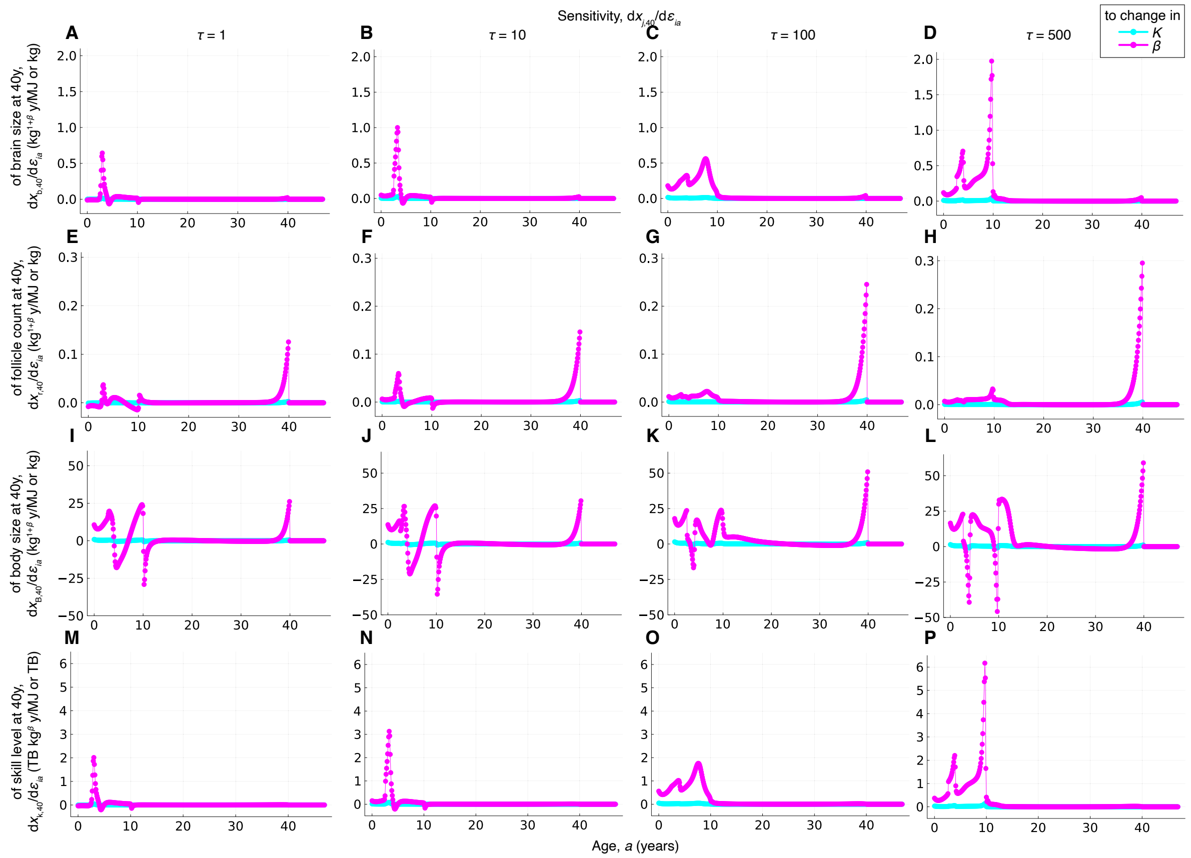


Figure S5: **Sensitivity of the phenotype at 40 years of age to change of Kleiber's law parameters.** Perturbing at particular ages the Kleiber's law exponent $\beta$ has large effects on the phenotypes at 40 years of age (**D**, **H**, **L**, **P** pink).

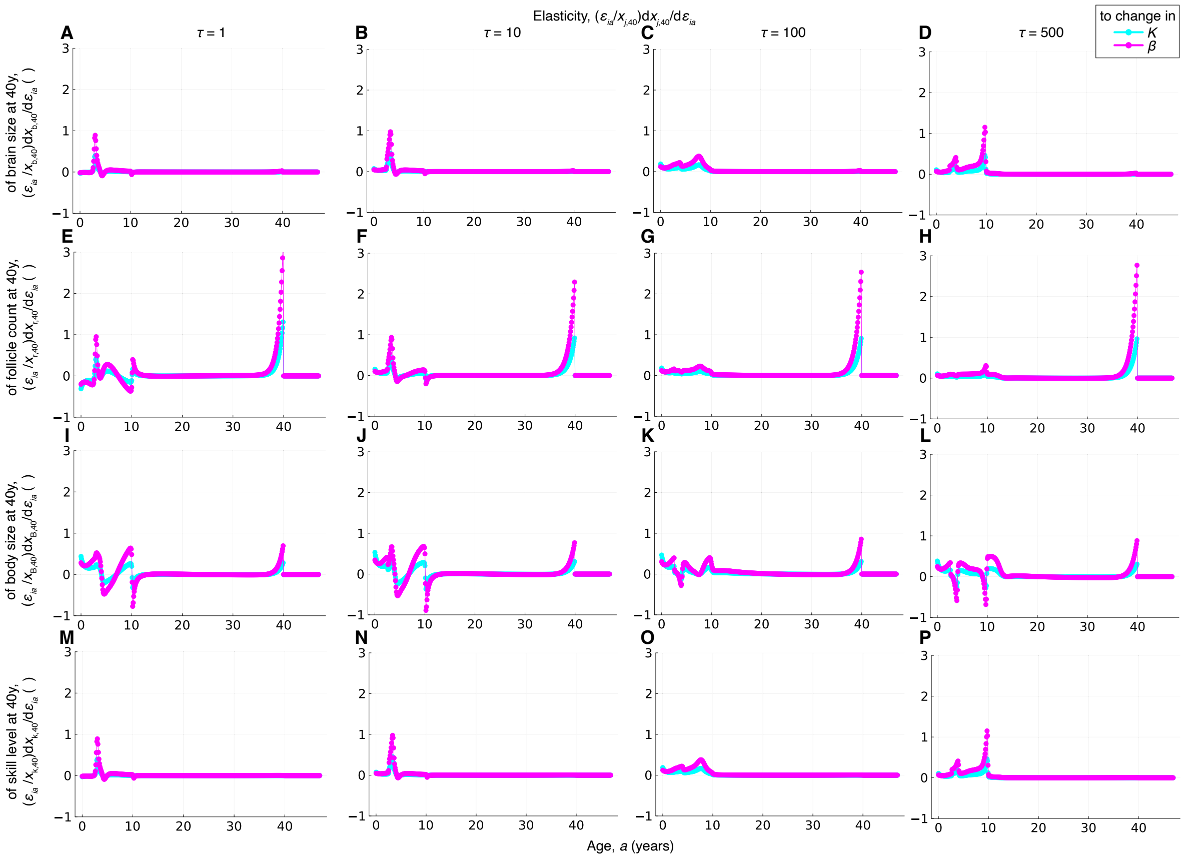


Figure S6: **Elasticity of the phenotype at 40 years of age to change of Kleiber's law parameters.**

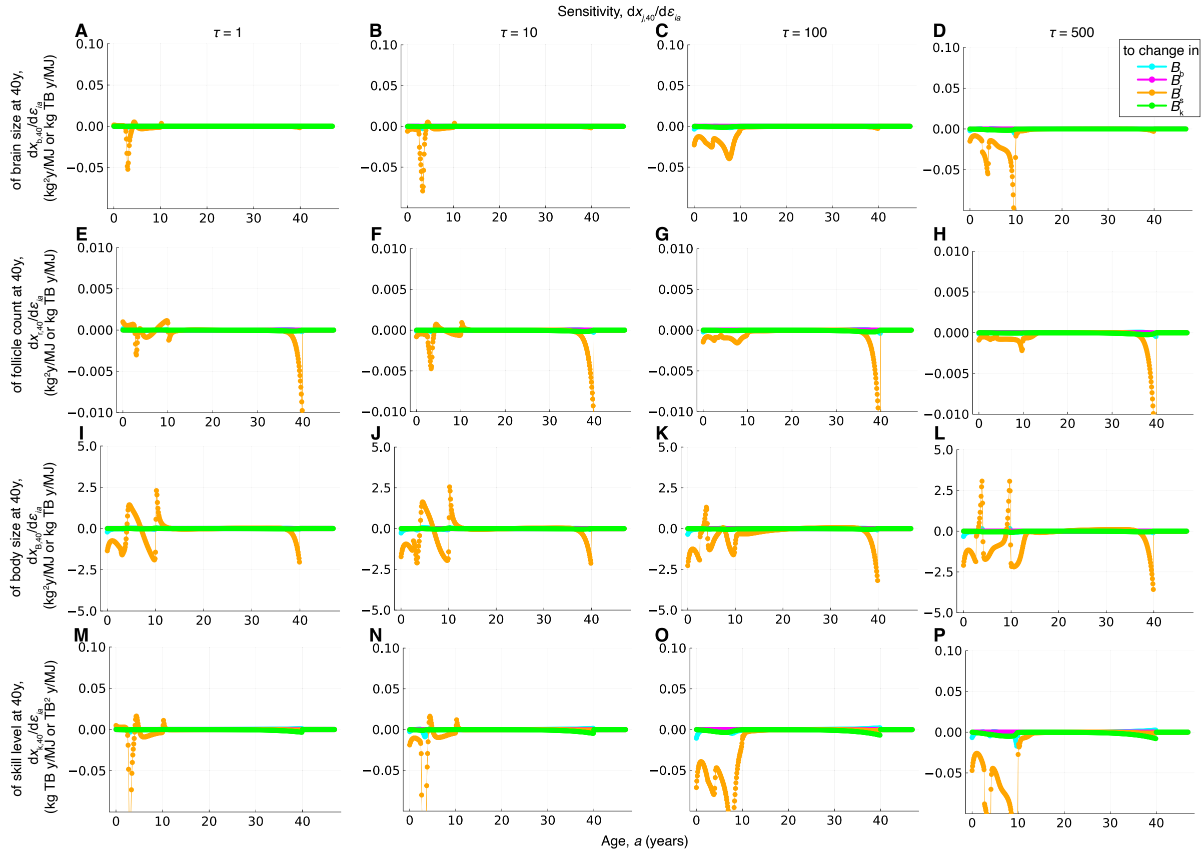


Figure S7: **Sensitivity of the phenotype at 40 years of age to change of metabolic costs of maintenance.** Perturbing the metabolic cost of maintenance of somatic tissue, particularly during childhood, has particularly strong effects on the phenotypes at 40 years of age (**D**, **L**, **P** orange).

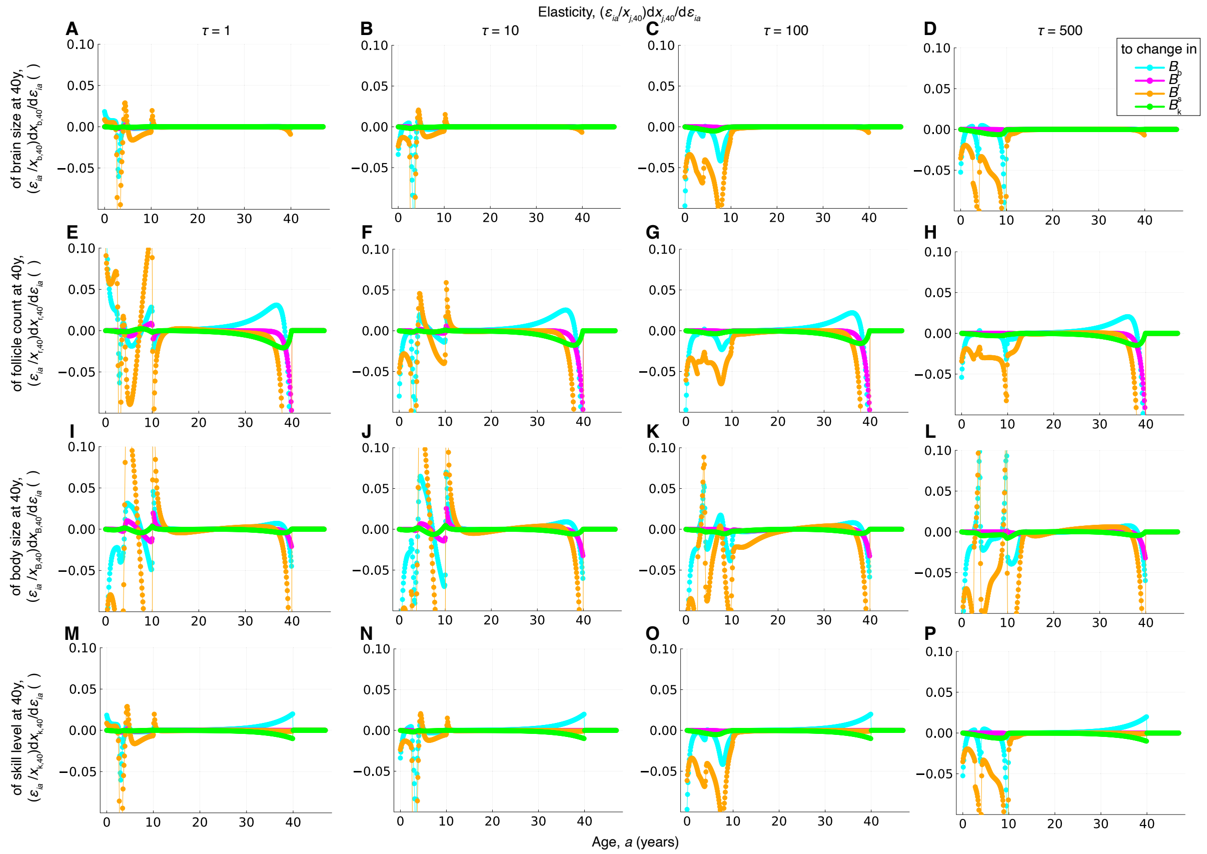


Figure S8: **Elasticity of the phenotype at 40 years of age to change of metabolic costs of maintenance.**

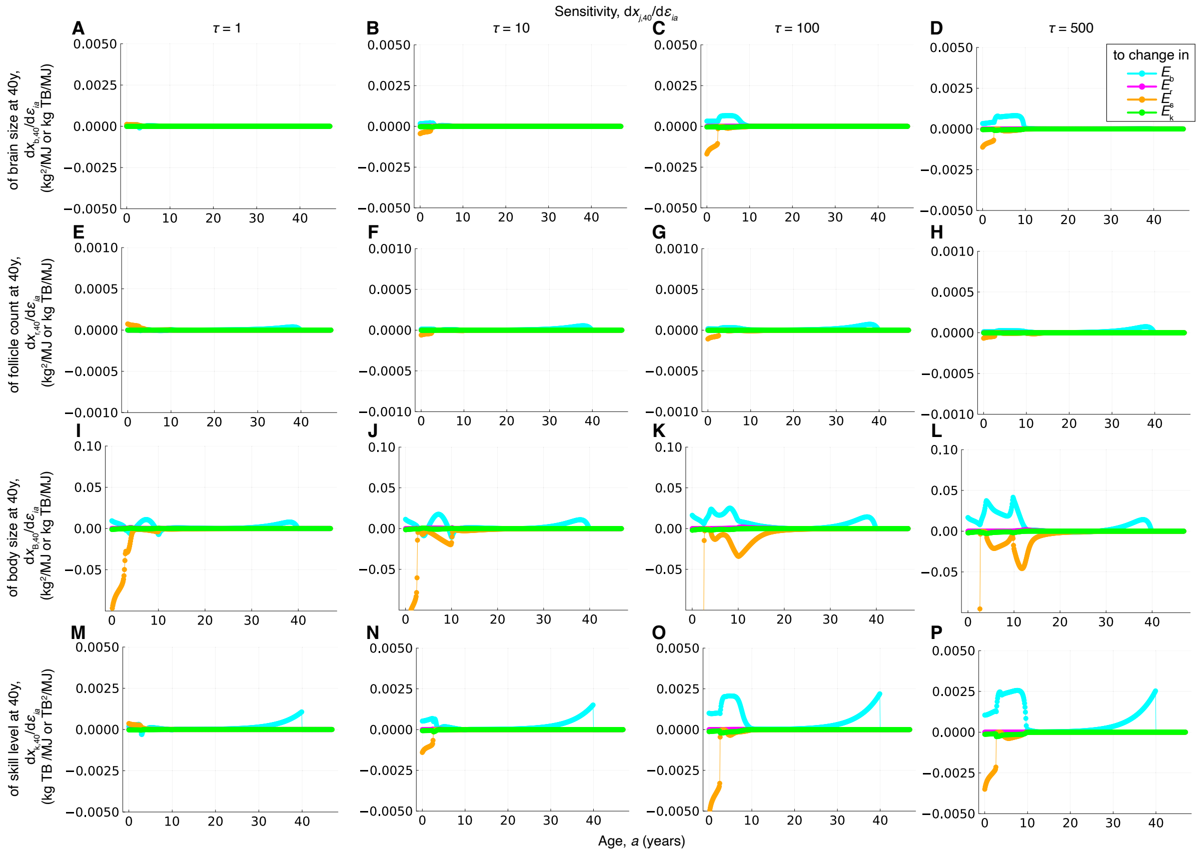


Figure S9: **Sensitivity of the phenotype at 40 years of age to change of metabolic costs of growth.** Increasing the metabolic costs of brain growth in childhood and near the evaluated age has a positive effect on the skill level at 40 years of age (**P** blue).

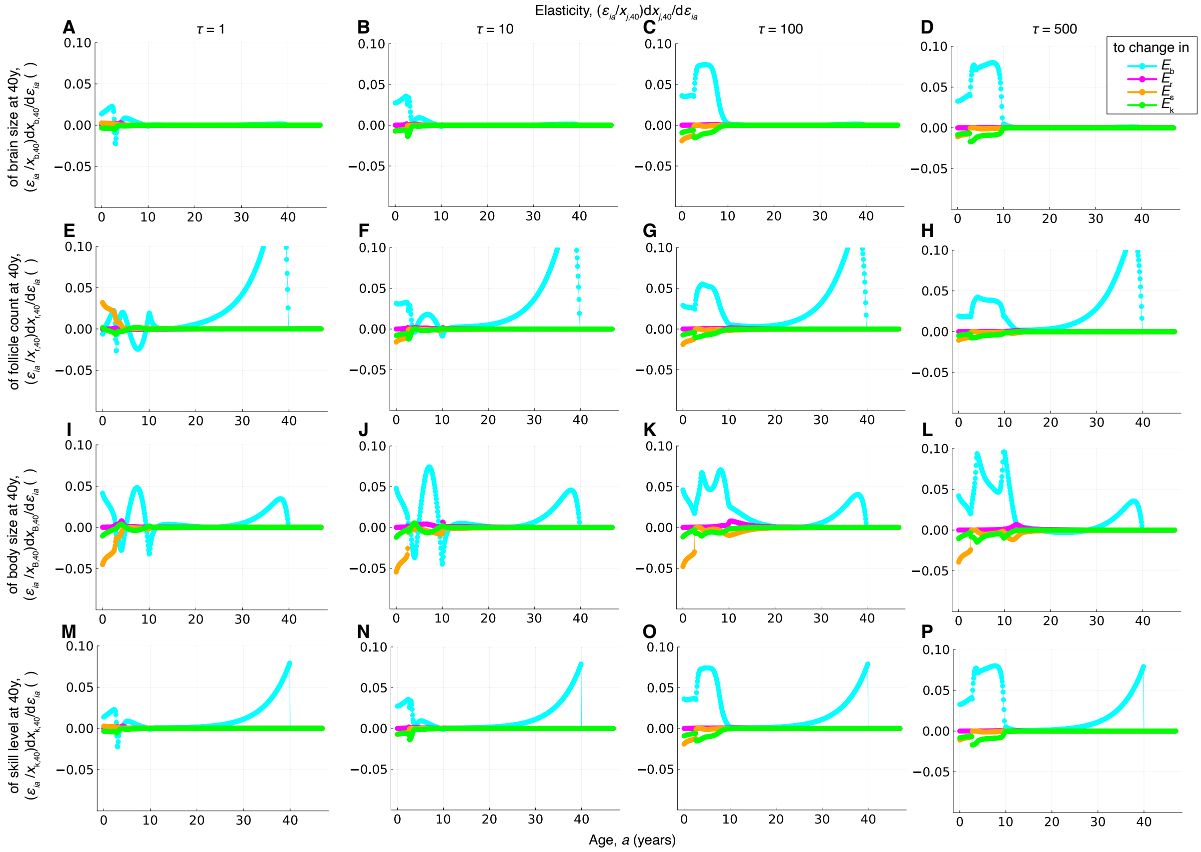


Figure S10: **Elasticity of the phenotype at 40 years of age to change of metabolic costs of growth.**

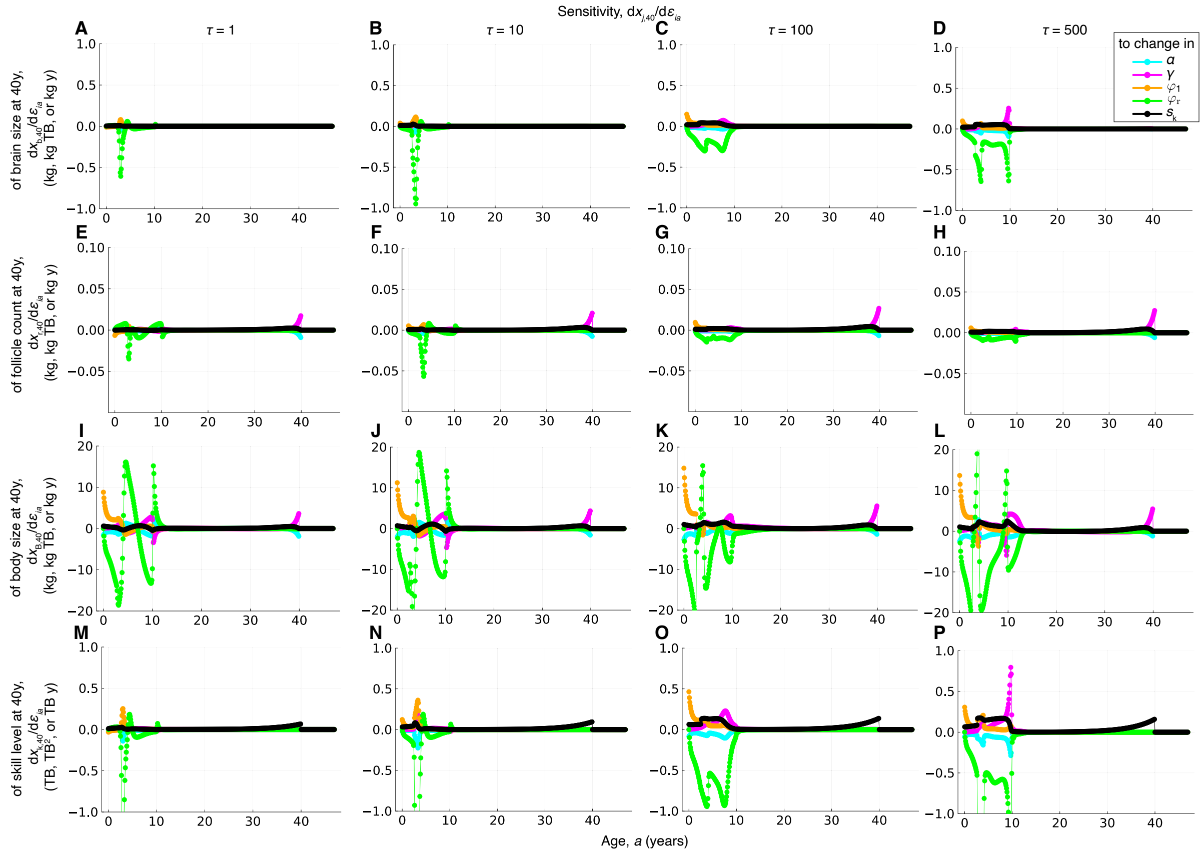


Figure S11: **Sensitivity of the phenotype at 40 years of age to change of skill parameters.** Increasing the rate of maternal care decline has particularly strong effects on the phenotypes at 40 years of age, typically negative but also positive for body size (**D**, **H**, **L**, **P** green).

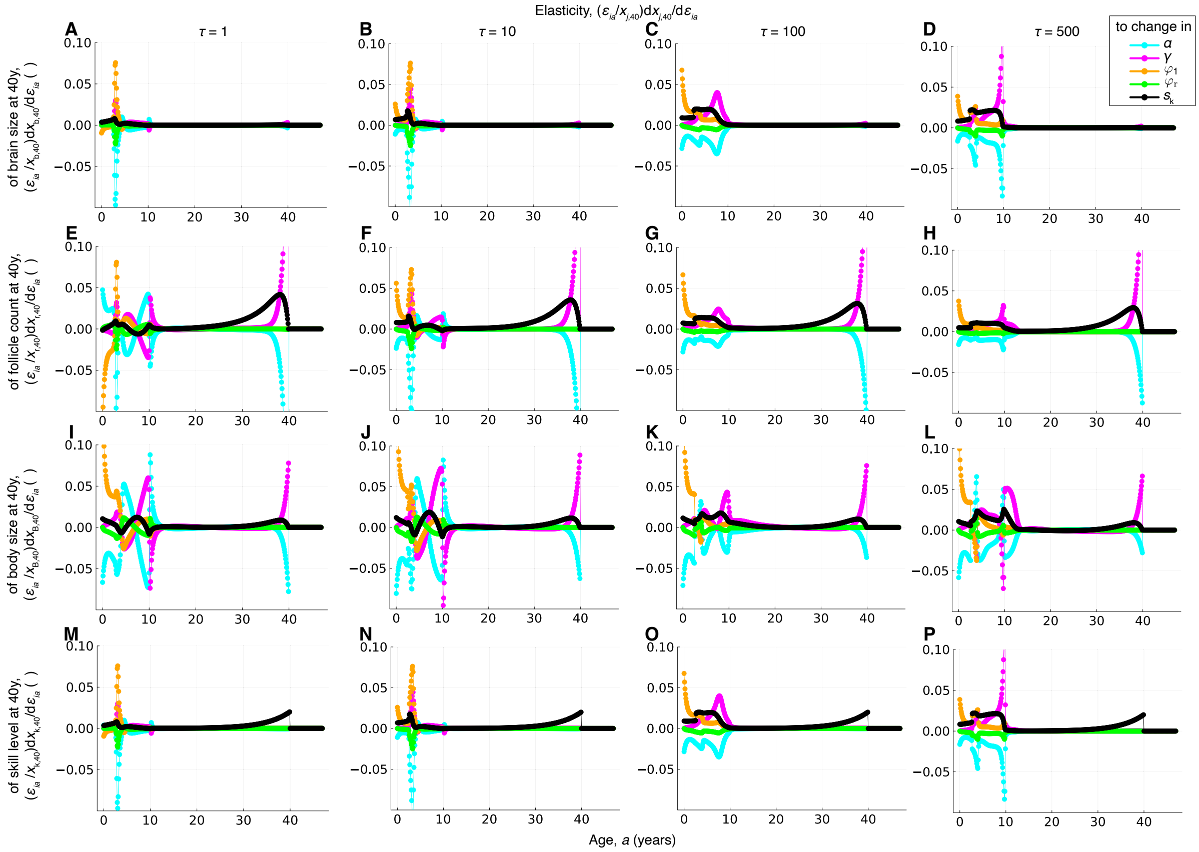


Figure S12: **Elasticity of the phenotype at 40 years of age to change of skill parameters.**

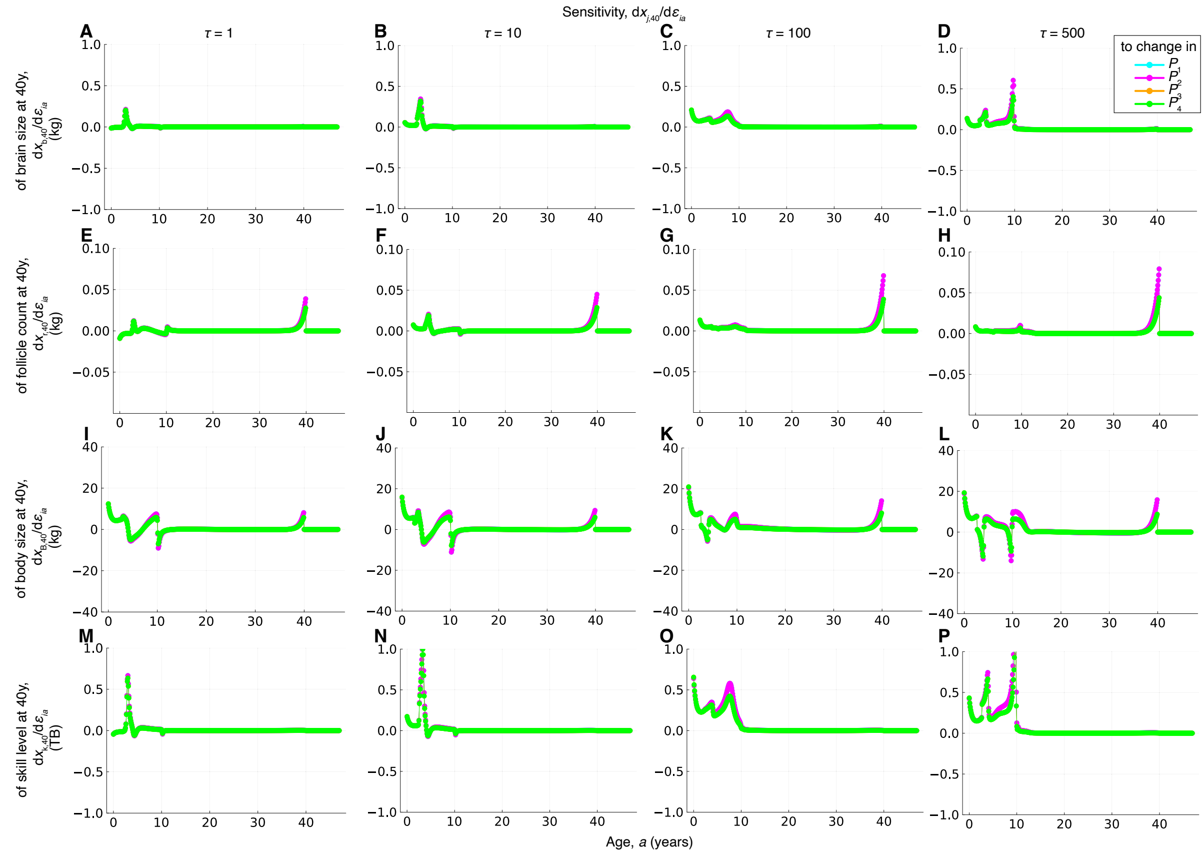


Figure S13: **Sensitivity of the phenotype at 40 years of age to change of challenge proportions.** The phenotypes at 40 years of age have a similar sensitivity to changes in any of the challenge proportions, although slightly more to cooperative challenges (pink). Adult skill level is positively affected by increases in the challenge proportions during childhood (**P**). In contrast, adult body size is negatively affected by increases in challenge proportions at the ages of brain growth onset and arrest (**L**).

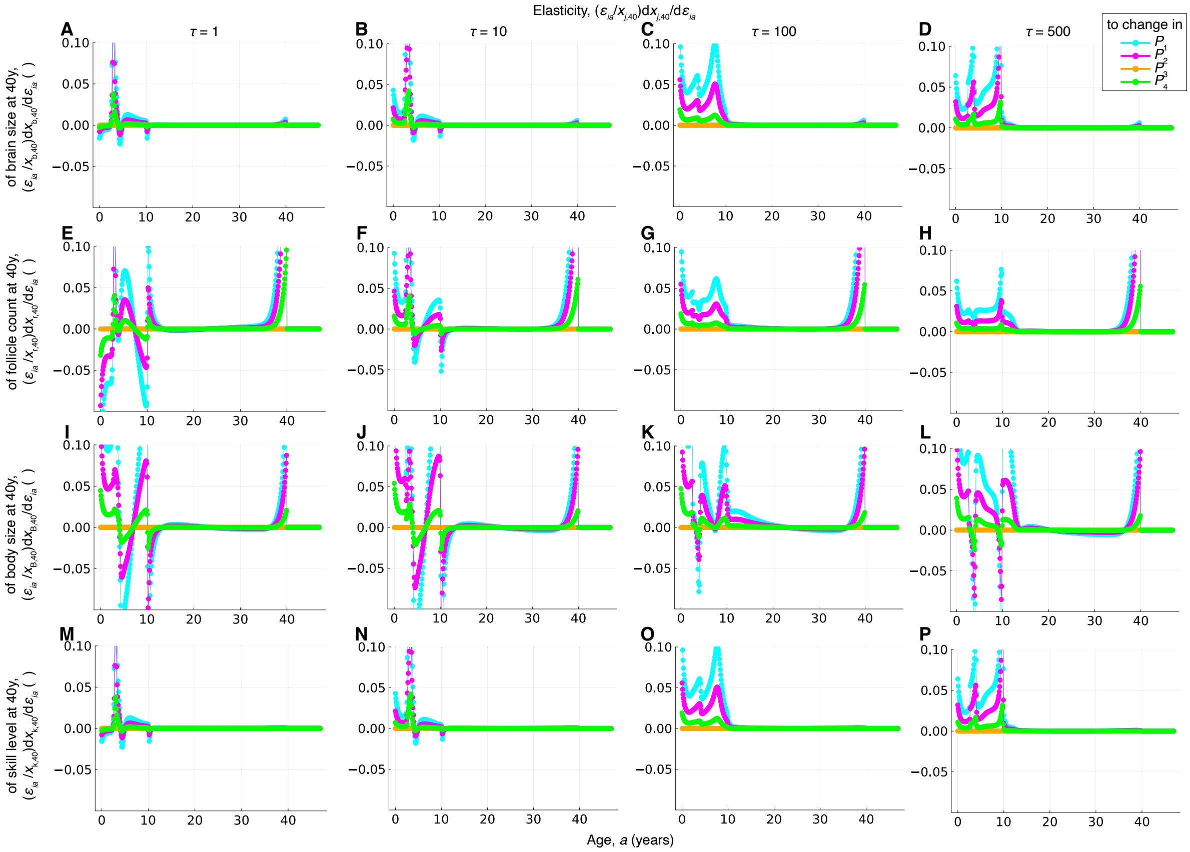


Figure S14: **Elasticity of the phenotype at 40 years of age to change of challenge proportions.**

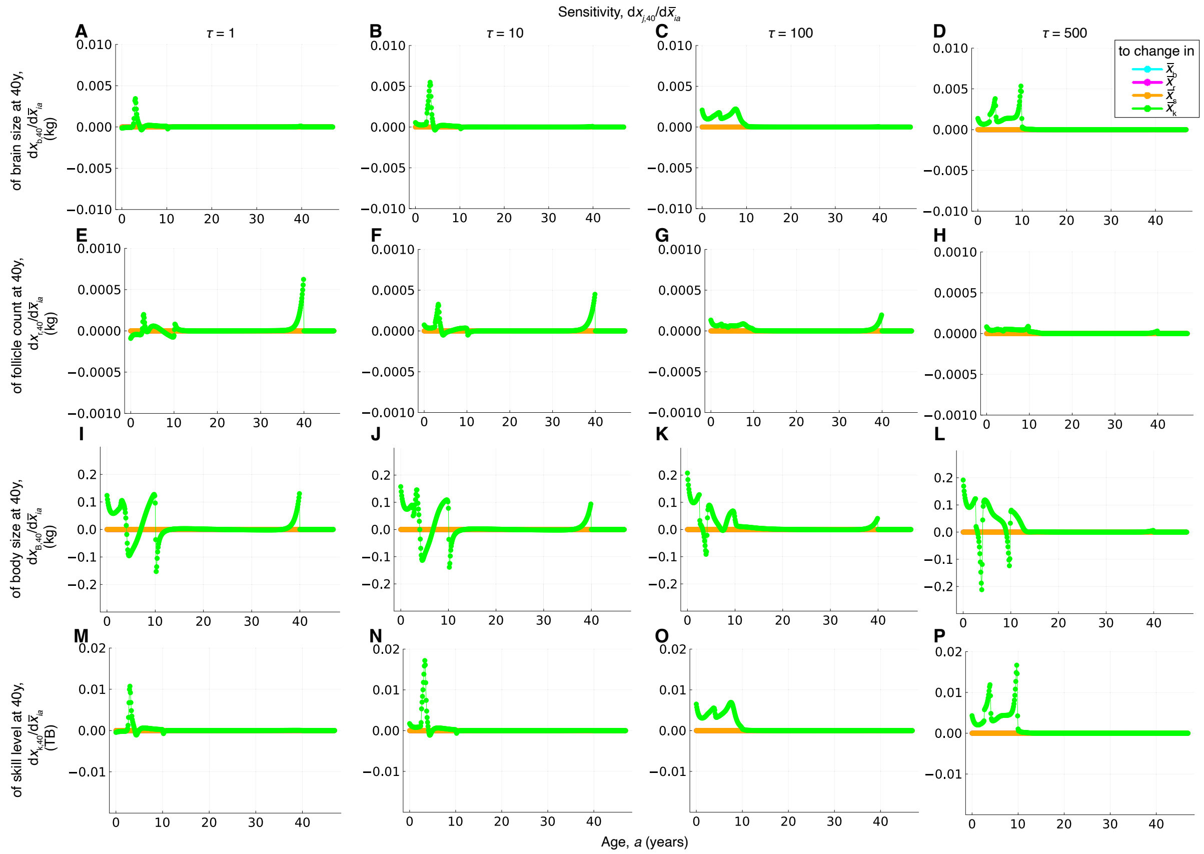


Figure S15: **Sensitivity of the phenotype at 40 years of age to change in social partners' phenotype.** The adult phenotype are insensitive to changes in the phenotypes of social partners, except their skill level (only green lines are non-zero). This is because only the social partner's skill level directly affects the development of a focal individual, but not other phenotypes of social partners. The adult phenotypes are particularly sensitive to changes in social partner's skill level during childhood (**D**, **H**, **L**, **P** green).

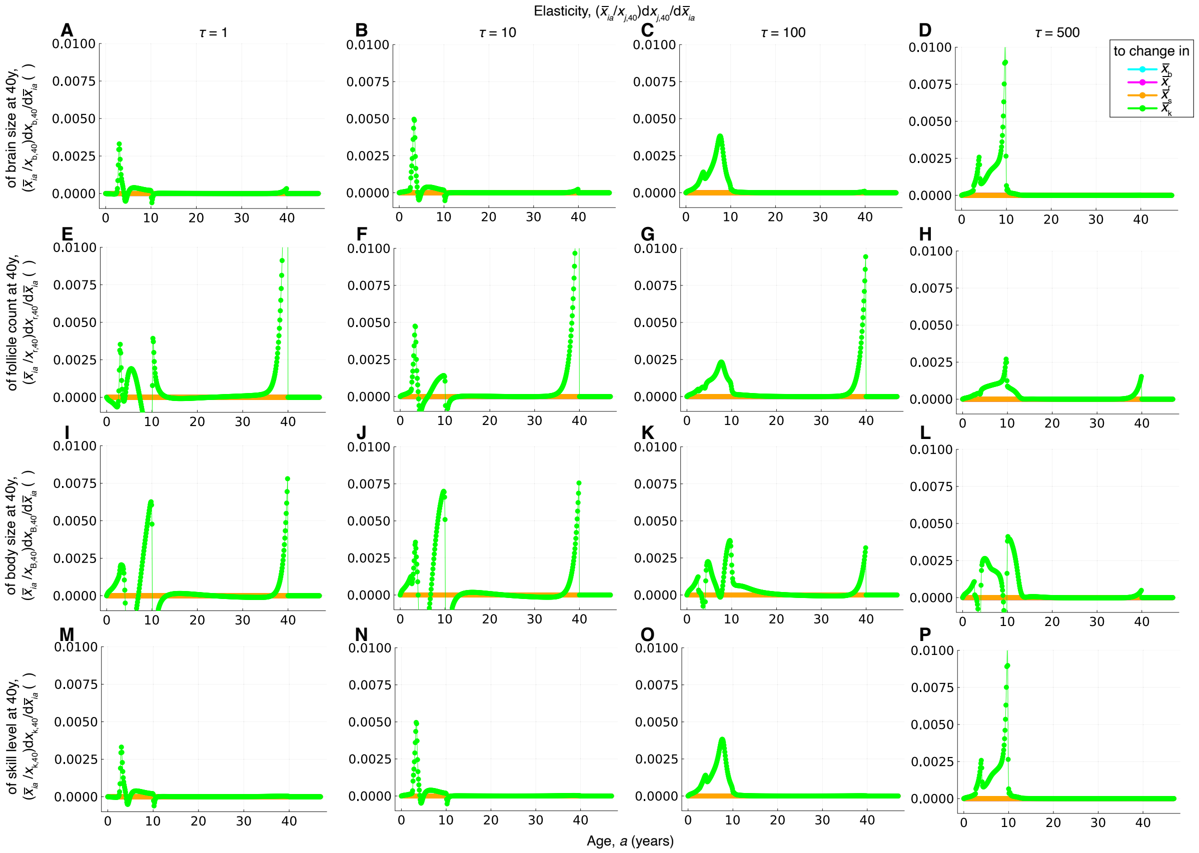


Figure S16: **Elasticity of the phenotype at 40 years of age to change in social partners' phenotype.**

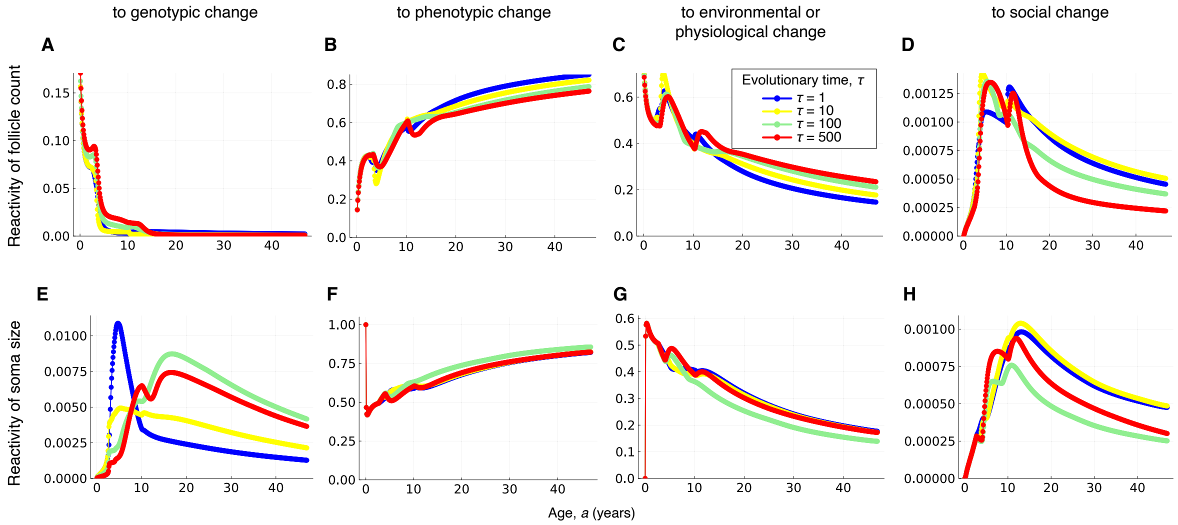


Figure S17: **Developmental reactivity of mature ovarian follicle count and somatic tissue size over the recovered hominin brain expansion.** Mature ovarian follicle count becomes non-reactive to genotypic change from about 14 years of age onwards (**A**).

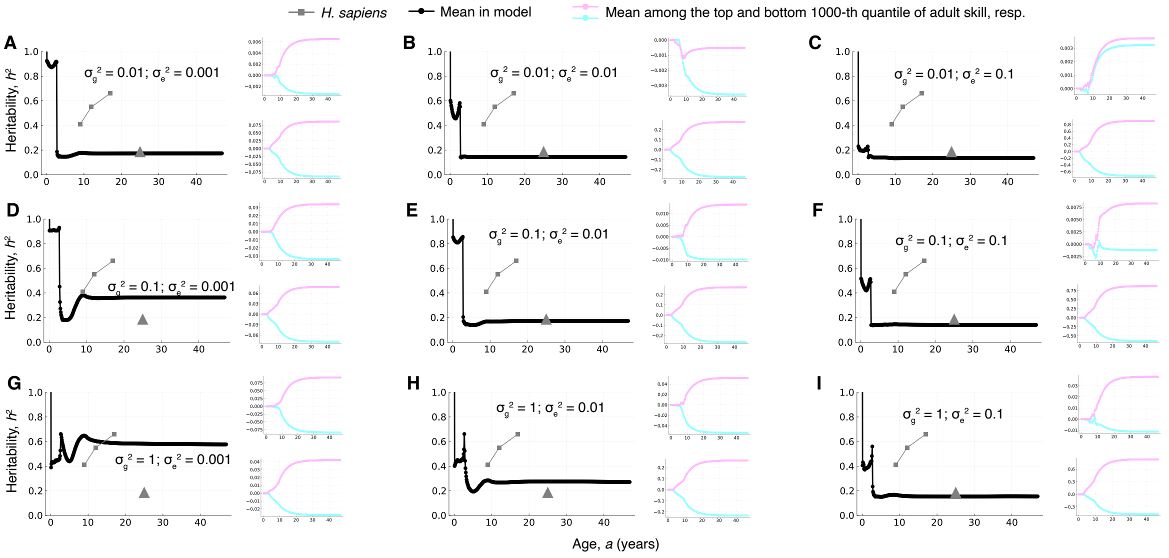


Figure S18: **Predicted heritability of skill level and first-order contributions to skill difference under various values of genotypic and environmental variation.** Each panel uses the indicated parameter values, where the large plot is that of heritability, the small top plot is that of the first-order contribution of genes, and the small bottom plot is that of the first-order contribution of the environment. Gray squares are the phenotypic-based heritability estimates of general cognitive ability of Haworth et al. (2010), for the ages estimated by that study. The gray triangle is the SNP heritability of cognitive performance estimated by Tan et al. (2024), placed here at 25 years of age but the age was not specified in that study. The SNP heritability estimated by Tan et al. (2024) is consistent with multiple values of genotypic and environmental variation, roughly when $\sigma_g^2 \leq 10\sigma_e^2$ (**A**, **B**, **C**, **E**, **F**, **I**). For the panels that recover the observed SNP heritability, the ratio of environmental-to-genetic contributions for adult top-skilled average individuals is roughly: (**A**) 0.075/0.006 = 12.5, (**B**) 0.2/(-0.01) = -20, (**C**) 0.8/0.003 = 267, (**E**) 0.2/0.01 = 20, (**F**) 0.75/0.0075 = 100, and (**I**) 0.6/0.03 = 20. This widely different ratios illustrate the limited ability of heritability to identify genetic contributions. Panel **E** corresponds to Fig. 5H,I,J of the main text, with some differences due to random variation. In contrast to heritability, the sensitivities, elasticities, and reactivities in Figs. 3 and 4 of the main text do not depend on genotypic and environmental variation $\sigma_g^2$ and $\sigma_e^2$ as they are individual-level properties rather than population-level ones.

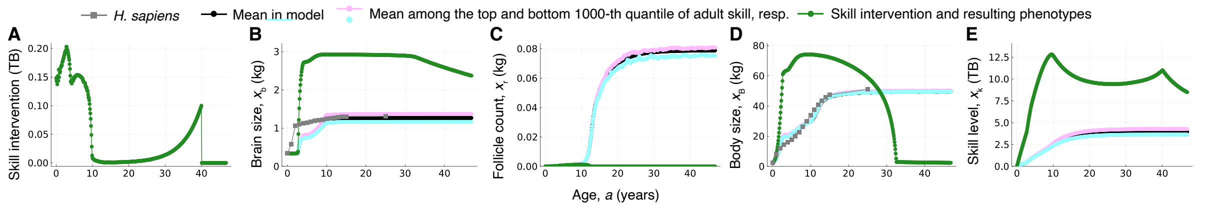


Figure S19: **Larger skill interventions disrupt development.** As in Fig. 6F-J, but the magnitude of skill intervention is ten times larger. (**A**) Skill level is intervened in the direction of steepest increase of adult skill level (i.e., changing skill level at the ages in the horizontal axis by 0.1 times the amount given by the sensitivity of adult skill level with respect to skill level at those ages). That is, a relatively large intervention in skill level is implemented. (**B-E**) Corresponding phenotypes developed. Although this skill intervention is weakened relative to that implemented for the environment (Fig. 6A-E), the bottom quantile individuals (green), who have genes detrimental to adult skill, develop (**B**) exceedingly large adult brain size, (**C**) infertility, (**D**) rapidly growing and then collapsing body size, and (**E**) exceedingly large skill level. We say "exceedingly" in the sense that such large phenotypes incur metabolic costs that are high enough to trigger the eventual collapse of the phenotypes over ontogeny as they make the growth metabolic rate negative (i.e., tissue maintenance demands more energy than the energy available). The model allows for these individuals as survival probability is constant, but if survival probability depended on these phenotypes these individuals would be inviable.
